# Phase-changing citrate macromolecule combats oxidative pancreatic islet damage, enables islet engraftment and function in the omentum

**DOI:** 10.1101/2023.10.25.564034

**Authors:** Jacqueline A. Burke, Yunxiao Zhu, Xiaomin Zhang, Peter D. Rios, Ira Joshi, Daisy Lopez, Hafsa Nasir, Sharon Roberts, Quetzalli Rodriguez, James McGarrigle, David Cook, Jose Oberholzer, Xunrong Luo, Guillermo A. Ameer

## Abstract

Clinical outcomes for total-pancreatectomy followed by intraportal islet autotransplantation (**TP-IAT**) to treat chronic pancreatitis (**CP**) patients are suboptimal due to the inflammatory state of the patient’s pancreas, oxidative tissue damage during the isolation process, and the harsh engraftment conditions in the liver’s vasculature, which include ischemia-reperfusion injury, and instant blood–mediated inflammatory reactions. We describe the use of the thermoresponsive, antioxidant macromolecule poly(polyethylene glycol citrate-co-N-isopropylacrylamide) (**PPCN**) to protect islet redox status and function *in vitro* and *in vivo* and to create a viable extrahepatic islet engraftment site in the abdomen. PPCN in aqueous media transitions from a liquid to an elastic hydrogel when exposed to body temperature via temperature-induced macromolecular self-assembly. Islets entrapped in the PPCN hydrogel and exposed to oxidative stress remain functional and support long-term euglycemia, in contrast to islets entrapped in a biologic scaffold (BS). When applied to the omentum of non-human primates (**NHPs**), PPCN is well-tolerated, safe, and mostly resorbed without fibrosis at 3 months post-implantation. To obtain autologous islets, a partial pancreatectomy was performed, followed by STZ administration to induce diabetes and destroy any remaining endogenous islets. Application of the autologous islets to the momentum using PPCN restored normoglycemia with minimal insulin requirements for over 100 days. These results support the use of PPCN as a scaffold for minimally invasive delivery of islets to the omentum of pancreatitis patients and highlight the importance of scaffold antioxidant properties as a new mechanism to protect islet function and maximize long-term autologous graft performance.

**One Sentence Summary:** Omentum islet transplantation using a thermoresponsive, antioxidative polymer supports autologous islet viability and function in nonhuman primates.

## INTRODUCTION

Chronic pancreatitis (**CP**) is a disease characterized by the progressive inflammation of the pancreatic acinar tissue (*1*). The etiology of this inflammation is varied and can include smoking, alcohol use, gallstones, and genetic variants (*1*). CP patients suffer from chronic abdominal pain, often leading to hospitalization, disability, and unemployment. As the inflammation persists, pancreatic tissue is destroyed as apoptotic acinar cells increase 10-fold and the mass of insulin-producing beta islet cells is reduced by 29% (*2*). There is no all-encompassing treatment for this disease. Management includes lifestyle modifications, optimization of pain medications, and nutrition. Surgical intervention is the last resort for relieving pain when pharmacological options no longer work (*1*).

CP patients unable to alleviate symptoms via less invasive measures often undergo total pancreatectomy (**TP**) followed by intrahepatic islet autologous transplantation (**IAT**), a procedure referred to as **TP-IAT** (*3–7*). Although islet transplantation has been improved through the development of standardized islet isolation procedures, long-term outcomes remain sub-optimal. Currently, islets are transplanted to the liver via intraportal infusion. Deleterious conditions such as liver thrombosis, instant blood-mediated inflammatory reactions, and oxidative stress are reported to contribute to significant damage to the transplanted islets (*4, 8, 9*). Approximately, 50 to 80% of the islets are destroyed via isolation and infusion (*10–12*). Given that CP patients have reduced islet masses under inflammatory conditions, this additional islet loss renders one-third of TP-IAT patients diabetic after the surgery (*13*). These findings highlight the importance of the transplant site and the need to support islet survival throughout the transplantation process. Therefore, there is a need for an alternate, extrahepatic islet transplant site and new islet delivery methods that provide a supportive microenvironment for islet function to improve the outcome of TP-IAT for CP patients.

The successful engraftment of islets at an extrahepatic site requires a microenvironment that can provide adequate vascularization and protection against oxidative stress conditions (*14–16*). Extrahepatic locations for islet transplantation have traditionally been limited to organ capsules, which have failed to be clinically adopted due to the invasive nature of the procedure (*17, 18*). The omentum has recently been investigated as a transplantation site in animals and humans due to its easy access, high vascularity, and potential to localize islets using preformed, solid scaffolds (*19*). However, the use of solid scaffolds makes it difficult to implement minimally invasive techniques, requires additional considerations for the placement and distribution of the islets within the scaffold, and can exacerbate inflammatory responses leading to fibrosis, limiting the widespread application of the procedure (*20, 21*). To address this issue, a recent clinical trial investigated the feasibility of using autologous plasma and recombinant human thrombin in a two-step endoscope-enabled procedure to deliver and secure allogeneic islets to the omentum for the treatment of type 1 diabetes (**T1D**) (*22*). However, allogeneic grafts at this site survived for less than 1 year, resulting in the termination of the trial (*23*). It is hypothesized that tissue-resident macrophages in the omentum prime alloreactive T cells to destroy allogeneic islets. Furthermore, the complexity of the procedure and the enhanced inflammatory status, and elevated oxidative and carbonyl stress conditions that are innate to autologous plasma from T1D patients likely contributed to variable outcomes (*23*). Given that alloreactive cell populations are not an issue for autologous transplantation, we hypothesize that the omentum could potentially be a viable transplantation site for CP patients if a scaffold can provide the proper microenvironment to protect and support the islets.

We hypothesized that body temperature-induced phase change of an antioxidant, water-soluble, degradable macromolecule would: 1) facilitate easy entrapment of islets and transportation to the omentum, 2) enable islet localization and engraftment on the target tissue upon delivery, and 3) significantly counter the negative effects of oxidative stress on islets after enduring pancreatitis, enzymatic isolation, and transplantation. Poly(polyethylene glycol citrate-co-N-isopropylacrylamide) (**PPCN**) is a citrate-containing macromolecule with a lower critical solution temperature that allows the transition from a liquid to a gel at physiological body temperature and has intrinsic antioxidant properties that mitigate oxidative damage to tissues (*24–28*). In this study, we report that PPCN protects mouse and human islets from oxidative stress-induced damage during the *in vitro* culture process as well as *in vivo* upon transplantation to the mouse fat pad. We also show that the application of PPCN to the omentum of nonhuman primates (**NHPs**) is safe and does not induce a deleterious foreign body response as the hydrogel is bioresorbable. Finally, we demonstrate that PPCN can support IAT in an NHP model for over 100 days, resulting in vascularized islets and minimal exogenous insulin requirements. As oxidative stress plays a pivotal role in the pathogenesis of pancreatitis, agents that can ameliorate oxidative stress should improve islet transplantation outcomes.

## RESULTS

### PPCN is anti-inflammatory, maintains islet viability, and supports insulin secretion in culture

PPCN was prepared via a two-step synthesis starting with a polycondensation reaction comprising citric acid, polyethylene glycol, and glycerol 1,3-diglycerol diacrylate (**GDD**) followed by free-radical polymerization with N-Isopropylacrylamide (**NIPAAm**) (**Fig. 1B**). PPCN dissolved in phosphate saline buffer (**PBS**) exhibits a reversible liquid to solid phase transition at the lower critical solution temperature (**LCST**) of 28°C, which is lower than that of the homopolymer poly(N-isopropylacrylamide) (**pNIPAAm**) (32°C) (**Fig. 1B-D**). Successful synthesis was confirmed using **^H^NMR**, **FTIR**, (**Fig. S1**), and rheology (**Fig. 1D**). At typical room temperatures, this LCST enables the easy addition of islets to the PPCN solution, their easy distribution to cell culture wells or delivery to target tissues in the body, and their entrapment at these locations via gelation at 37°C (**Fig. 1A,C,D**) (*29, 30*).

**Fig. 1.**
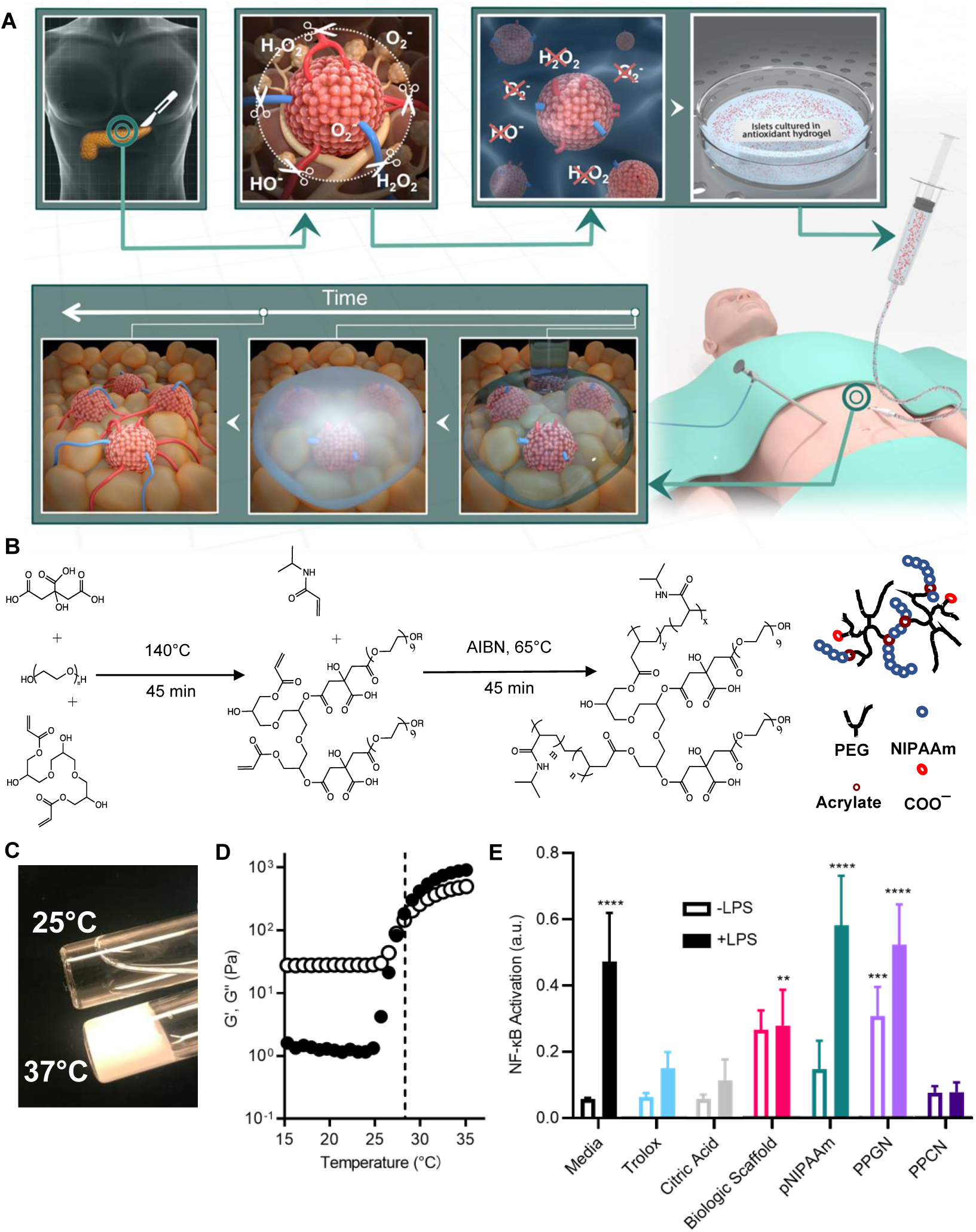
PPCN’s thermoresponsive antioxidant property facilitates islet preservation. (**A**) Schematic of PPCN-mediated islet protection against oxidative stress to preserve islet function throughout omentum transplantation. Top: Organ removal, islet tissue isolation, and transfer to a room temperature islet culture media containing the antioxidant thermoresponsive macromolecule PPCN that protects islets against oxidative damage during culture. Bottom: The thermoresponsive, phase-changing property of PPCN allows easy delivery of the islets in the liquid, localization through body temperature-induced gelation, and engraftment of islets into the omentum using laparoscopic surgery. (**B**) Schematic illustrating the synthesis of PPCN. (**C**) Digital photo showing the thermoresponsive transition of PPCN from a liquid (25°C) to a hydrogel (37°C). All samples were prepared in PBS at a concentration of 100 mg/mL and neutralized to pH 7.4. (**D**) Rheological determination of the lower critical solution temperature of the PPCN. (black marker – storage modulus G’; white marker – loss modulus G”). (**E**) Assessment of antioxidative properties for protection of RAW 264.7 macrophages cells against lipopolysaccharide (LPS)-induced NF-kB activation via Quanti-Blue cell-based assay. All data are presented as mean Nf-κB activation (a.u.) ± SD with *p<0.05; ** p<0.01; *** p<0.001; **** p<0.0001 relative to PPCN. Statistical significance was determined by two-way ANOVA with Tukey’s multiple comparisons test. (n = 5).

The anti-inflammatory properties of PPCN were confirmed *in vitro* as per the inhibition of NF-kB activation in the RAW-blue cell line (**Fig. 1E**). RAW-blue cells are engineered RAW264.7 macrophages that have been used to evaluate the intracellular antioxidant response due to lipopolysaccharide (**LPS**)-induced NF-kB expression (*31, 32*). Antioxidants are known to suppress NF-κB activation as well as the subsequent transcription of inflammation-related genes (*22, 33, 34*). Cells exposed to LPS in the presence of the antioxidant Trolox, an analog of vitamin E, reduced NF-kB expression by 68% relative to cells exposed to LPS in cell culture media. PPCN exhibited an 84% reduction in NF-kB expression (**Fig. 1E**). Cells exposed to a hydrogel formed from BS, a clinically used hydrogel for islet transplantation to the omentum in humans (*22*), effected a 41% inhibition of NF-kB expression, respectively (**Fig. 1E**). A non-antioxidant version of PPCN made with glutaric acid, instead of citric acid (**Fig. S2**), referred to as poly(polyethylene glycol glutarate-co-N-isopropylacrylamide) (**PP<u>G</u>N**) did not elicit the protective effects of PPCN on cells. Of note, we have previously shown that PPCN confers antioxidative properties due to the combination of citric acid and diols (*26, 35–37*) These intrinsic antioxidant properties include free-radical scavenging, iron chelation, and lipid peroxidation inhibition. The replacement of citric acid with glutaric acid elimates these antioxidative properties.

To assess the impact of PPCN on islet function, the viability and insulin secretion function of mouse (**Fig. 2A,C,E**) and human (**Fig. 2B,D,F**) islets in standard suspension culture, and after entrapment in BS, PPCN, or pNIPAAm were evaluated *in vitro*. Freshly isolated islets were either cultured on tissue culture plastic or mixed with autologous BS, PPCN, or pNIPAAm solutions at room temperature. Islets were successfully entrapped in each hydrogel by adding thrombin to the BS or incubating the islet-PPCN or islet-pNIPAAm mixtures at 37°C. The viability of the entrapped islets was monitored over time with the resazurin assay (**Fig. 2**). For both mouse (**Fig. 2A**) and human (**Fig. 2B**) islets, viability was maintained at 48 hours of *in vitro* culture except for the pNIPAAm treated group. Islets from both species entrapped in pNIPAAm experienced a viability loss of 40% at 24 hours and 60% at 48 hours (**Fig. 2A,B**).

**Fig. 2.**
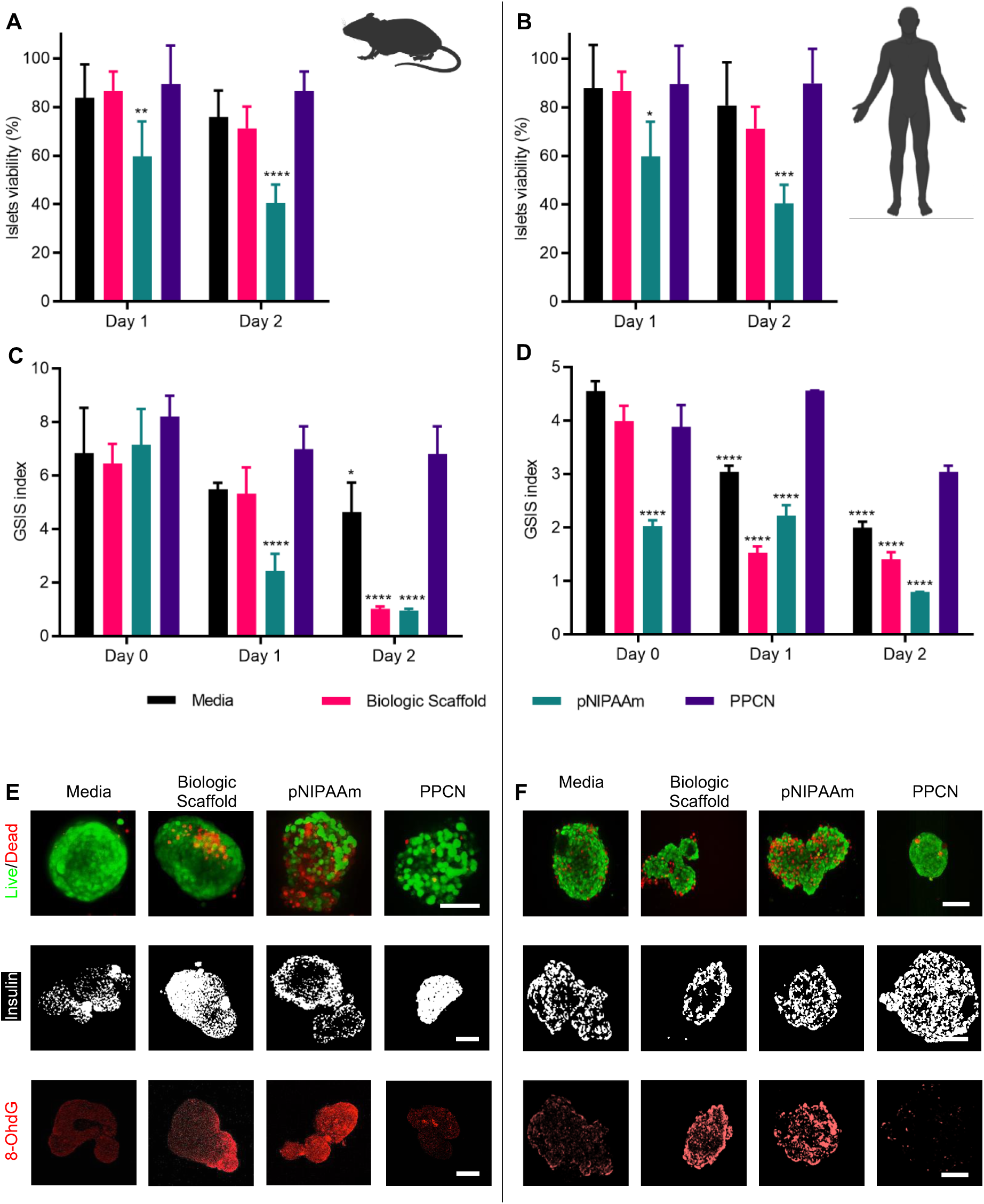
PPCN facilitates the preservation of both mouse and human islet viability and insulin secretion function *in vitro*. (**A,B**) Mouse and human islet viability as measured fluorescently by resazurin reduction after prolonged culture under various conditions. (**C,D**) Corresponding glucose-stimulated insulin secretion (GSIS) index of the cultured islets. All data are presented as mean ± SD with *p<0.05; ** p<0.01; *** p<0.001; **** p<0.0001 relative to PPCN. Statistical significance was determined by two-way ANOVA with Tukey’s multiple comparisons test. (n = 5). (**E,F**) Live/dead and immunostaining of mouse islets for insulin and 8-OHdG after 2 days of *ex vivo* culture under various conditions. (scale bar: 100 µm).

Islet functionality was measured by insulin secretion in response to glucose stimulation using the *in vitro* glucose stimulation/insulin secretion (**GSIS**) test, which reports a stimulation index (the GSIS index) (**Fig. 2C,D**). A GSIS index of 1 or below indicates the complete loss of glucose response (*38*). During *in vitro* culture, at day 0, mouse islets in all four groups have an average GSIS index of 7.15±1.23 (**Fig. 2C**). For mouse islets, GSIS index of the pNIPAAm group drops by 65% at 24 hours of culture, which is consistent with the loss of islet viability observed in this group (**Fig. 2C**). A relatively smaller drop in the GSIS index was measured for the other three groups (26% for suspension culture, 17.5% for BS, and 14.8% for PPCN) (**Fig. 2C**). Islets cultured in BS for 48 hours also lost their glucose responsiveness as their GSIS index dropped from 6.45±0.72 to 1.01±0.09 (**Fig. 2C**). Partial disassembly of the BS-entrapped islets also observed after 2 days in culture. Within 24 to 48 hours of culture, a 25% and 7.3% decrease in GSIS index was observed for islets in suspension culture and islets in PPCN, respectively, confirming the protective role of PPCN (**Fig. 2C**). Similar results were observed for human islets (**Fig. 2B,D**).

Live/dead staining of the mouse and human islets after 24 hours of culture confirmed the presence of many dead cells in the pNIPAAm-treated group whereas islets in suspension culture, in BS, or in PPCN exhibited similarly high viabilities (**Fig. 2E,F**). Intracellular insulin staining also revealed that murine islets cultured in pNIPAAm, and human islets cultured in BS showed signs of an insulin-deficient core, whereas islets cultured in PPCN showed uniform insulin expression across the entire islet structure. To assess whether the protective effects of PPCN may be due to its antioxidant properties, we probed for the nuclear DNA oxidation marker 8-oxo-2’-deoxyguanosine (**8-OHdG**) on both mouse and human islets after 48 hours in culture. Staining for 8-OHdG revealed significantly more oxidized residues in islets with lower function as measured by the GSIS index (**Fig. 2C-F**). In conclusion, culturing islets in PPCN enhances the preservation of sensitivity to glucose and associated insulin secretion function, likely by minimizing oxidation.

### PPCN protects islets against induced oxidative stress, thereby preserving function during culture

To further understand whether the antioxidant property of PPCN would preserve islets viability and function, redox-sensitive islets were created by expressing the transgene for the redox-sensitive green fluorescent protein (**roGFP**) gene in the cytosol of freshly harvested islets via a lentiviral vector. The incorporation of one disulfide bond between cysteine A147 and A204 in the protein structure of roGFP enables its use as a redox reporter that can indicate the oxidation status through the quantification of the relative fluorescence intensity at two excitation wavelengths 405 and 488 nm (**Fig. S3A,B**) (*39, 40*). Under normal reduced conditions, the protein is excited at 488 nm; however, upon exposure to an oxidizing environment, the disulfide bond formation leads to a shift in the excitation wavelength that peaks at 405 nm. This shift provides a signal difference that can be used to monitor and quantify the islet’s redox status within scaffolds by measuring the ratio between the fluorescence intensity emission after excitation at 405 or 488 nm. Transgene expression of roGFP in the cytosol did not affect the normal viability and insulin secretion function of both human and mouse islets (**Fig. S3C-F**).

To mimic oxidative damage to islets associated with CP *in vitro*, 10 μM hydrogen peroxide (**H_2_O_2_**) was added to each of the islet culture environments described in the aforementioned paragraph. This H_2_O_2_ concentration was chosen to mimic physiological H_2_O_2_ concentrations produced by acinar cells in experimental models of pancreatitis (*41*). Confocal microscopy imaging was used to monitor the progression of oxidative damage in both mouse and human islets. (**Fig. 3A,B**) At time 0, before the introduction of the H_2_O_2_, a dominant signal at 488 nm (reduced form shown as green) was observed in islets from all four groups with a baseline average oxidation percentage of 13% for mouse islets and 10% for human islets (**Fig. 3A-D**). Five minutes after introducing H_2_O_2_, an increase in oxidation of 27.0%, 23.1%, 20.6 %, and 14.6% was measured for mouse islets in the control media suspension culture, BS, pNIPAAm, and PPCN, respectively. At 30 minutes, the oxidation significantly increased for islets cultured in media suspension (36.5%), BS (30.4%), and pNIPAAm (30.4%), whereas oxidation of islets entrapped in PPCN increased to 19.7%. At 12 hours, 44% oxidation was observed in the PPCN group, whereas 80% oxidation was observed in the remaining groups. A similar trend was also observed for human islets (**Fig. 3B,D**); however, human islets appeared more resistant to oxidation overall.

**Fig. 3.**
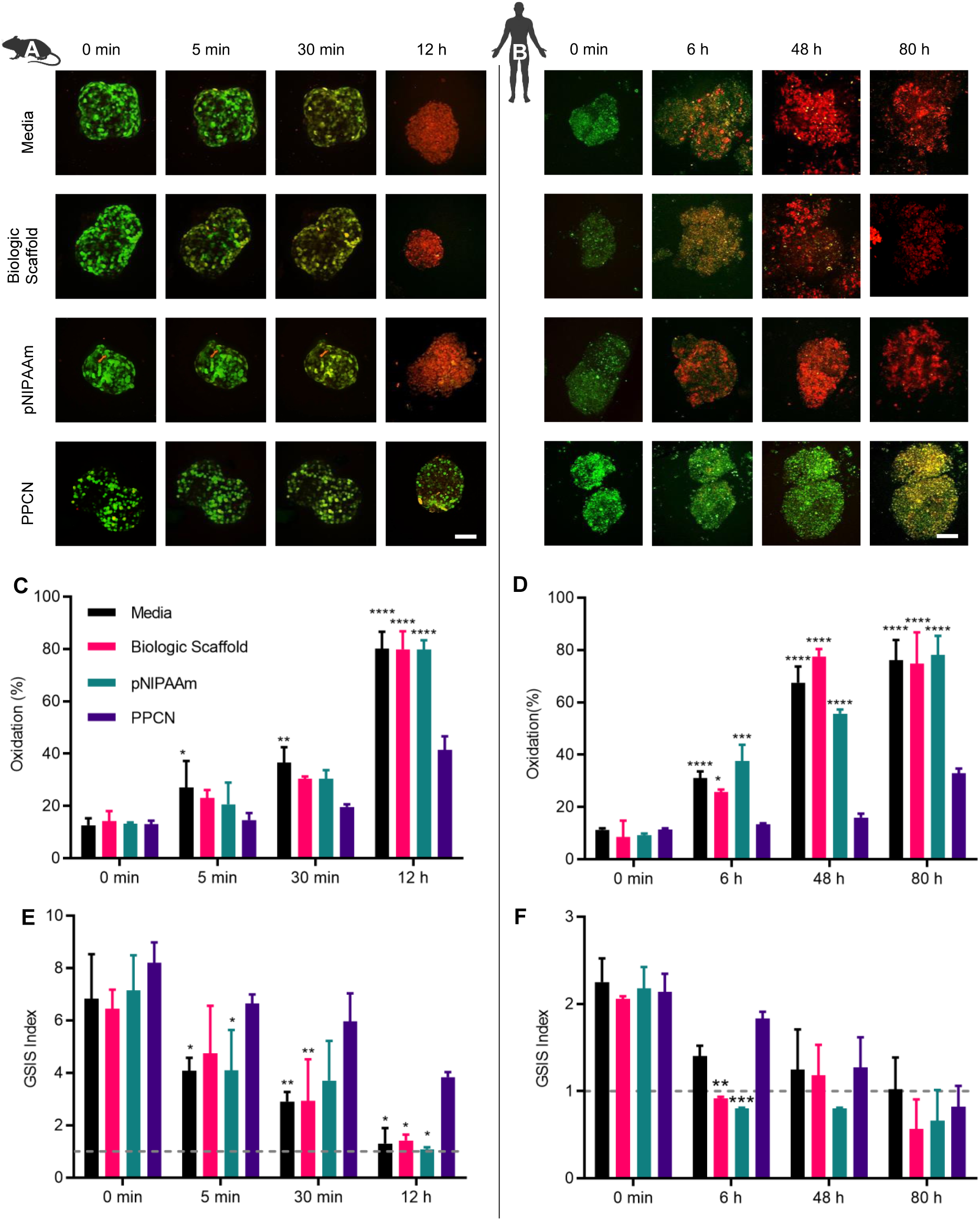
PPCN protects mouse and human islets against physiologic oxidative stress levels induced *in vitro*. (**A**,**B**) Oxidation rate of mouse and human islets, respectively, overexpressing roGFP under various culture conditions. 10 µM H_2_O_2_ was used to induce oxidative damage. (Green–488 nm excitation signal, reduced status; Red–405 nm excitation signal, oxidized status. (scale bars: 100 µm). (**C**,**D**) Quantification of oxidation (%) in roGFP-overexpressing islets. (**E**,**F**) Glucose-stimulated insulin secretion (GSIS) index of islets stressed with H_2_O_2_. All data are presented as mean ± SD with *p<0.05; ** p<0.01; *** p<0.001; **** p<0.0001 relative to PPCN. Statistical significance was determined by two-way ANOVA with Tukey’s multiple comparisons test. (n = 5).

To study the correlation between the progression of oxidative damage and loss of islet function, the roGFP-measured oxidation was compared to GSIS (**Fig. 3A-F**). Six hours after introducing H_2_O_2_, an increase in oxidation to 31.1%, 25.8%, 37.6 %, and 13.5% was measured for human islets in control media suspension culture, BS, pNIPAAm, and PPCN, respectively (**Fig. 3B,D**). PPCN’s protective effect is more evident at the 80-hour time point, as 76.3% of the human islets cultured in the other three environments showed signs of oxidation compared to 34.0% of those cultured in PPCN (**Fig. 3D**). Significant islet disaggregation was observed in the control media suspension culture, BS, and pNIPAAm groups, whereas the morphology of the islets entrapped in PPCN remained intact. GSIS index of both mouse and human islets were measured on similar timepoints relative to oxidation studies (**Fig. 3E,F**). For the mouse islets, a decrease in the GSIS index was observed as early as 5 minutes after introducing the H_2_O_2_. Relative to t=0, at 5 minutes the GSIS index for islets suspended in cell culture media, BS, pNIPAAm, and PPCN decreased by 40.2%, 26.5%, 42.7%, and 18.9% to 4.08, 4.74, 4.09 and 6.64, respectively. At 30 minutes, the GSIS index for islets in media, BS, pNIPAAm, and PPCN decreased to 2.90, 2.93, 3.70, and 5.97, respectively. At 12 hours, islets in media, BS, and pNIPAAm lost glucose responsiveness, as indicated by GSIS of approximately 1. A GSIS equal to or less than 1, specifies that the islets are not properly sensing glucose concentration and responding with the appropriate insulin secretion (*38*). In contrast, at 12 hours, islets in PPCN were able to maintain a GSIS stimulation index of 3.84, confirming the protective properties of PPCN (**Fig. 3E**). To assess whether the results obtained with murine cells would also translate to human cells, human islets were evaluated using the same experimental set-up (**Fig. 3F**). The results demonstrated a significant loss of function at the 6-hour time point, as the GSIS index dropped from 2.15 to 1.40, 2.06 to 0.92, 2.18 to 0.80, and 2.13 to1.83 for islets in media suspension culture, BS, pNIPAAm, and PPCN, respectively. Human islets in media suspension culture, BS, and pNIPAAm completely lost insulin secretion response to glucose. In contrast, PPCN reduced islet function loss as per GSIS results, demonstrating its superiority to the other test materials. (**Fig. 3F**).

### PPCN is a versatile islet delivery vehicle that preserves islet function *in vivo*

To evaluate whether PPCN could be used to deliver islets to an extrahepatic site, the abdominal fat pad of the mouse was used to mimic islet transplantation to the omentum in humans. This model was selected because both structures are well-vascularized fat tissue located in the intraperitoneal cavity (*42*). The key steps for the islet transplantation procedure are summarized in **Fig. 4A**. Upon application to the fat pad, complete gelation of the PPCN occurred within seconds of contact with the tissue, securing all the islets on the fat pad. No suturing or tissue glue was required for this step due to the tissue-adhesive nature of PPCN. The entire procedure was accomplished within 5 min. In contrast, BS took longer to solidify via the thrombin crosslinking reaction, making it difficult to control the final location of the islets. Additional control groups were included in the study. Islet transplantation to the kidney capsule (**KC**) was included as a positive control as it is widely used as an extrahepatic islet transplantation location in small animals. However, KC transplantation is not performed in humans due to anatomical differences (*17*). Furthermore, intraportal islet transplantation was used as an additional positive control, as this is the clinically used site of transplantation in humans. However, with this site, it is not possible to retrieve the transplanted islets while keeping the animal alive to ensure that any changes in blood glucose (**BG**) are a result of the transplanted islets and not residual pancreatic function.

**Fig. 4.**
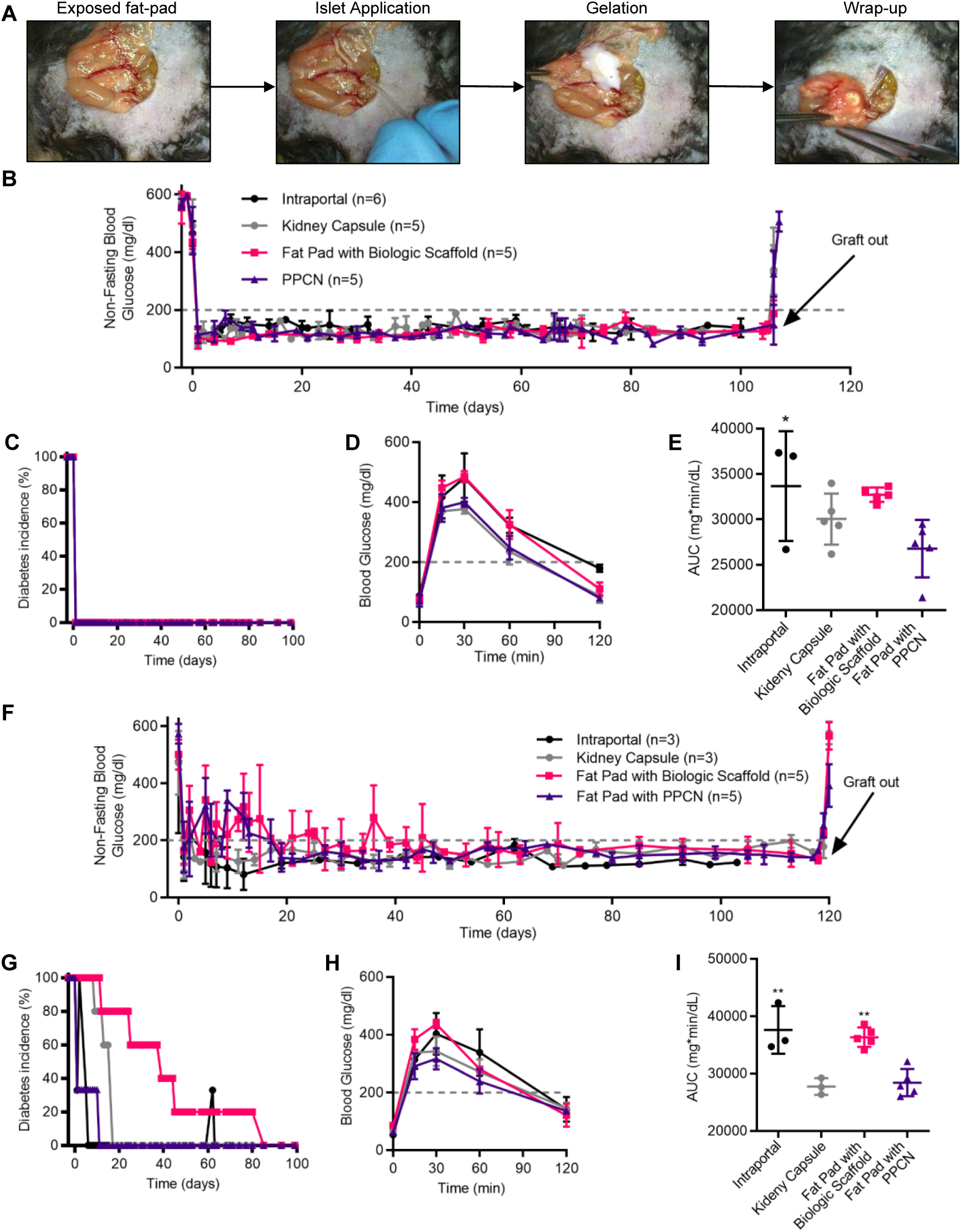
PPCN localizes islets to a target transplantation site and restores normoglycemia with marginal mass. (**A**) PPCN facilitates islet transplantation to the mouse fat pad. (**B**) Non-fasting blood glucose concentration (mg/dl) of mice transplanted with approximately 8,200 IEQ/kg body weight of islets to the liver (intraportal), kidney capsule or on the fat pad with biologic scaffold or PPCN. (**C**) Diabetes incidence (%) (blood glucose concentration greater than 200 mg/dl) by treatment group. (**D**,**E**) Glycemic profile (**D**) and area under the curve of the profile (AUC, mg * min * dL^-1^) (**E**) during the IPGTT study performed at 1 month after the transplantation. (**F**) Non-fasting blood glucose measurements of mice transplanted with a marginal islet mass (approximately 4,100 IEQ/kg body weight). (**G**) Diabetes incidence (%) (blood glucose concentration greater than 200 mg/dl) by treatment group. (**H,I**) Glycemic profile (**H**) and AUC (**I**) of the profile during the IPGTT study performed at 1 month after the transplantation. All data are presented as mean ± SD with *** p<0.001 relative to PPCN. Statistical significance was determined by two-way ANOVA with Tukey’s multiple comparisons test. (n ≥ 3).

When transplanting 8,200 islets equivalent (**IEQ**) per kilogram body weight (*43*), animals in all three groups achieved euglycemia the day following the transplantation procedure (**Fig. 4B,C**). Euglycemia was maintained in all four groups until day 104 post-transplantation, at which point a second survival surgery was performed to remove the transplanted islet graft to confirm the source of insulin production (with the exception of the intraportal group). After recovering from the surgery, hyperglycemia was detected in all animals, confirming that the islets contained within the fat pad (or KC) were responsible for maintaining euglycemia (100% converted to euglycemia, n=5) (**Fig. 4B**).

Intraperitoneal glucose tolerance tests (**IPGTT**) were performed 30 days post-transplantation. The quantified area under the curve (**AUC**) for the PPCN group was slightly lower than those for the KC and BS groups, but no significant difference was observed using 8,200 IEQ islets for transplantation (**Fig. 4D,E**).

Given the complex morphology of a pancreatitis patient’s pancreas, islet isolation is technically challenging. Thus, the islet yield from a pancreatitis patient is significantly lower compared with a standard donor (*44*). To investigate the ability of PPCN to preserve islet function and maintain normoglycemia under conditions associated with pancreatitis, a marginal mass of islets was transplanted for the subsequent study. When 4,100 IEQ/kg body weight was used (less than one donor per animal) (*45*), animals that received islets in the liver (intraportal), KC, abdominal fat pad via BS, and abdominal fat pad via PPCN achieved euglycemia within 5.3±0.6, 4.3±3.7, 25.3±13.5, and 13±4 days, respectively (**Fig. 4F,G**). IPGTT performed at one-month post-transplant shows that animals transplanted with BS had higher BG at the 15-, 30- and 60-minute time points, while BG values for the PPCN group were comparable to those of the KC control (**Fig. 4H**). The AUC in the BS group is significantly larger than those of the PPCN and kidney control groups (p<0.01) (**Fig. 4I**). The islet graft was explanted at 120 days post-transplantation. Upon graft removal, euglycemic animals that had received islets via PPCN or BS to the fat pad or in suspension to the KC all reverted to the hyperglycemia state within 48 hours.

### PPCN reduces inflammation, mitigates DNA oxidative damage, and supports neovascularization of transplanted islets

Histological and immunofluorescence staining was performed on the explanted islet grafts (fat pad and KC). Masson’s trichrome (**MT**) and hematoxylin & eosin (**H&E**) staining were used to assess collagen production and islet graft morphology (**Fig. 5A left**). Antibody probes against insulin and alpha-smooth muscle actin (**α-SMA**) confirmed the production of insulin, the presence of intact islet structures, and intra-islets neovascularization in the transplanted islets (**Fig. 5A right**). Both BS and PPCN were completely absorbed, leaving islets surrounded by native adipose tissue. Few collagen fibrils and inflammatory cells were observed in the MT-stained sections at the transplant area, suggesting the absence of BS or PPCN-induced chronic foreign body response. Because increased oxidative stress has previously been reported to be closely associated with pancreatitis-related islet damage (*15, 46, 47*), co-staining of the 8-OHdG marker was conducted to evaluate oxidation-induced DNA damage in the islet grafts (**Fig. 5B**). The expression of 8-OHdG was significantly higher in islets engrafted in the fat pad using BS whereas no signal was observed from islets engrafted with PPCN. Although many 8-OHdG positive cells were also observed in the kidney control group, unlike the BS group, the majority of the 8-OHdG positive cells in the KC group were present in the native kidney tissue and not in the grafted islets. There was significantly (p<0.05) more vascularization throughout the islets engrafted using PPCN relative to BS according to H&E and α-SMA staining (**Fig. 5C**). The favorable islet engraftment due to PPCN is further confirmed by the larger size of the islets relative to islets transplanted using BS (**Fig. 5D**).

**Fig. 5.**
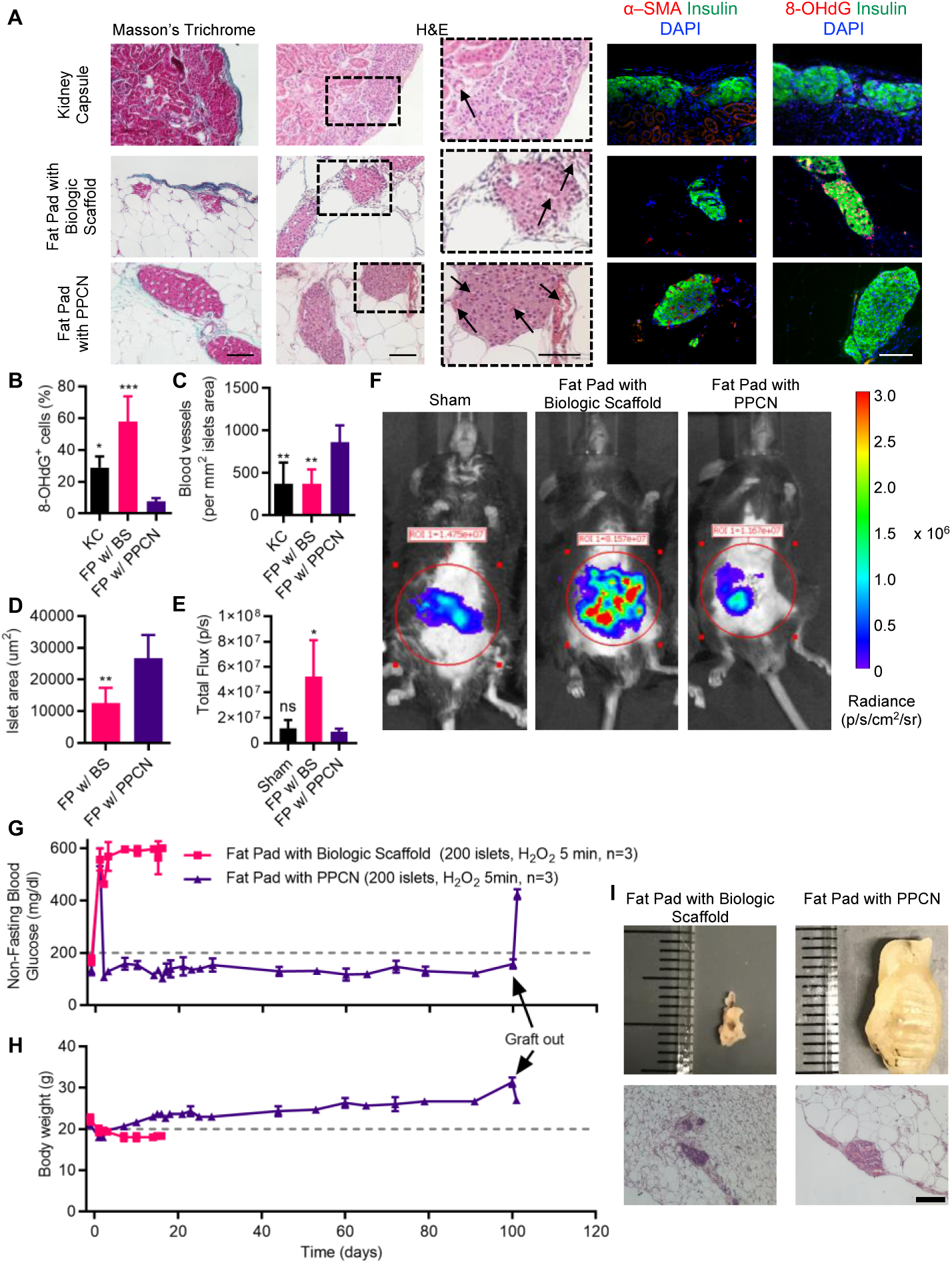
PPCN protects transplanted islets against oxidation-induced DNA damage *in vivo*. (**A**) Representative histology and immunofluorescence images of islets transplanted to the kidney capsule (KC), fat pad with biologic scaffold (FP w/ BS) or fat pad with PPCN (FP w/ PPCN), including, from left to right: Masson’s trichrome, hematoxylin and eosin (H&E), anti-α-SMA (red) and anti-8-OHdG (red) with anti-insulin (green) and nuclear dye DAPI (blue) counterstain. For H&E staining, arrows indicate the presence of blood vessels. (scale bars: 100 µm). (**B**) Quantification of nuclear oxidation based on 8-OHdG positive cells in immunofluorescence images. (**C**) Quantification of intra-islet vascular density using α-SMA positive structures in immunofluorescence images. (**D**) Quantification of islet size by area in FP groups (w/ BS or PPCN). (**E**,**F**) Quantification (**E**) and digital images (**F**) of reactive oxidative and nitrogen species *in vivo* 24 hours post transplantation as measured via IVIS by total flux of L-012 activity. (**G-I**) To evaluate protective antioxidative effects *ex vivo*, islets were pre-stressed with 10 μM H_2_O_2_ for 5 min *in vitro* prior to the transplantation. (**G**) Non-fasting blood glucose concentration (mg/dl) of mice transplanted with approximately 8,200 IEQ/kg body weight of pre-stressed islets to the FP w/ BS or PPCN. (**H**) Body weight of mice transplanted with approximately 8,200 IEQ/kg body weight of pre-stressed islets to the FP w/ BS or PPCN. Mice in the FP w/ BS group were euthanized at day 18 post-transplantation due to severe weight loss. (**I**) Digital images (top) and H&E histology (bottom) of the fat pad explanted at 18 days for the BS group or 100 days for the PPCN group. All data are presented as mean ± SD (n ≥ 3; ** p<0.01). All data are presented as mean ± SD with *p<0.05; ** p<0.01; *** p<0.001 relative to PPCN. Statistical significance was determined by T-test, one-way or two-way ANOVA with Tukey’s multiple comparisons test. (n ≥ 3).

Given that it has been reported that initial oxidative stress on the islets has been shown to negatively impact engraftment (*15, 48*), we assessed reactive oxidative and nitrogen species at 24 hours post-transplantation, in real-time, using an L-012 probe (*40*). Analysis by IVIS revealed that islet transplantation with BS cause a significant (p<0.05) increase in reactive species at the site of transplantation, whereas reactive species in the PPCN treatment group resembled the sham condition (**Fig. 5E,F**). Animals that received islets transplanted via PP**<u>G</u>**N, the non-antioxidant version of PPCN, exhibited higher oxidative stress within the fat pad than those that received islets using PPCN (**Fig. S4**).

To further evaluate PPCN for the protection of islets against oxidative tissue damage *in vivo*, freshly isolated islets were exposed to H_2_O_2_ in cell culture medium, at a concentration (10 µM) mimicking that secreted by pancreatitis-induced acinar cells (*41*). After 5 minutes of H_2_O_2_ exposure, the H_2_O_2_-containing medium was removed, and the islets were transplanted into the fat pad of syngeneic hyperglycemic recipient mice that had previously undergone chemical (streptozotocin) pancreatectomy. Non-fasting BG levels of these graft recipients were closely monitored before and after the transplantation. Although no significant morphological changes and oxidation damage were observed after 5 minutes of exposure to H_2_O_2_ according to *in vitro* culture studies, the *in vivo* performance of these H_2_O_2_ exposed islets is significantly different (**Fig. 5G**). Euglycemia was established in the recipients that received islets entrapped using PPCN within 24 hours post-transplantation, similar to the mice that received 8,200 IEQ/kg body weight (**Fig. 3A**). In contrast, all BS islets grafts exposed to H_2_O_2_ remained hyperglycemic and animals had to be euthanized 15 days post-transplantation due to significant weight loss (**Fig. 5G,H**). A dramatic difference in the tissue volume at the transplant site was also observed at the time of graft removal. Islets grafted using PPCN were approximately 10 times the size of the islets grafted using BS (**Fig. 5I**). These results highlight the destructive effects of oxidative stress on the *in vivo* function of islets post-transplantation and PPCN’s capacity to protect islets against oxidative damage and associated loss of function.

### PPCN is well tolerated, does not elicit a deleterious foreign body response, and is resorbed when applied to the omentum of non-human primates (NHPs)

Unlike the abdominal fat pad found in small rodents, the omentum in large animals such as NHPs and humans is a natural defense mechanism for the pathophysiology of intra-abdominal diseases due to its well-vascularized structure and angiogenic properties (*49*). The pro-inflammatory environment of the omentum in a human could impact islet function; hence, the compatibility of the scaffold with this tissue is of utmost importance (*49*). The similarity in islet architecture and function, as well as the size and anatomy of the omentum between humans and NHPs, motivated us to investigate the tissue response to PPCN in NHPs, specifically Rhesus macaques (*50*). PPCN application to the omentum via laparotomy is shown in **Fig. 6A**. PPCN can easily be applied as a liquid through syringes and rapidly transitions into an opaque hydrogel within seconds upon contact with the tissue at body temperature. The graft was secured and covered with surrounding omentum tissue without the need for sutures or staples. Over the course of the 3-month study, the health of the animals, including disposition, body weight, complete blood count, and chemistry, was carefully monitored. Body weight was maintained or increased (**Fig. 6B**). Kidney health, as assessed by blood urea nitrogen (**BUN**) and creatinine concentrations showed no sustained impairment in function (**Fig. 6C,D**). Similarly, liver function, as measured by bilirubin and aspartate aminotransferase (AST), was maintained throughout the study (**Fig. 6E,F**). Complete blood counts, blood chemistries, urinalysis, and body temperature for all animals over the course of the study are documented in **Tables S1-S4**. At approximately 3 months post-implantation, the implantation site was surgically accessed and inspected for any signs of inflammation or a foreign body response to the PPCN (**Fig. 6A Explant left**). Internal inspection of the implant via laparoscopic camera was normal. In all animals, a small amount of white matter that appeared to be remaining PPCN (∼20% of originally applied material) was observed. The entire omentum was extended from the body cavity for gross inspection (**Fig. 6A Explant middle**). No signs of a foreign body response were noted. The entire omentum was explanted for histopathology. The histopathology report confirmed the absence of inflammation, fibrosis, or tissue abnormalities, except for a few areas that showed signs of remaining PPCN. Tissue composition, including vascularization, also appeared normal (**Fig. 6A Explant right**). In all, PPCN applied to the omentum was well tolerated by large NHPs, as the animals maintained their baseline health throughout the study with no changes in behavior, blood parameters, or omental inflammation.

**Fig. 6.**
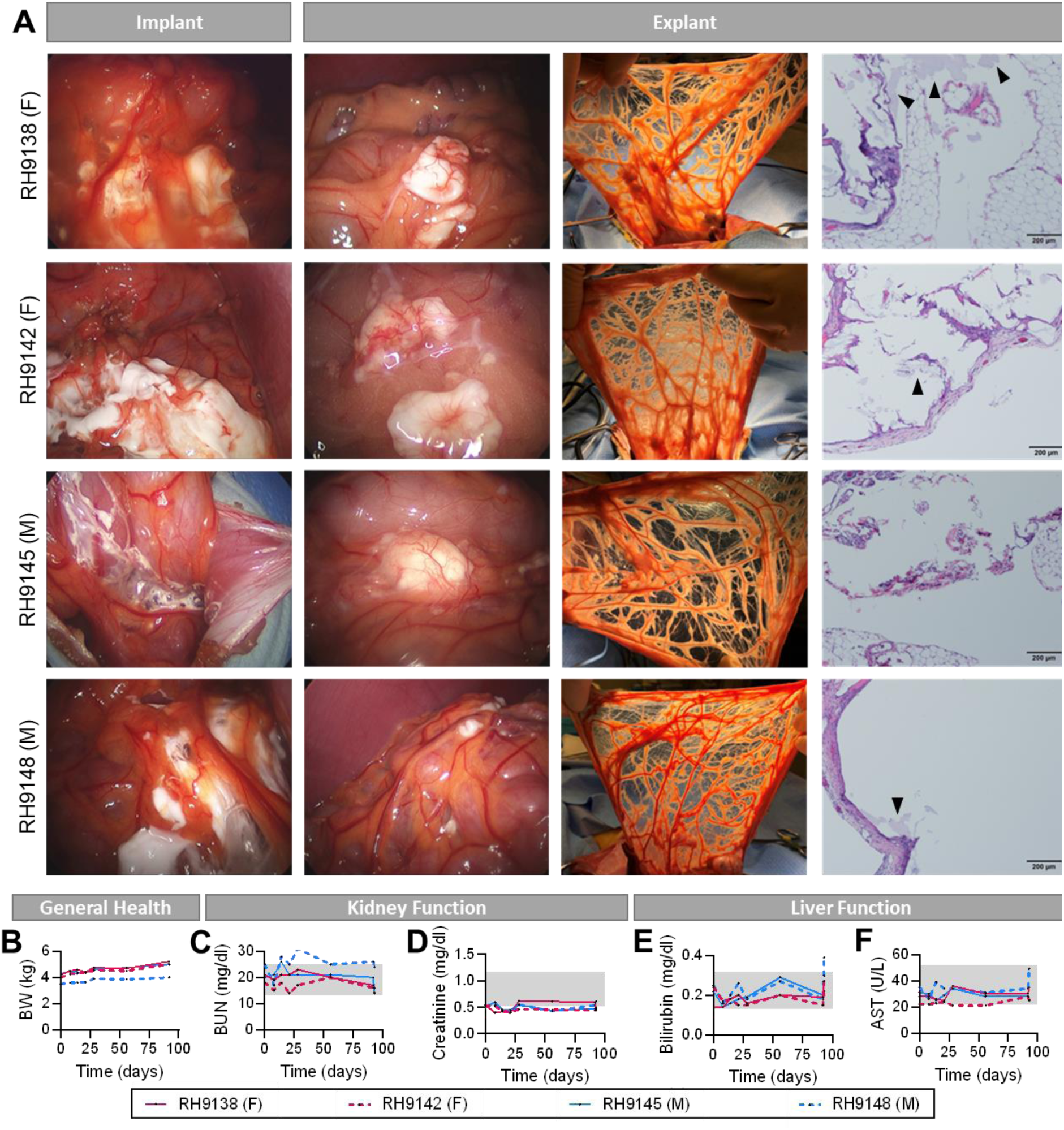
PPCN is biocompatible with the nonhuman primate omentum. Approximately 4 ml of poly(polyethylene glycol citrate-co-N isopropyl acrylamide) (PPCN) was implanted via laparotomy in the omentum of each of the Rhesus macaque nonhuman primates (NHPs) (n=4; 2 female (F), 2 male (M)). Two batches of PPCN were utilized. Batch A was implanted in RH9138 (F) and RH9145 (M). Batch B was implanted in RH9142 (F) and RH9148 (M). Explantation occurred approximately 3 months after implantation. (**A**) Implant: Laparoscopic digital photos of PPCN implantation in the omentum of 4 Rhesus macaques (2 female, 2 male). Explant: Left—Laparoscopic digital photos of PPCN within the omentum of 4 Rhesus macaques (2 female, 2 male) approximately 3 months after implantation. Middle— Digital photos of the omentum upon explanation. Right—Hematoxylin and eosin (H&E) histology of explanted omentum following 3-month implantation of PPCN. (**B-F**) Assessment of general health, kidney function and liver function in NHPs with omentum PPCN implant. (**B**) Body weight (kg) was used to assess general health. (**C,D**) Kidney function was monitored via blood urea nitrogen (BUN; mg/dl) (**C**) and creatinine (mg/dl) (**D**) concentration in serum. (**E**,**F**) Liver function was assessed via bilirubin (mg/dl) (**E**) and aspartate aminotransferase (AST; U/L) (**F**) concentration in serum. Day 0 values were measured preoperatively and serve as a baseline. Gray boxes represent the normal ranges for each parameter. See Tables S1-S4 for body temperature, complete blood counts, blood chemistries, and urinalysis for all animals over the course of the study.

### PPCN enables islet function following autologous islet transplantation to the omentum of nonhuman primates (NHP)

To model the treatment for chronic pancreatitis with TP-IAT in an NHP model (**Fig. 7A**), first, a distal pancreatectomy was performed, followed by chemical induction of diabetes using streptozotocin (*51, 52*). Isolated islets were cultured overnight and stained to assess viability (left) and purity (right) (**Fig. 7B**). The viability was 75±32% for RH9139 and 93±15% for RH9144. Purity was over 95% for both animals. A glucose-stimulated insulin secretion (GSIS) assay was performed. The stimulated index was 1.0±0.1 and 2.5±0.5 for RH9139 and RH9144, respectively. The transplanted islet dose was 39,473 IEQ and 49,638 IPN (6,073 IEQ/kg) for RH9139 and 68,685 IEQ and 61,811 IPN (7,717 IEQ/kg) for RH9144. H&E staining of the residual pancreatic tissue upon necropsy confirmed the absence of native, endogenous islets (**Fig. 7C**). The following day after the pancreatectomy, autologous islet transplantation was performed in the omentum via laparotomy using approximately 4 ml of PPCN (**Fig. 7D**). NHPs were monitored closely for over 100 days. Over the course of the study, the health of the animals, including disposition, body weight, blood chemistry, and BG, was carefully monitored. Body weight was maintained or increased (**Fig. 7E**). Kidney health, as assessed by blood urea nitrogen (**BUN**) and creatinine concentrations showed no sustained impairment in function (**Fig. 7F,G**). Similarly, liver function, as measured by bilirubin and AST, was maintained throughout the study (**Fig. 7H,I**). Complete blood counts, blood chemistries, and urinalysis for both animals over the course of the study are documented in **Tables S5-S7**. BG was monitored (**Fig. 7J Left axis**) and insulin glargine was given on a sliding scale (**Fig. 7J Right axis**) as needed. For RH9139, for the first month following transplantation, BG was maintained at an average (mean) of 186 mg/dl with an average of 5.1 U of insulin administered per day. After 1 month, the average (mean) BG decreases to 123 mg/dl and the exogenous insulin requirement drops to an average of 2.8 U per day. For RH9144, for the first month following transplantation, BG averages 190 mg/dl with an average of 5.2 U of insulin administered. After 1 month, the average BG drops to 134 mg/dl and the exogenous insulin requirement drops to an average of 3 U per day. C-peptide was detectable in both animals throughout the course of the study. Upon a survival omentectomy, C-peptide was no longer detectable, confirming C-peptide secretion was due to autologous islets transplanted on the omentum and not due to any remaining native, endogenous islets post-pancreatectomy and post-STZ administration (**Fig. 7K**). Upon explanation, functional insulin-producing islets are found within the omentum of both animals (**Fig. 7L**).

**Fig. 7.**
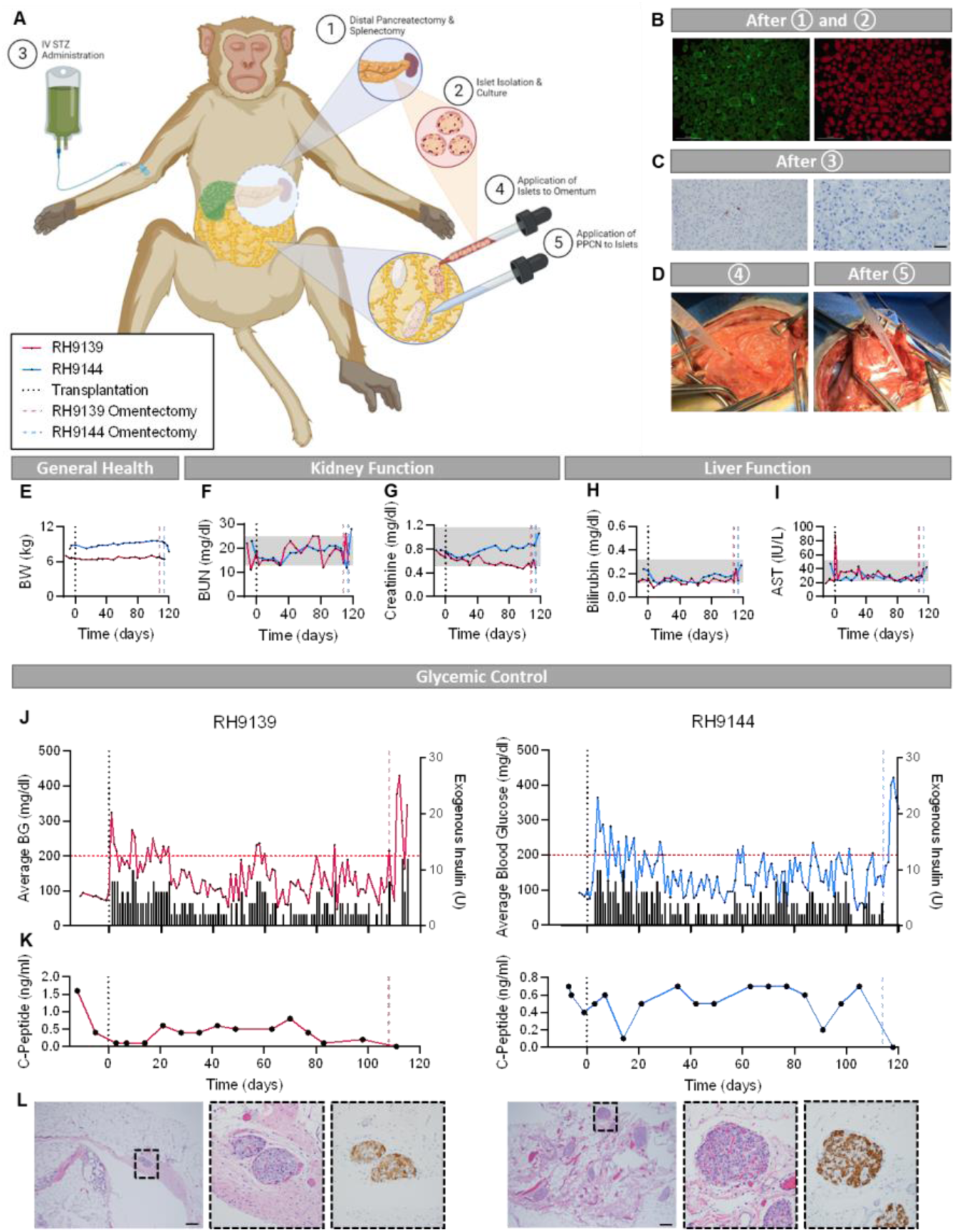
PPCN enables autologous omentum islet transplantation in a nonhuman primate model. (**A**) Schematic detailing the procedure for distal pancreatectomy-autologous islet transplantation to the omentum using poly(polyethylene glycol citrate-co-N isopropyl acrylamide) (PPCN) in a nonhuman primate (NHP). On day −1, distal pancreatectomy was performed on Rhesus macaque nonhuman primates (NHPs) (n=2; 1 female (F), 1 male (M)). To ensure all native islets were destroyed, the distal pancreatectomy was followed with chemical induction of diabetes via streptozotocin IV (STZ; 80 mg/kg). Islets were isolated and cultured overnight. (**B**) Isolated islets were stained to assess viability (left) and purity (right), using fluorescein diacetate (FDA; green)/propidium iodine (PI) and dithizone, respectively. (**C**) Hematoxylin and eosin (H&E) histology was performed on the head of the pancreas to confirm total pancreatectomy. (**D**) On day 0, autologous islet transplantation was performed via laparotomy in the omentum using approximately 4 ml of PPCN. NHPs were monitored closely for over 100 days. Blood glucose (BG) was monitored via FreeStyle Libre 2 Sensors and/or AccuCheck Guide blood glucose meter. Insulin glargine was given on a sliding scale based on blood glucose. (**E-I**) Assessment of general health, kidney function and liver function in NHPs having undergone TP-IAT in the omentum with PPCN. (**E**) Body weight (kg) was used to assess general health. (**F,G**) Kidney function was monitored via blood urea nitrogen (BUN; mg/dl) (**F**) and creatinine (mg/dl) (**G**) concentration in serum. (**H,I**) Liver function was assessed via bilirubin (mg/dl) (**H**) and aspartate aminotransferase (AST; U/L) (**I**) concentration in serum. Day −7 values (for RH9139) and −6 values (for RH9144) measured before any procedures and serve as a baseline. Gray boxes represent the normal ranges for each parameter. (**J**) Average (daily) BG (mg/dl; left axis) and daily exogenous insulin glargine (U) requirements (right axis) over the course of the study. (**K**) C-peptide concentration in whole blood (ng/ml; left axis) over the course of the study. (**J,K**) The vertical black dotted lines at day 0 indicates the time of islet transplantation. The colored vertical dashed lines indicate the time of omentectomy, for each NHP. (**L**) H&E histology (left—scale bar = 250 µm—and middle) and immunohistochemistry for insulin (right) of islets embedded within the omentum after omentectomy. See Tables S5-S7 for complete blood counts, blood chemistries, and urinalysis for both animals over the course of the study.

## DISCUSSION

Since first performed in 1977, TP-IAT has been a final hope for people suffering from chronic pancreatitis (*53*). A reduction in pain occurs in 85% of patients following this procedure (*54*). This procedure has enabled CP patients to reduce their dependence on narcotics to manage pain and improve their quality of life (*54*). This is particularly important today with the growing opioid epidemic in the United States (*55*). Despite improvements in islet isolation techniques, long-term graft survival, and sustained insulin independence remain challenges with this procedure in the CP population (*4*). CP patients have a smaller islet mass (*2*) and the inflamed acinar tissue surrounding the islets secretes H_2_O_2_, promoting islet-damaging oxidative stress (*41*). With intraportal islet transplantation, up to 50-80% of islets are lost upon infusion (*11*). Given the reduced islet reserves of CP patients, additional islet loss often leads to diabetes. 3 years post-transplant, 70% of TP-IAT patients require exogenous insulin (*54*). Thus, research into alternative, safe, and efficient extrahepatic transplantation sites that meet the needs of this patient population is warranted and the motivation for our study (*56*).

Our laboratory pioneered the development of citrate-based biomaterials (CBBs) for regenerative engineering and regenerative medicine applications (*26*). These biomaterials, which can be engineered to have tailored mechanical and degradation properties, have been investigated for several applications including the regeneration or reconstruction of cardiovascular, bladder, dermal, and musculoskeletal tissues (*37, 57–64*).

CBBs have reached a major translational milestone as they are used in bioresorbable implantable medical devices recently cleared by the U.S. Food and Drug Administration for use in musculoskeletal surgeries with commercial applications in ankle, knee, and shoulder reconstruction (*65*). We have developed a temperature-responsive, phase-changing, easy-to-use, citrate-based macromolecule, PPCN, with intrinsic anti-inflammatory and antioxidant properties. We hypothesized that PPCN protects islets isolated from CP patients against oxidative stress and prolongs their function *in vitro* and during extrahepatic islet transplantation (**Fig. 1A**).

The great omentum is a location that is ideal from the clinical perspective due to its large, well-vascularized area, and accessibility via minimally invasive procedures (*4, 45*). However, since its physiological role involves protecting the peritoneal cavity from invading infectious diseases, the pro-inflammatory environment at the omentum site may result in unexpected severe inflammatory responses towards the transplanted islets and the biomaterial used to deliver them (*49*). To date, several natural and synthetic biomaterials have been investigated as vehicles to facilitate the engraftment of islets in the omentum (*66–68*). Pedraza et al., used a polydimethylsiloxane (**PDMS**) porous scaffold in an STZ-induced diabetic rat model to restore euglycemia with 10,000 IEQ/kg body weight (1,800 IEQ/rat). However, a fibrous capsule developed around the graft area according to the histology data, due to the foreign body response elicited by the use of PDMS (*69*). Modifications to the PDMS with the angiogenic growth factor platelet-derived growth factor (**PDGF-BB**) or fibrin gel were able to slightly reduce the number of islets to 8,333 IEQ/kg body weight (250 IEQ per mouse); however, it took 19 days for the recipient to achieve euglycemia. (*70*) Berman et al. demonstrated the use of a porous polyglactin and poly-P-dioxanone scaffold (Codman Ethisorb^TM^ Duan Patch) to achieve minimum exogenous insulin requirements (0.3 to 0.4 IU/kg/day) with 5093 IEQ/kg autologous islets transplanted to the omentum of cynomolgus macaques (*52*). Immunofluorescence and histological staining of the explanted islet graft demonstrated elevated host cell infiltration around the graft area. Stendahl *et al.* investigated the use of vascular endothelial growth factor (**VEGF**) and fibroblast growth factor-2 (**FGF-2**) with heparin-binding peptide amphiphile (**HBPAs**) nanofibers in a poly (L-lactic acid) scaffold. In their study, only 78% of the mice receiving the VEGF/FGF-1-releasing scaffold achieved euglycemia within 54 days after transplantation (*71*). Furthermore, bi-layered PEG and PEG-VEGF islet encapsulation systems have recently been described (*72*). However, islet transplantation with 17,391 IEQ/Kg (4,000 IEQ/rat) did not resolve hyperglycemia, indicating insufficient insulin secretion. Berman et al. evaluated autologous BS hydrogel as a vehicle to deliver human islets to the omentum with a clinically relevant number of islets (8,200 IEQ/kg body weight (1,300 IEQ/rat)). Euglycemia was achieved the day after the transplantation (*22*). However, the pro-inflammatory and pro-oxidative environment of autologous plasma from T1D patients likely contributed to graft failures during the first clinical trial of islet transplantation to the omentum in humans (*73, 74*). Therefore, to develop a clinically useful material for extrahepatic transplantation of pancreatic islets, BS and PPCN were both evaluated in this study. In mice, we evaluated two islet doses: 8,200 IEQ/kg body weight and 4,100 IEQ/kg body weight per recipient (**Fig. 4**). These islet masses represent 50% and 25% of the total islets found in healthy mouse pancreas, the latter being a substantial reduction in islet dose when compared to previous reports (*20, 75–77*). Using 8,200 IEQ/kg body weight, euglycemia was achieved in both the PPCN and BS groups confirming the non-inferiority of PPCN to BS. However, in the marginal islet study, PPCN was a superior delivery vehicle as euglycemia was achieved 13±4 days after the transplantation in contrast to the 25±13 days in the BS group (**Fig. 4G**). Delayed insulin response was also observed in the BS group when the transplanted islets were exposed to sudden BG changes via the IPGTT test, indicating insufficient glucose control. Furthermore, unlike scaffolds reported by others, PPCN was completely resorbed by the time of explantation. No cell infiltration or fibrosis was observed around the graft area and islets were incorporated into the surrounding adipose tissue with enhanced intra-islets vasculature (**Fig. 5A**).

The islet isolation process and *in vitro* cell culture have been shown to expose islets to oxidative stress, both by facilitating the generation of reactive oxygen species (**ROS**) and by hindering the adaptive upregulation of cellular antioxidants (*15, 78, 79*). Islets isolated from the CP patients are subject to ROS secreted by the surrounding inflamed acinar tissue. Furthermore, insulin-producing beta cells have significantly lower levels of antioxidant enzymes catalase, superoxide dismutase, and glutathione peroxidase, which make them particularly vulnerable to oxidative cell damage especially during ischemia-reperfusion injury (*79–82*). These findings have therefore prompted studies that use antioxidant peptides and oxygen-generating strategies to improve islet function. An example is the addition of the peptide carnosine during *ex vivo* islet culture and oxygen-generating PDMS-CaO_2_ scaffolds for islet encapsulation (*83, 84*). Although intriguing *in vitro* results have been reported, antioxidant scaffold approaches have not been evaluated *in vivo* for islet transplantation (*85*). One potential explanation for the observed superior performance of PPCN may be its intrinsic antioxidant property, which is due to the polyethylene oxide citrate moieties present within the polymer backbone. Reduced DNA oxidative damage was consistently observed in islets engrafted with PPCN as per the 8-OHdG staining.

To determine whether there is a link between reducing oxidative stress and the preservation of islet viability and insulin secretion function, oxidative stress reporter islets were created using roGFP overexpression (**Fig. 5F**). Intracellularly expressed roGFP has been used in several studies as an effective reporter protein to allow real-time non-destructive monitoring of the cellular redox state (*86, 87*). We hereby report for the first time the use of this redox probe technology to evaluate the protective properties of an antioxidant biomaterial. When exposed to H_2_O_2_-induced oxidative stress, representative of CP, both human and mouse islets entrapped in PPCN experienced a significant delay in the progression of oxidation, thereby better-preserving viability, and glucose-stimulated insulin secretion response. To further demonstrate the impact of oxidation damage during the *ex vivo* culture period on islet performance post-transplantation, oxidative stress was induced for 5 minutes in the same way as the *in vitro* studies through low-dose H_2_O_2_ treatment before the transplantation. After the transplantation, no sign of islet damage was observed in the PPCN group as euglycemia was achieved the next day after transplantation similar to results from the original 8,200 IEQ/kg study. However, in the case of the BS group, hyperglycemia persisted after the transplantation, indicating the complete loss of insulin secretion function in those islets (**Fig. 5G**). Our results provide compelling evidence regarding the link between oxidative stress *in vitro* and islet insulin secretion function *in vivo* and that an antioxidant microenvironment can preserve the insulin secretion function of the isolated islets.

Given the anatomical and immunological differences between mice and humans, we investigated the biocompatibility of PPCN within the NHP omentum. Unlike mice, both NHP and humans are bipeds that stand and walk upright. Furthermore, both species possess an omentum with the physiological capability to produce an inflammatory response. The PPCN implantation procedure was performed via laparotomy. PPCN is applied at room temperature as a clear liquid and immediately forms a white gel upon contact with the omentum due to the polymer’s thermoresponsive nature. Using body temperature to gel a macromolecule is advantageous over other methods that use enzyme- or light-activated *in situ* polymerized scaffolds, as these methods result in insufficient encapsulation due to the uneven polymerization or additional damage to the encapsulated islets as well as the surrounding native tissue (*88*). Procedures that involve long gelation times risk allowing islets to leak into the IP cavity. Unlike methods used to date, delivery of PPCN can be easily achieved via an endoscope-enabled procedure. Three months following implantation, approximately 80% of the PPCN was reabsorbed. Signs of systemic inflammation were not evident, as per blood tests and gross examination or histopathology, demonstrating the safety of PPCN when applied to the greater omentum of a large NHP.

Given the promising results of the *in vivo* biocompatibility studies, we assessed the use of PPCN for IAT in the omentum in NHPs. Total elimination of the resident islets was achieved via distal pancreatectomy, followed by chemical induction of diabetes via streptozotocin IV. This method of pancreatectomy decreased surgical time, as resection from the duodenum was not required while ensuring no viable islets remain. The islet doses of 6,073 IEQ/kg for RH9139 and 7,717 IEQ/kg for RH9144 align with this reduced yield. Isolated islets were secured onto the omentum using the PPCN gel. One month after the procedure, BG averages of 123 mg/dl and 134 mg/dl for RH9139 and RH9144, respectively, fall within the normal range for Rhesus macaques (58-153 mg/dl). Minimal insulin was required to maintain normoglycemia. For RH9144, after the 1-month engraftment period, a mean of 0.41 U/kg/day and 0.32 U/kg/day were required for RH9139 and RH9144, respectively. These results confirm the engraftment of functional islets within PPCN in the NHP omentum.

Although we have demonstrated the translational potential of PPCN as an additional treatment tool for CP patients, there are limitations to our study. Our animal models focus on the impact of CP on the islets and do not reflect the systemic inflammation that CP patients undergo after years of living with this disease. Generally, CP patients experience elevated inflammatory cytokines in their blood, including IL-1β, IL-6, IL-8, IFNγ, MIC-1, NGAL, TGFβ, and TNFα (*89, 90*). This inflammation likely plays a role in islet engraftment and potentially contributes to the negative outcomes of TP-IAT experienced by CP patients. Future work should also focus on the development of models of CP that can be used to test clinically relevant approaches for CP therapies. One may consider the use of an *in vitro* model with elevated cytokine concentions mimicking that of a CP patient. For *in vivo* work, important considerations include the use of upright animals and chemical or surgical induction of CP that accurately models pancreatic inflammation and islet loss and systemic inflammation. To facilitate the clinical use of PPCN in TP-IAT, a head-to-head study comparing intraportal islet delivery and PPCN-mediated omentum islet transplantation should be conducted in NHPs. Given our results in mice and NHPs that demonstrate improved islet transplantation outcomes using PPCN and the practical and biological challenges with intraportal islet delivery such as liver thrombosis and instant blood-mediated inflammatory reactions, phase-changing antioxidant biomaterials such as PPCN can play an important role in islet function preservation, extrahepatic engraftment, and improving the quality of life of CP patients.

## MATERIALS AND METHODS

### Study design

The overall objective of this study was to assess the ability of PPCN to preserve the viability and function of insulin-producing islets *in vitro* and *in vivo*, under oxidative stress conditions associated with CP. To achieve this goal, first, we performed *in vitro* studies with murine and human islets. Islets were encapsulated in PPCN or control materials prior to culture. We used live/dead staining and GSIS assay to measure viability and function, respectively. Transduction of murine and human islets will an oxidative report enabled us to assess redox status over time as a function of culture conditions. Next, we assessed PPCN in an STZ-induced syngeneic islet transplant murine model. PPCN was used to secure islets to the mouse fat pad. Evaluation of PPCN was assessed upon restoration of normoglycemia, response to a glucose challenge, and terminal histopathology. PPCN-enabled islet transplant to the murine fat pad was compared against fat pad transplant with BS, KC, and intraportal islet transplant using various islet does and oxidative stress conditions. Biocompatibility of PPCN was evaluated in the omentum of 4 NHPs (2 female, 2 male) for 3 months. Outcomes were assessed based on body weight, complete blood cell counts (**CBC**), white blood cell differential, blood chemistry, urinalysis, gross observation upon explantation, histopathology upon explantation, and full necropsy. To mimic a total pancreatectomy procedure, we performed a distal pancreatectomy followed by chemical induction of diabetes via STZ. Islets were isolated, cultured overnight, and transplanted to the omentum of 2 NHPs with PPCN (1 female, 1 male). NHPs were monitored for over 100 days. BG was assessed daily. Insulin was given on a sliding scale. Outcomes were measured using the same criteria as for the biocompatibility studies with the addition of normoglycemia, insulin requirements, intravenous dextrose tolerance test (**IVDTT**), and glucagon test. Histopathology, complete blood counts, blood chemistry, and urinalysis were conducted in a blinded manner.

### Human tissue

Human islets were obtained from Northwestern University Human Islet Transplant Program (Institutional Review Board exemption: STU00207825).

### Animals

8 to 12-week-old, male C57BL/6 and Balb/c mice were purchased from Jackson Labs. Mice were housed in the Center for Comparative Medicine at Northwestern University. All animal protocols were approved by Northwestern University’s Institutional Animal Care and Use Committee (**IACUC**).

4 to 10 kg rhesus macaques were purchased from approved animal vendors. Animals were negative for Herpes B, tuberculosis, simian immunodeficiency virus, simian retrovirus, and Simian T-lymphotropic virus. All animal protocols were approved by the University of Illinois at Chicago and Northwestern University’s IACUC.

### Materials

All chemicals used in the study including citric acid, poly(ethylene glycol), glycerol 1,3-diglycerolate diacrylate, poly-N-isopropylacrylamide, collagenase (type XI), dextran, and Thrombin from murine plasma were purchased from Sigma-Aldrich.

### PPCN synthesis, characterization and solution preparation

PPCN was synthesized from citric acid, poly(ethylene glycol), glycerol 1,3-diglycerolate diacrylate, and poly-N-isopropylacrylamide following the previously published method (*26*). The resulting PPCN was characterized via ^1^H-NMR and ATR-FTIR, neutralized to pH 7 with sodium hydroxide, sterilized with ethylene oxide gas sterilization, and properly vented before use. To encapsulate islets, a 100 mg/ml PPCN solution was made by dissolving lyophilized PPCN in sterile PBS.

### Murine islet isolation

Mice were first anesthetized with an intraperitoneal injection of ketamine and xylene. After a midline abdominal incision, cold collagenase solution was injected into the pancreas via the cannulated bile duct. The collagenase-infused pancreas was then dissected and incubated at 37°C for 15 min. After the digestion, the large undigested connective tissue was removed by passing the digested pancreas through a mesh screen. The filtrate was then applied to a discontinuous dextran gradient to separate islets from the remaining connective tissue fragments. After two gradient washes, the purified islets were hand-picked and counted under the microscope.

### *In vitro* islet entrapment, viability, and insulin secretion study

The entrapment of islets within PPCN and pNIPAAm was achieved utilizing their thermoresponsive nature. Islets were purified, counted, and sorted into cell strainers. Cell culture media was drained from the cell strainer immediately before adding room temperature PPCN or pNIPAAm in the islets. The islet and PPCN or pNIPAAm mixture was then incubated at 37°C for 5 mins to solidify. After the hydrogel formed, warm culture media was added to the well to support the islet’s growth before returning the plate to the incubator. The encapsulation of islets within BS gel was done in a similar fashion, except a thrombin calcium solution was added into an initial mixture of plasma and islets to solidify the BS gel before the addition of growth media.

The viability of the encapsulated islets was assessed after 24 days of incubation using two different methods: the resazurin assay for quantification (Sigma) and the Live/Dead assay for visualization (Life Technologies). Both assays were performed following the manufacturer’s protocol.

Low (2.8 mM) and high (28 mM) glucose solutions were prepared in Kreb’s buffer for the glucose-stimulated insulin secretion test. The concentrations were determined based on the NIH human islets standard operating procedure. Briefly, after the removal of growth media from the encapsulated islets, the islets were first washed with the low glucose solution and then sequentially incubated in a) low glucose equilibration solution, b) low glucose solution, and c) high glucose solution for 1 hour each. After the incubation, the solution from b) and c) were collected and measured using an insulin ELISA kit (Thermo Fisher for mouse islets, and Mercodia for human islets). The stimulation index was defined as the ratio of stimulated (high glucose) to baseline (low glucose) insulin secretion.

The test was done by comparing the different amounts of insulin secreted by the islets when subject to low (2.8 mM) and high (28 mM) glucose concentrations. The results were reported as stimulation index (SI), which is acquired by dividing the amount of insulin produced by the islets in the high glucose solution by the insulin amount produced in the low glucose solution.

### roGFP transduction and oxidation inhibition study

To access the oxidation status of the islets under varied culture conditions, freshly isolated islets were treated with engineered lentivirus encoding the roGFP gene. roGFP expression was monitored using a fluorescent microscope after the transduction. The expression level of the roGFP protein was monitored for 96 hours after the transduction of the roGFP viral vector. The reduced protein signal (488 nm) was observed to be gradually increasing and reached a maximum at 72 hours, while the oxidized protein signal remained at a minimum level at that point. Once the roGFP protein expression reached its maximum in the viral vector-treated islets, these roGFP-islets were either cultured in standard media suspension, BS, pNIPAAm homopolymer, or PPCN.

The oxidation inhibition study was carried out in 15 well glass bottom slides (ibidi). roGFP-islets were split into each well before treatment with various conditions (PPCN, pNIPAAm, or BS). Baseline (0 minutes) confocal images of the islets under each condition were taken with two excitation wavelengths (405 nm and 475 nm) and one fixed emission wavelength of 509 nm. Hydrogen peroxide was then added to the islets culture with a final concentration of 10 µM. The oxidation status of the islets was monitored under confocal microscopy at each time point. The oxidation percentage was quantified based on the fluorescent intensity under the two wavelengths using ImageJ.

### Murine syngeneic islet transplantation

Donor animals were pre-treated with STZ to induce diabetes one week before the transplantation. Three different transplant locations were applied in the study. The abdominal fat pad transplantation is used for PPGN, BS, and PPCN islet transplantation. Intraportal transplantation to the liver and KC transplantation were used as the positive control. Both procedures were conducted following previously published procedures. For the abdominal fat pad model, A small incision was first created on the abdominal part of the animal to expose the abdominal fat pad, followed by the application of room-temperature islets-PPCN suspension. Purified islets were transplanted with 40 µl of PPGN, BS, or PPCN. After the gel solidified on the fat pad, all the islets were secured in place before returning the fatpad back into the intraperitoneal cavity. After the transplantation surgery, the non-fasting BG of the animals was monitored daily for the first two weeks post-surgery, and then once a week afterward. At the end of the study, graft explant was conducted via a survival surgery. Animals were allowed to recover from the surgery, and their BG levels were monitored for another 48 hours, after which time the mice were sacrificed.

### Murine intraperitoneal glucose tolerance testing

IPGTT was performed one-month post-transplantation. The animals were fasted for 16 hours before receiving an intraperitoneal injection of 2g/kg body weight of 50% dextrose (Abbott Labs) solution. BG was measured at 0, 15, 30, 60, and 120 minutes after the glucose injection.

### Tissue collection and immunofluorescent staining

Islets containing fat pad and kidney were harvested and processed for paraffin sectioning. Immunofluorescent staining for blood vessels, insulin, cell death, and oxidation marker (8-OHdG) was performed following the manufacturer’s protocol. Digital images were acquired with a Nikon fluorescent microscope. Images were then processed with ImageJ.

### Biocompatibility and safety of PPCN in the omentum of NHPs

Rhesus macaques were placed under general anesthesia. PPCN (4 ml) was applied onto the greater omentum via laparotomy. Body temperature and weight, complete blood cell counts, white blood cell differential, and blood chemistry were measured before and at several intervals, after the surgery, until euthanasia to assess for infection, liver function, kidney function, and blood composition. At approximately 90 post implantations, the animals were euthanized. A full necropsy was performed. Histology was performed by the Veterinary Diagnostic Laboratory at the College of Veterinary Medicine at the University of Illinois Chicago. A full histopathology report was provided.

### Autologous islet transplantation to the omentum using PPCN in NHPs

Baseline BG, IVDTT, glucagon test, CBC, white blood cell differential, blood chemistries, urinalysis, body temperature and weight, and serum C-peptide were performed on Rhesus macaques (n=2; 1 female, 1 male). NHPs were placed under general anesthesia, and a distal pancreatectomy was performed, followed by IV administration of STZ (80 mg/kg). The islets were isolated and purified using previously established methods. Following overnight culture, islets were counted, assessed for viability using FDA/PI, purity using dithizone staining, and function via GSIS assay. Under anesthesia, the purified islets were transplanted via laparotomy to the omentum on NHPs. Approximately, 4 ml of PPCN was applied to secure the islets into place on the omentum. BG was monitored daily using an AccuCheck Guide BG meter or FreeStyle Libre2 continuous glucose monitor. Insulin glargine (Lantus, Sanofi, 100 U/ml) was administered on a sliding scale twice per day. Baseline assessments were repeated at regular intervals throughout the study. At over 100 days post-transplant, the animals were euthanized. A full necropsy was performed. Histology was performed by the Veterinary Diagnostic Laboratory at the College of Veterinary Medicine at the University of Illinois Chicago. A full histopathology report was provided.

### Statistical analysis

Statistical analyses were performed using Prism 6, Two-way ANOVA was used to measure differences for experiments with multiple data sets with a Tukey test performed between groups with significant differences to correct for the multiple pair-wise comparisons. A value of p ≤ 0.05 was considered to be statistically significant. Values are reported as the mean±SD.

## List of Supplementary Materials

Fig. S1 to S4

Tables S1 to S7

## Acknowledgments

This research was supported by the Integrated Molecular Structure Education and Research Center (IMSERC) at Northwestern University, which has received support from the Soft and Hybrid Nanotechnology Experimental (SHyNE) Resource (NSF grant no. ECCS-1542205), the State of Illinois and the International Institute for Nanotechnology (IIN); the Northwestern University Center for Advanced Molecular Imaging (CAMI), which is supported by NCI grant no. CCSG P30 CA060553 awarded to the Robert H. Lurie Comprehensive Cancer Center; the MRSEC program (NSF grant no. DMR-1720139) at the Materials Research Center; the IIN; and the State of Illinois, through the IIN.

## Funding

W81XWH-19-1-0230, Department of Defense: GAA

Center for Advanced Regenerative Engineering: GAA

DGE-1842165, National Science Foundation Graduate Research Fellowship Program: JAB

Dr. John N. Nicholson Fellowship: JAB

## Author contributions

Conceptualization: JAB, YZ, GAA

Methodology: JAB, YZ, XZ, PDR, IJ, DL, HN, SR, QR, JM, DC, JO, XL

Investigation: JAB, YZ, GAA

Visualization: JAB, YZ

Funding acquisition: GAA

Project administration: JAB, YZ, PDR

Supervision: JO, XL, GAA

Writing – original draft: JAB, YZ, GAA

Writing – review & editing: JAB, GAA

## Competing interests

CellTrans, Employees: PDR, IJ, DL, HN, SR, QR, JM, DC, JO

CellTrans, Financial holdings: PDR, IJ, JM, DC, JO

SNC Therapeutics, Position of influence: JAB

VesselTek BioMedical, LLC (VTB), Financial holdings: GAA

PCT Patent Application US/2012/023293, EPO application 12741537.0 Injectable Thermoresponsive Polyelectrolytes, issued 6-2018, Inventor: GAA

VesselTek BioMedical, LLC (VTB) licensing rights to patent US/2012/023293

## Data and materials availability

All data are available in the main text or the supplementary materials.

