## Supplement for "Phase-changing citrate macromolecule combats oxidative pancreatic islet damage, enables islet engraftment and function in the omentum"

Jacqueline A. Burke *et al.*

**The PDF file includes:**

Figs. S1 to S4
Tables S1 to S7

**Supplementary Figures**

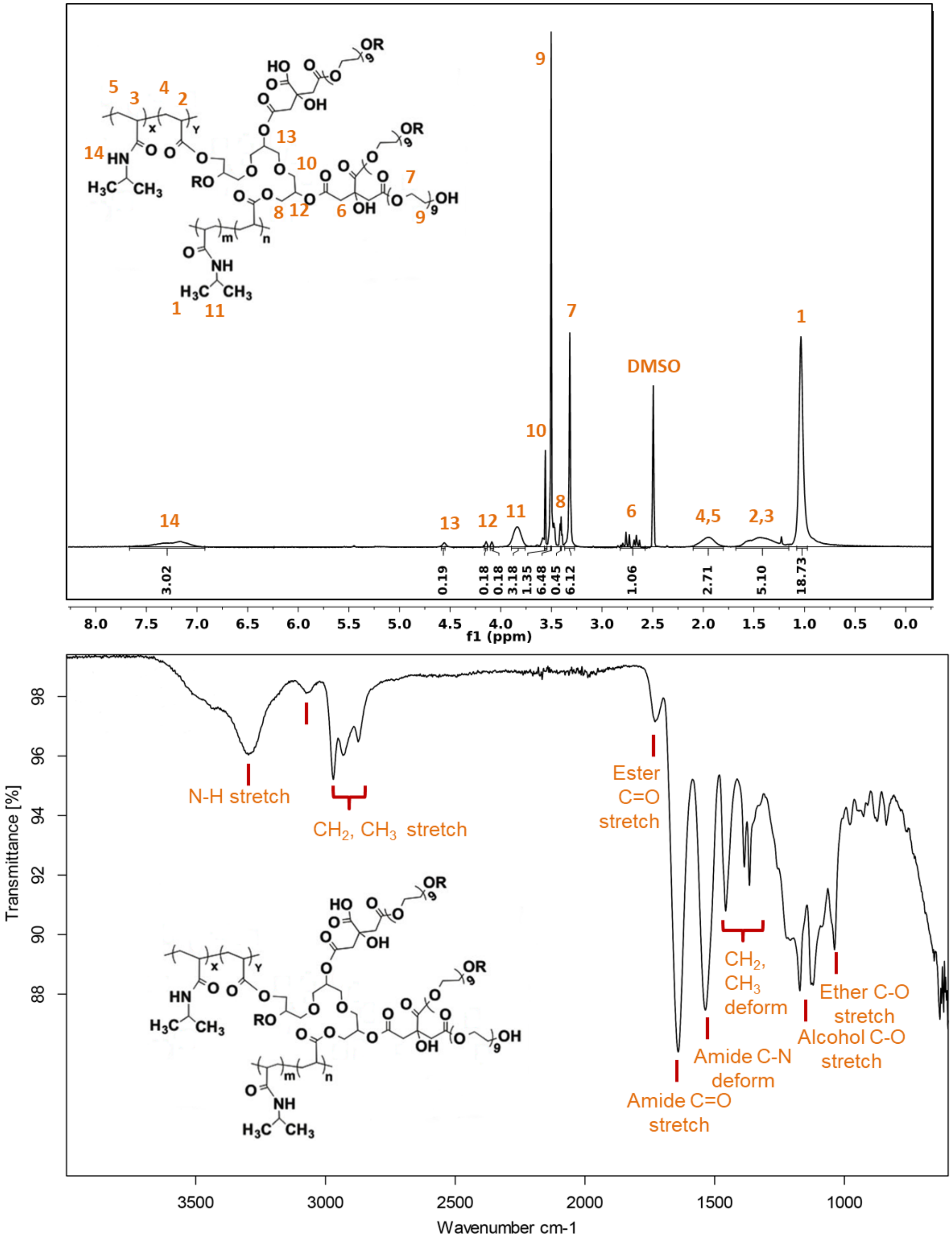


**Fig. S1. $^1$H-NMR and ATR-FTIR spectra confirm the formation of poly(polyethylene glycol citrate-co-N-isopropylacrylamide) (PPCN).**
(Top) Proton nuclear magnetic resonance spectroscopy (**$^1$H-NMR**); (Bottom) Attenuated total reflection Fourier transform inferred spectroscopy (**ATF-FTIR**).

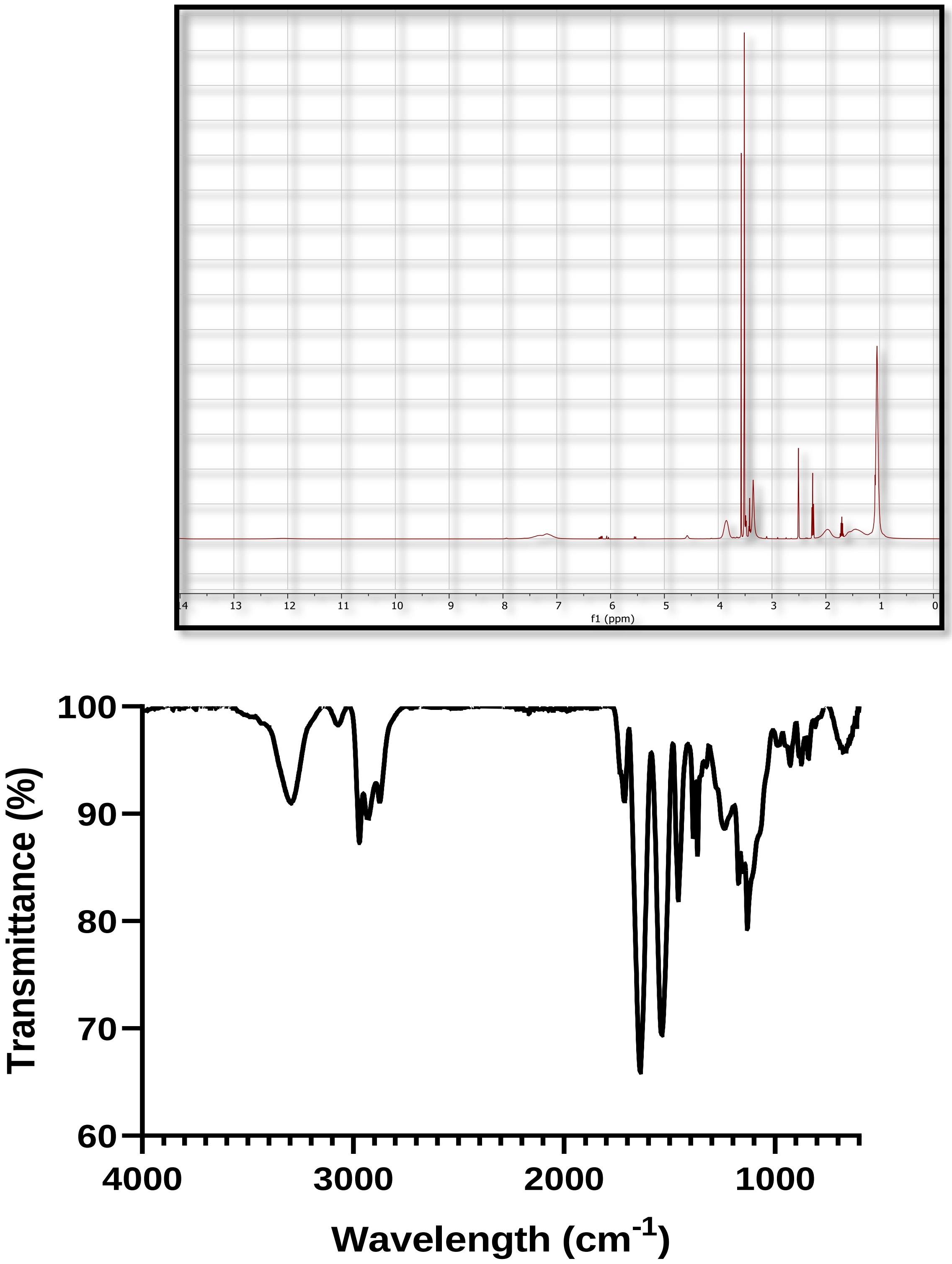


**Fig. S2. ¹H-NMR and ATR-FTIR spectra confirm the formation of poly(polyethylene glycol glutarate-co-N-isopropylacrylamide) (PPGN).**

(Top) Proton nuclear magnetic resonance spectroscopy ($^1$**H-NMR**); (Bottom) Attenuated total reflection Fourier transform inferred spectroscopy (**ATF-FTIR**).

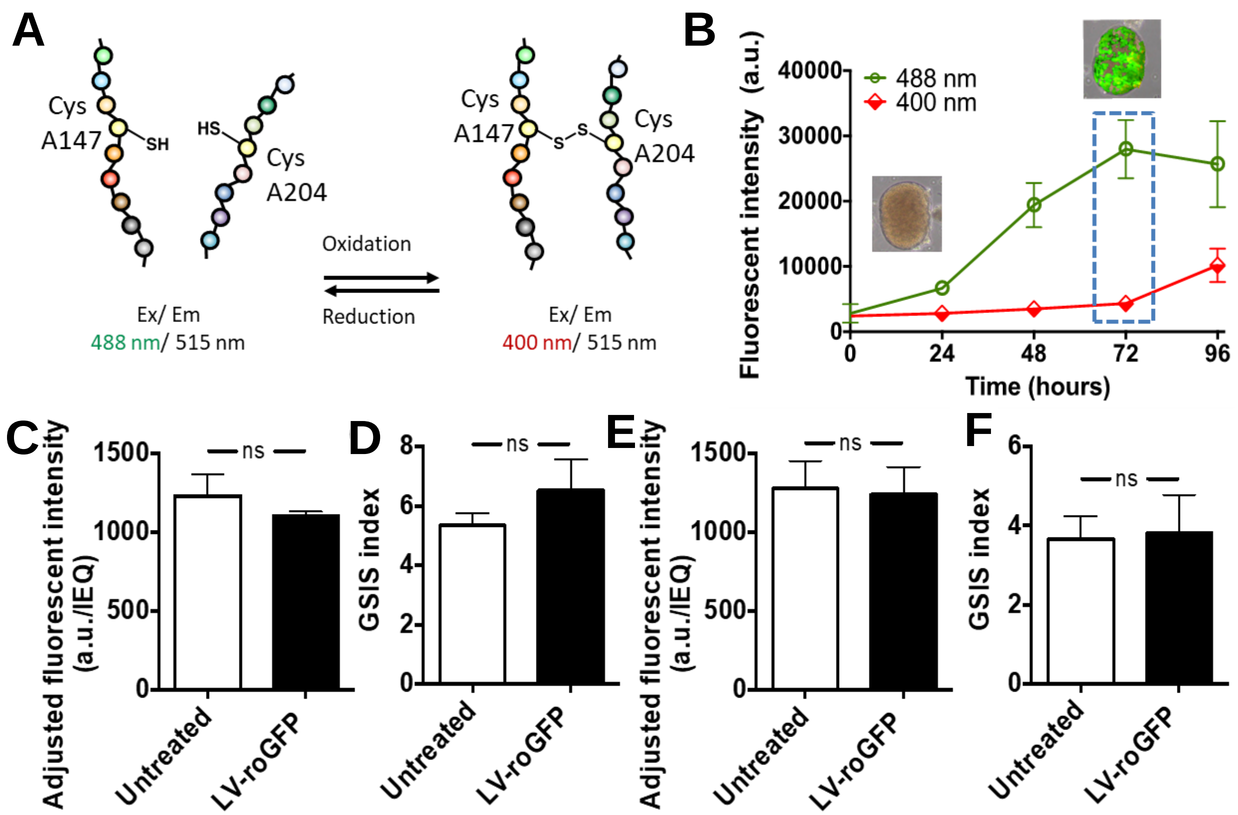


**Fig. S3. Overexpression of roGFP does not affect islet viability and insulin secretion function.**
(**A**) Schematic of the transition between the reduced and oxidized forms of roGFP. (**B**) The fluorescence intensity of reduced and oxidized RoGFP signals over time (insert: RoGFP overexpressing islets at time 0 and time 72 hours after the addition of the viral vector). **(C-F)** Viability and islet insulin secretion function were preserved after roGFP overexpression for both mouse (**C,D**) and human (**E,F**) islets. All data are presented as mean ± SD (n ≥ 3; ns: p>0.05).

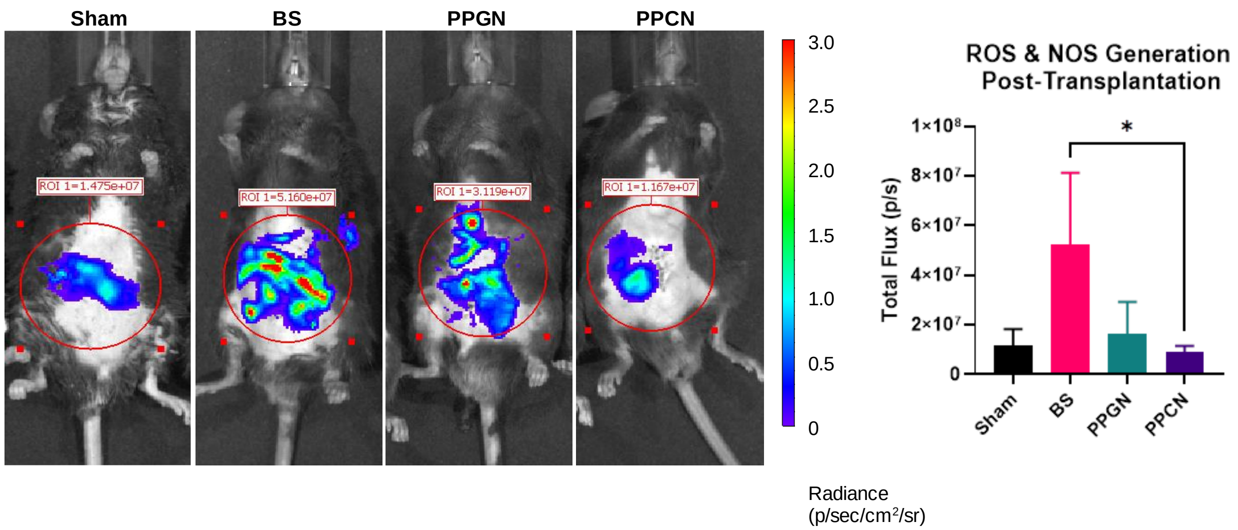


**Fig. S4. Assessment of reactive oxidative species (ROS) following islet transplantation including fat pad transplantation with PPGN.** IVIS images (**left**) and quantification (**right**) of reactive oxidative and nitrogen species *in vivo* 24 hours post-transplantation as measured via IVIS by the total flux of L-012 activity. All data are presented as mean ± SD with *p<0.05. Statistical significance was determined by one-way ANOVA with Tukey's multiple comparisons test. (n = 3).

**Table S1. Complete blood counts for NHPs over the course of the PPCN biocompatibility study.**
(n = 4; 2 female (F), 2 male (M)); WBC = white blood cells; RBC = red blood cells; HGB = hemoglobin; HCT = hematocrit; MCV = mean corpuscular volume; MCH = mean corpuscular hemoglobin; MCHC = mean corpuscular hemoglobin concentration; PLT = platelets; MPV = mean platelet volume; RDW = red cell distribution width; NEUT = neutrophils; LYMPH = lymphocytes; MONO = monocytes; EOS = eosinophils; BASO = basophils; LUC = large unstained cells; RETIC =reticulocytes; MACRO = macrocytosis; HC-VAR = hemoglobin concentration variance; PLATCLMP = platelet clumps; HYPER = hyperchromia.

| **Monkey:** | **RH9138 (F)** | | | | | | | | |
|---|---|---|---|---|---|---|---|---|---|
| **Condition:** | Day of Tx | | | | | | | Day of Retrieval | **Units** |
| **Date:** | 5/18/2020 | 5/26/2020 | 6/1/2020 | 6/8/2020 | 6/15/2020 | 7/13/2020 | 8/10/2020 | 8/18/2020 | |
| **Day:** | 0 | 8 | 14 | 21 | 28 | 56 | 84 | 92 | |
| **WBC** | 7.8 | 10.17 | 8.29 | 7.22 | 16.19 | 7.84 | 6.76 | 7.39 | x10^3/µL |
| **RBC** | 4.73 | 4.81 | 4.91 | 4.7 | 5.13 | 5.26 | 4.87 | 4.41 | x10^6/µL |
| **HGB** | 11.2 | 11.3 | 11.3 | 11.1 | 12.1 | 12.3 | 11.6 | 10.4 | g/dL |
| **HCT** | 34.8 | 34.7 | 36.3 | 34.4 | 39.4 | 37.2 | 35.6 | 32.5 | % |
| **MCV** | 73.6 | 72.1 | 73.9 | 73.1 | 76.8 | 70.6 | 73.1 | 73.8 | fL |
| **MCH** | 23.6 | 23.6 | 23.0 | 23.7 | 23.6 | 23.4 | 23.8 | 23.7 | pg |
| **MCHC** | 32.1 | 32.7 | 31.1 | 32.4 | 30.8 | 33.2 | 32.6 | 32.1 | g/dL |
| **PLT** | 356 | 353 | 353 | 352 | 403 | 277 | 327 | 257 | x10^3/µL |
| **MPV** | 7.5 | 7.9 | 7 | 6.9 | 7.4 | 6.8 | 7.2 | 6.9 | fL |
| **RDW** | 13.1 | 12.8 | 13.1 | 14.4 | 12.5 | 12.6 | 13.9 | 14 | % |
| **%NEUT** | 38.6 | 44.8 | 34.6 | 46.7 | 62.6 | 52.4 | 42.2 | 50.3 | % |
| **%LYMPH** | 57.7 | 49.3 | 57.2 | 48.2 | 34.3 | 43.4 | 53.5 | 46.1 | % |
| **%MONO** | 1.1 | 2.0 | 1.8 | 2.0 | 1.5 | 1.3 | 2.1 | 1.6 | % |
| **%EOS** | 1.90 | 3.10 | 5.40 | 2.40 | 0.80 | 2.30 | 1.30 | 1.20 | % |
| **%BASO** | 0.2 | 0.2 | 0.3 | 0.1 | 0.3 | 0.2 | 0.2 | 0.2 | % |
| **%LUC** | 0.5 | 0.5 | 0.7 | 0.6 | 0.4 | 0.4 | 0.7 | 0.6 | % |
| **#NEUT** | 3.01 | 4.56 | 2.86 | 3.37 | 10.14 | 4.1 | 2.85 | 3.72 | x10^3/µL |
| **#LYMPH** | 4.50 | 5.02 | 4.74 | 3.48 | 5.56 | 3.41 | 3.62 | 3.41 | x10^3/µL |
| **#MONO** | 0.09 | 0.21 | 0.15 | 0.14 | 0.25 | 0.10 | 0.14 | 0.12 | x10^3/µL |
| **#EOS** | 0.15 | 0.31 | 0.45 | 0.18 | 0.14 | 0.18 | 0.09 | 0.09 | x10^3/µL |
| **#BASO** | 0.02 | 0.02 | 0.02 | 0.01 | 0.04 | 0.01 | 0.02 | 0.02 | x10^3/µL |
| **#LUC** | 0.04 | 0.05 | 0.06 | 0.04 | 0.06 | 0.03 | 0.05 | 0.04 | x10^3/µL |
| **%RETIC** | 2.21 | 2.18 | 2.13 | 2.12 | 2.07 | 1.29 | 2.15 | 2.81 | % |
| **#RETIC** | 104.6 | 104.9 | 104.5 | 99.5 | 106.2 | 68 | 104.8 | 123.9 | x10^9/µL |
| **MACRO** | | | | + | + | | | | |
| **HC-VAR** | | | | + | | | | | |
| **HYPER** | | | | + | | +++ | + | + | |

**Table S1 (Continued). Complete blood counts for NHPs over the course of the PPCN biocompatibility study.** (n = 4; 2 female (F), 2 male (M)); WBC = white blood cells; RBC = red blood cells; HGB = hemoglobin; HCT = hematocrit; MCV = mean corpuscular volume; MCH = mean corpuscular hemoglobin; MCHC = mean corpuscular hemoglobin concentration; PLT = platelets; MPV = mean platelet volume; RDW = red cell distribution width; NEUT = neutrophils; LYMPH = lymphocytes; MONO = monocytes; EOS = eosinophils; BASO = basophils; LUC = large unstained cells; RETIC =reticulocytes; MACRO = macrocytosis; HC-VAR = hemoglobin concentration variance; PLATCLMP = platelet clumps; HYPER = hyperchromia.

| Monkey: | RH9142 (F) | | | | | | | | |
|---|---|---|---|---|---|---|---|---|---|
| Condition: | Day of Tx | | | | | | | Day of Retrieval | Units |
| Date: | 5/18/2020 | 5/26/2020 | 6/1/2020 | 6/8/2020 | 6/15/2020 | 7/13/2020 | 8/10/2020 | 8/18/2020 | |
| Day: | 0 | 8 | 14 | 21 | 28 | 56 | 84 | 92 | |
| WBC | 6.82 | 14.97 | 11.6 | 8.79 | 8.67 | 11.73 | 7.55 | 5.88 | x10^3/µL |
| RBC | 4.52 | 4.72 | 4.69 | 4.44 | 4.37 | 4.62 | 4.45 | 4.12 | x10^6/µL |
| HGB | 10.8 | 11.3 | 11.1 | 10.5 | 10.3 | 11.2 | 10.9 | 9.9 | g/dL |
| HCT | 33.6 | 35.1 | 35.5 | 32.4 | 33.7 | 33.7 | 33.6 | 31.3 | % |
| MCV | 74.2 | 74.3 | 75.6 | 73.0 | 77.1 | 72.9 | 75.5 | 75.8 | fL |
| MCH | 23.9 | 24.0 | 23.7 | 23.8 | 23.6 | 24.2 | 24.4 | 24.1 | pg |
| MCHC | 32.2 | 32.3 | 31.4 | 32.6 | 30.6 | 33.2 | 32.4 | 31.8 | g/dL |
| PLT | 271 | 349 | 345 | 310 | 302 | 308 | 341 | 241 | x10^3/µL |
| MPV | 7.3 | 7.2 | 6.2 | 6.2 | 6.4 | 6 | 6.1 | 6.2 | fL |
| RDW | 13 | 12.7 | 13.2 | 13.1 | 13 | 12.8 | 13.4 | 13.6 | % |
| %NEUT | 59.2 | 52.9 | 29.8 | 38.0 | 40.2 | 53.5 | 42.7 | 67.2 | % |
| %LYMPH | 38.0 | 40.9 | 61.9 | 57.3 | 54.6 | 41.8 | 51.5 | 29.9 | % |
| %MONO | 1.8 | 4.4 | 3.6 | 2.3 | 3.2 | 2.9 | 2.9 | 1.7 | % |
| %EOS | 0.70 | 1.20 | 4.10 | 1.70 | 1.40 | 0.80 | 2.00 | 0.80 | % |
| %BASO | 0.1 | 0.3 | 0.3 | 0.2 | 0.1 | 0.2 | 0.2 | 0.1 | % |
| %LUC | 0.2 | 0.4 | 0.4 | 0.5 | 0.5 | 0.8 | 0.7 | 0.3 | % |
| #NEUT | 4.04 | 7.91 | 3.45 | 3.34 | 3.49 | 6.27 | 3.22 | 3.95 | x10^3/µL |
| #LYMPH | 2.59 | 6.12 | 7.18 | 5.04 | 4.73 | 4.91 | 3.89 | 1.76 | x10^3/µL |
| #MONO | 0.12 | 0.66 | 0.41 | 0.20 | 0.27 | 0.34 | 0.22 | 0.10 | x10^3/µL |
| #EOS | 0.05 | 0.17 | 0.48 | 0.15 | 0.13 | 0.1 | 0.15 | 0.05 | x10^3/µL |
| #BASO | 0 | 0.04 | 0.04 | 0.02 | 0.01 | 0.02 | 0.02 | 0 | x10^3/µL |
| #LUC | 0.01 | 0.07 | 0.04 | 0.05 | 0.04 | 0.09 | 0.05 | 0.02 | x10^3/µL |
| %RETIC | 1.1 | 1.31 | 1.7 | 2.1 | 1.64 | 1.2 | 1.33 | 1.8 | % |
| #RETIC | 49.7 | 62 | 80 | 93.1 | 71.8 | 55.4 | 58.9 | 74.4 | x10^9/µL |
| PLTCLMP | | | | | | | | | |
| HYPER | | | | + | | + | + | | |
| MACRO | | | | | + | | + | + | |

**Table S1 (Continued). Complete blood counts for NHPs over the course of the PPCN biocompatibility study.** (n = 4; 2 female (F), 2 male (M)); WBC = white blood cells; RBC = red blood cells; HGB = hemoglobin; HCT = hematocrit; MCV = mean corpuscular volume; MCH = mean corpuscular hemoglobin; MCHC = mean corpuscular hemoglobin concentration; PLT = platelets; MPV = mean platelet volume; RDW = red cell distribution width; NEUT = neutrophils; LYMPH = lymphocytes; MONO = monocytes; EOS = eosinophils; BASO = basophils; LUC = large unstained cells; RETIC =reticulocytes; MACRO = macrocytosis; HC-VAR = hemoglobin concentration variance; PLATCLMP = platelet clumps; HYPER = hyperchromia.

| Monkey: | RH9145 (M) | | | | | | | | |
|---|---|---|---|---|---|---|---|---|---|
| Condition: | Day of Tx | | | | | | | Day of Retrieval | Units |
| Date: | 5/18/2020 | 5/26/2020 | 6/1/2020 | 6/8/2020 | 6/15/2020 | 7/13/2020 | 8/10/2020 | 8/18/2020 | |
| Day: | 0 | 8 | 14 | 21 | 28 | 56 | 84 | 92 | |
| WBC | 5.33 | 12.94 | 9.03 | 8.71 | 9.98 | 8.19 | 7.45 | 6.76 | x10^3/µL |
| RBC | 4.85 | 4.91 | 4.78 | 4.89 | 5.01 | 5.1 | 4.77 | 4.93 | x10^6/µL |
| HGB | 11.4 | 11.2 | 11.0 | 11.0 | 11.1 | 11.6 | 11.1 | 11.4 | g/dL |
| HCT | 34.9 | 35.4 | 34.4 | 34.4 | 36.6 | 35.2 | 34.4 | 35.7 | % |
| MCV | 71.9 | 72.1 | 72.0 | 70.3 | 73.1 | 69.1 | 72.2 | 72.4 | fL |
| MCH | 23.6 | 22.9 | 22..9 | 22.6 | 22.2 | 22.7 | 23.2 | 23.1 | pg |
| MCHC | 32.8 | 31.8 | 31.8 | 32.1 | 30.4 | 32.9 | 32.2 | 31.9 | g/dL |
| PLT | 322 | 389 | 352 | 369 | 384 | 347 | 394 | 315 | x10^3/µL |
| MPV | 6.6 | 6.8 | 5.6 | 5.6 | 5.8 | 5.6 | 5.7 | 5.9 | fL |
| RDW | 12.5 | 12.5 | 12.7 | 12.5 | 12.4 | 13.1 | 13.2 | 13.4 | % |
| %NEUT | 24.8 | 54.4 | 32.2 | 23.9 | 18.6 | 24.3 | 18.3 | 27.8 | % |
| %LYMPH | 70.8 | 42.0 | 62.2 | 68.6 | 75.2 | 70.6 | 77.7 | 65.8 | % |
| %MONO | 2.3 | 1.9 | 2.5 | 1.6 | 1.6 | 1.3 | 1.2 | 1.2 | % |
| %EOS | 1.20 | 1.00 | 2.00 | 4.70 | 3.40 | 2.50 | 1.50 | 4.10 | % |
| %BASO | 0.2 | 0.3 | 0.2 | 0.3 | 0.4 | 0.3 | 0.3 | 0.1 | % |
| %LUC | 0.7 | 0.4 | 0.8 | 0.9 | 0.8 | 1 | 1.1 | 1 | % |
| #NEUT | 1.32 | 7.04 | 2.91 | 2.08 | 1.86 | 1.99 | 1.36 | 1.88 | x10^3/µL |
| #LYMPH | 3.77 | 5.44 | 5.62 | 5.98 | 7.51 | 5.79 | 5.79 | 4.45 | x10^3/µL |
| #MONO | 0.12 | 0.25 | 0.23 | 0.14 | 0.16 | 0.10 | 0.09 | 0.08 | x10^3/µL |
| #EOS | 0.07 | 0.13 | 0.18 | 0.41 | 0.34 | 0.2 | 0.11 | 0.27 | x10^3/µL |
| #BASO | 0.01 | 0.04 | 0.02 | 0.03 | 0.04 | 0.03 | 0.02 | 0.01 | x10^3/µL |
| #LUC | 0.04 | 0.06 | 0.07 | 0.08 | 0.08 | 0.08 | 0.08 | 0.07 | x10^3/µL |
| %RETIC | 1.22 | 0.95 | 0.95 | 1.2 | 1.23 | 1.09 | 1.17 | 1.34 | % |
| #RETIC | 59.1 | 46.4 | 45.4 | 58.7 | 61.7 | 55.6 | 55.7 | 66.2 | x10^9/µL |
| PLTCLMP | + | | | | | | | | |
| HYPER | | | | | | ++ | | | |

**Table S1 (Continued). Complete blood counts for NHPs over the course of the PPCN biocompatibility study.** (n = 4; 2 female (F), 2 male (M)); WBC = white blood cells; RBC = red blood cells; HGB = hemoglobin; HCT = hematocrit; MCV = mean corpuscular volume; MCH = mean corpuscular hemoglobin; MCHC = mean corpuscular hemoglobin concentration; PLT = platelets; MPV = mean platelet volume; RDW = red cell distribution width; NEUT = neutrophils; LYMPH = lymphocytes; MONO = monocytes; EOS = eosinophils; BASO = basophils; LUC = large unstained cells; RETIC =reticulocytes; MACRO = macrocytosis; HC-VAR = hemoglobin concentration variance; PLATCLMP = platelet clumps; HYPER = hyperchromia.

| Monkey: | RH9148 (M) | | | | | | | | |
|---|---|---|---|---|---|---|---|---|---|
| Condition: | Day of Tx | | | | | | | Day of Retrieval | Units |
| Date: | 5/18/2020 | 5/26/2020 | 6/1/2020 | 6/8/2020 | 6/15/2020 | 7/13/2020 | 8/10/2020 | 8/18/2020 | |
| Day: | 0 | 8 | 14 | 21 | 28 | 56 | 84 | 92 | |
| WBC | 8.78 | 13.47 | 11.05 | 9.54 | 16.08 | 10.35 | 11.69 | 10.09 | x10^3/µL |
| RBC | 5.47 | 5.49 | 5.4 | 5.41 | 5.3 | 5.45 | 5.59 | 5.49 | x10^6/µL |
| HGB | 12.9 | 12.7 | 12.6 | 12.5 | 12.1 | 12.7 | 13.1 | 12.6 | g/dL |
| HCT | 40.3 | 40.5 | 40.2 | 39.2 | 40.6 | 38.7 | 41.4 | 40.8 | % |
| MCV | 73.6 | 73.8 | 74.4 | 72.5 | 76.6 | 70.9 | 74.0 | 74.3 | fL |
| MCH | 23.6 | 23.2 | 23.3 | 23.1 | 22.9 | 23.4 | 23.5 | 22.9 | pg |
| MCHC | 32 | 31.4 | 31.3 | 31.9 | 29.9 | 32.9 | 31.8 | 30.9 | g/dL |
| PLT | 357 | 387 | 347 | 349 | 382 | 282 | 332 | 342 | x10^3/µL |
| MPV | 7.5 | 7.7 | 7.3 | 6.8 | 7.2 | 6.7 | 7.3 | 7 | fL |
| RDW | 12.6 | 12.3 | 12.7 | 12.8 | 12.6 | 13.2 | 13.4 | 13.5 | % |
| %NEUT | 22.0 | 35.1 | 19.0 | 17.5 | 35.7 | 18.6 | 16.6 | 68.6 | % |
| %LYMPH | 75.4 | 62.0 | 77.4 | 79.7 | 60.5 | 79.1 | 80.8 | 29.7 | % |
| %MONO | 1.2 | 1.4 | 1.6 | 0.9 | 2.0 | 0.8 | 0.8 | 0.8 | % |
| %EOS | 0.40 | 0.70 | 0.90 | 1.00 | 0.80 | 0.30 | 0.40 | 0.60 | % |
| %BASO | 0.3 | 0.3 | 0.4 | 0.3 | 0.3 | 0.3 | 0.4 | 0.2 | % |
| %LUC | 0.6 | 0.5 | 0.7 | 0.6 | 0.6 | 0.8 | 1.1 | 0.2 | % |
| #NEUT | 1.93 | 4.72 | 2.1 | 1.67 | 5.74 | 1.93 | 1.94 | 6.91 | x10^3/µL |
| #LYMPH | 6.63 | 8.35 | 8.55 | 7.60 | 9.74 | 8.19 | 9.44 | 3.00 | x10^3/µL |
| #MONO | 0.11 | 0.19 | 0.17 | 0.09 | 0.33 | 0.09 | 0.09 | 0.08 | x10^3/µL |
| #EOS | 0.04 | 0.09 | 0.1 | 0.1 | 0.13 | 0.03 | 0.05 | 0.06 | x10^3/µL |
| #BASO | 0.02 | 0.05 | 0.04 | 0.03 | 0.06 | 0.03 | 0.04 | 0.02 | x10^3/µL |
| #LUC | 0.05 | 0.06 | 0.08 | 0.05 | 0.09 | 0.09 | 0.13 | 0.02 | x10^3/µL |
| %RETIC | 1.3 | 0.83 | 1.54 | 1.57 | 1.5 | 1.09 | 1.03 | 1.07 | % |
| #RETIC | 71.1 | 45.8 | 83.1 | 85.2 | 79.7 | 59.3 | 57.6 | 58.9 | x10^9/µL |
| PLTCLMP | | | | | + | | | | |
| HYPER | | | | | | ++ | | | |

**Table S2. Blood chemistry panel for NHPs over the course of the PPCN biocompatibility study.**
(n = 4; 2 female (F), 2 male (M)); ALT = alanine transaminase; AST = aspartate aminotransferase; BUN = blood urea nitrogen; GGT = gamma-glutamyl transferase; I = inorganic; LIP = lipemia; ICH = icterus; HEM = hemolysis.

| **Monkey:** | **RH9138 (F)** | | | | | | | | |
|---|---|---|---|---|---|---|---|---|---|
| **Condition:** | Day of Tx | | | | | | | Day of Retrieval | **Units** |
| **Date:** | 5/18/2020 | 5/26/2020 | 6/1/2020 | 6/8/2020 | 6/15/2020 | 7/13/2020 | 8/10/2020 | 8/18/2020 | |
| **Day:** | 0 | 8 | 14 | 21 | 28 | 56 | 84 | 92 | |
| Albumin | 4.008 | 3.605 | 3.692 | 4.149 | 4.343 | 4.613 | 4.149 | 3.733 | g/dL |
| Alkaline Phosphatase | 251 | 274 | 274 | 297 | 318 | 236 | 219 | 233 | U/L |
| ALT | 29 | 23 | 20 | 30 | 26 | 29 | 24 | 24 | U/L |
| AST | 28 | 28 | 26 | 25 | 36 | 30 | 30 | 35 | U/L |
| Urea Nitrogen (BUN) | 21 | 19 | 21 | 21 | 23 | 20 | 17 | 16 | mg/dL |
| Calcium | 8.85 | 8.91 | 8.43 | 9.09 | 9.25 | 9.21 | 8.81 | 8.39 | mg/dL |
| Creatinine | 0.5402 | 0.3978 | 0.4171 | 0.3939 | 0.6184 | 0.6079 | 0.5911 | 0.4869 | mg/dL |
| GGT | 63 | 68 | 65 | 65 | 67 | 59 | 48 | 45 | U/L |
| Glucose | 68 | 60 | 74 | 77 | 76 | 74 | 68 | 60 | mg/dL |
| I. Phosphorous | 6.4 | 5.9 | 5.6 | 5.2 | 6 | 4.7 | 4 | 3.9 | mg/dL |
| Total Bilirubin | 0.14 | 0.14 | 0.18 | 0.2 | 0.16 | 0.2 | 0.19 | 0.27 | mg/dL |
| Total Protein | 6.072 | 5.573 | 5.412 | 6.489 | 6.585 | 7.012 | 6.2 | 5.545 | g/dL |
| Sodium | 144 | 143 | 142 | 144 | 145 | 145 | 144 | 145 | mmol/L |
| Potassium | 3.3 | 3.2 | 3.2 | 3.6 | 3.3 | 3.3 | 3.4 | 3.3 | mmol/L |
| Chloride | 104 | 105 | 104 | 108 | 107 | 104 | 106 | 108 | mmol/L |
| LIP | N | N | N | N | N | N | N | N | |
| ICH | N | N | N | N | N | N | N | N | |
| HEM | N | N | N | N | N | N | N | N | |
| Globulin | 2.064 | 1.968 | 1.72 | 2.34 | 2.242 | 2.399 | 2.051 | 1.812 | g/dL |
| Alb/Glob Ratio | 2 | 2 | 2 | 2 | 2 | 2 | 2 | 2 | |

**Table S2 (Continued). Blood chemistry panel for NHPs over the course of the PPCN biocompatibility study.** (n = 4; 2 female (F), 2 male (M)); ALT = alanine transaminase; AST = aspartate aminotransferase; BUN = blood urea nitrogen; GGT = gamma-glutamyl transferase; I = inorganic; LIP = lipemia; ICH = icterus; HEM = hemolysis.

| **Monkey:** | **RH9142 (F)** | | | | | | | | |
|---|---|---|---|---|---|---|---|---|---|
| **Condition:** | Day of Tx | | | | | | | Day of Retrieval | **Units** |
| **Date:** | 5/18/2020 | 5/26/2020 | 6/1/2020 | 6/8/2020 | 6/15/2020 | 7/13/2020 | 8/10/2020 | 8/18/2020 | |
| **Day:** | 0 | 8 | 14 | 21 | 28 | 56 | 84 | 92 | |
| Albumin | 4.04 | 3.98 | 4.148 | 3.987 | 4.007 | 4.449 | 4.063 | 3.773 | g/dL |
| Alkaline Phosphatase | 364 | 391 | 377 | 359 | 306 | 248 | 213 | 249 | U/L |
| ALT | 19 | 20 | 25 | 28 | 20 | 24 | 39 | 25 | U/L |
| AST | 22 | 22 | 23 | 24 | 21 | 21 | 28 | 25 | U/L |
| Urea Nitrogen (BUN) | 18 | 15 | 18 | 14 | 17 | 20 | 16 | 14 | mg/dL |
| Calcium | 8.38 | 8.78 | 8.82 | 8.84 | 9.24 | 8.97 | 8.83 | 8.06 | mg/dL |
| Creatinine | 0.5171 | 0.5155 | 0.4403 | 0.4017 | 0.4636 | 0.4523 | 0.4421 | 0.4481 | mg/dL |
| GGT | 53 | 53 | 54 | 49 | 47 | 46 | 46 | 40 | U/L |
| Glucose | 61 | 64 | 79 | 65 | 75 | 58 | 55 | 63 | mg/dL |
| I. Phosphorous | 5.7 | 5.5 | 5.9 | 6.2 | 4.6 | 5.4 | 5 | 4.6 | mg/dL |
| Total Bilirubin | 0.24 | 0.17 | 0.18 | 0.15 | 0.15 | 0.2 | 0.15 | 0.3 | mg/dL |
| Total Protein | 5.81 | 6.054 | 5.963 | 5.915 | 5.889 | 6.745 | 6.17 | 5.553 | g/dL |
| Sodium | 145 | 143 | 142 | 144 | 144 | 145 | 145 | 145 | mmol/L |
| Potassium | 3.2 | 3 | 3.2 | 3.3 | 3.4 | 3.3 | 3.4 | 3.1 | mmol/L |
| Chloride | 107 | 104 | 104 | 106 | 108 | 106 | 108 | 111 | mmol/L |
| LIP | N | N | N | N | N | N | N | N | |
| ICH | N | N | N | N | N | N | N | N | |
| HEM | N | N | N | N | N | N | N | N | |
| Globulin | 1.77 | 2.074 | 1.815 | 1.928 | 1.862 | 2.296 | 2.107 | 1.78 | g/dL |
| Alb/Glob Ratio | 2 | 2 | 2 | 2 | 2 | 2 | 2 | 2 | |

**Table S2 (Continued). Blood chemistry panel for NHPs over the course of the PPCN biocompatibility study.** (n = 4; 2 female (F), 2 male (M)); ALT = alanine transaminase; AST = aspartate aminotransferase; BUN = blood urea nitrogen; GGT = gamma-glutamyl transferase; I = inorganic; LIP = lipemia; ICH = icterus; HEM = hemolysis.

| **Monkey:** | **RH9145 (M)** | | | | | | | | |
|---|---|---|---|---|---|---|---|---|---|
| **Condition:** | Day of Tx | | | | | | | Day of Retrieval | **Units** |
| **Date:** | 5/18/2020 | 5/26/2020 | 6/1/2020 | 6/8/2020 | 6/15/2020 | 7/13/2020 | 8/10/2020 | 8/18/2020 | |
| **Day:** | 0 | 8 | 14 | 21 | 28 | 56 | 84 | 92 | |
| Albumin | 4.106 | 4.142 | 4.127 | 4.211 | 4.33 | 4.3 | 4.082 | 3.965 | g/dL |
| Alkaline Phosphatase | 442 | 403 | 366 | 428 | 428 | 423 | 356 | 419 | U/L |
| ALT | 31 | 31 | 36 | 35 | 40 | 36 | 35 | 38 | U/L |
| AST | 31 | 30 | 25 | 29 | 34 | 28 | 28 | 26 | U/L |
| Urea Nitrogen (BUN) | 20 | 21 | 26 | 21 | 21 | 21 | 20 | 14 | mg/dL |
| Calcium | 9.17 | 9.35 | 9.32 | 9.19 | 9.81 | 9.11 | 8.97 | 8.95 | mg/dL |
| Creatinine | 0.5171 | 0.5939 | 0.4557 | 0.4485 | 0.5487 | 0.4367 | 0.4794 | 0.4481 | mg/dL |
| GGT | 69 | 70 | 65 | 67 | 70 | 77 | 68 | 66 | U/L |
| Glucose | 56 | 91 | 83 | 77 | 80 | 72 | 95 | 61 | mg/dL |
| I. Phosphorous | 6.3 | 5.3 | 5.1 | 5.6 | 5.2 | 5.8 | 5.7 | 6 | mg/dL |
| Total Bilirubin | 0.22 | 0.14 | 0.16 | 0.18 | 0.19 | 0.29 | 0.2 | 0.22 | mg/dL |
| Total Protein | 6.065 | 6.267 | 6.222 | 6.818 | 6.712 | 6.482 | 6.165 | 5.98 | g/dL |
| Sodium | 145 | 144 | 141 | 144 | 144 | 144 | 143 | 143 | mmol/L |
| Potassium | 3.5 | 3.3 | 3.2 | 3.3 | 3.3 | 3.3 | 3.7 | 3.3 | mmol/L |
| Chloride | 104 | 102 | 102 | 105 | 103 | 105 | 106 | 105 | mmol/L |
| LIP | N | N | N | N | N | N | N | N | |
| ICH | N | N | N | N | N | N | N | N | |
| HEM | N | N | N | N | N | N | N | N | |
| Globulin | 2.049 | 2.125 | 2.095 | 2.607 | 2.382 | 2.182 | 2.083 | 2.015 | g/dL |
| Alb/Glob Ratio | 2 | 2 | 2 | 2 | 2 | 2 | 2 | 2 | |

**Table S2 (Continued). Blood chemistry panel for NHPs over the course of the PPCN biocompatibility study.** (n = 4; 2 female (F), 2 male (M)); ALT = alanine transaminase; AST = aspartate aminotransferase; BUN = blood urea nitrogen; GGT = gamma-glutamyl transferase; I = inorganic; LIP = lipemia; ICH = icterus; HEM = hemolysis.

| **Monkey:** | **RH9148 (M)** | | | | | | | | |
|---|---|---|---|---|---|---|---|---|---|
| **Condition:** | Day of Tx | | | | | | | Day of Retrieval | |
| **Date:** | 5/18/2020 | 5/26/2020 | 6/1/2020 | 6/8/2020 | 6/15/2020 | 7/13/2020 | 8/10/2020 | 8/18/2020 | |
| **Day:** | 0 | 8 | 14 | 21 | 28 | 56 | 84 | 92 | **Units** |
| Albumin | 3.833 | 3.8 | 4.041 | 4.015 | 3.918 | 3.952 | 3.876 | 3.886 | g/dL |
| Alkaline Phosphatase | 723 | 391 | 671 | 759 | 664 | 681 | 598 | 704 | U/L |
| ALT | 30 | 35 | 27 | 34 | 29 | 28 | 30 | 38 | U/L |
| AST | 35 | 26 | 39 | 34 | 34 | 31 | 34 | 49 | U/L |
| Urea Nitrogen (BUN) | 25 | 17 | 28 | 25 | 31 | 25 | 26 | 24 | mg/dL |
| Calcium | 9.21 | 9.27 | 9.16 | 9.15 | 9.16 | 9.01 | 8.89 | 9.11 | mg/dL |
| Creatinine | 0.5017 | 0.539 | 0.4635 | 0.3939 | 0.5487 | 0.4289 | 0.5389 | 0.6108 | mg/dL |
| GGT | 63 | 74 | 60 | 62 | 58 | 62 | 60 | 59 | U/L |
| Glucose | 61 | 61 | 65 | 60 | 63 | 67 | 65 | 78 | mg/dL |
| I. Phosphorous | 7.3 | 6 | 6.9 | 6.4 | 6.2 | 6.5 | 6.4 | 6.8 | mg/dL |
| Total Bilirubin | 0.25 | 0.16 | 0.2 | 0.26 | 0.18 | 0.27 | 0.18 | 0.39 | mg/dL |
| Total Protein | 5.8 | 6.33 | 5.878 | 6.157 | 5.808 | 5.829 | 5.829 | 5.749 | g/dL |
| Sodium | 143 | 145 | 142 | 144 | 143 | 144 | 144 | 146 | mmol/L |
| Potassium | 3.4 | 3.7 | 3.4 | 3.4 | 3.4 | 3.5 | 3.4 | 3.4 | mmol/L |
| Chloride | 103 | 106 | 103 | 106 | 103 | 106 | 105 | 105 | mmol/L |
| LIP | N | N | N | N | N | N | N | N | |
| ICH | N | N | N | N | N | N | N | N | |
| HEM | N | N | N | N | N | N | N | N | |
| Globulin | 1.967 | 2.53 | 1.837 | 2.142 | 1.89 | 1.877 | 1.953 | 1.863 | g/dL |
| Alb/Glob Ratio | 2 | 2 | 2 | 2 | 2 | 2 | 2 | 2 | |

**Table S3. Urinalysis for NHPs over the course of the PPCN biocompatibility study.** (n = 4; 2 female (F), 2 male (M)); RBC = red blood cells; WBC = white blood cells.

| | **Monkey:** | **RH9138 (F)** | | | | | | | |
|---|---|---|---|---|---|---|---|---|---|
| | **Condition:** | Day of Tx | | | | | | | Day of Retrieval |
| | **Date:** | 5/18/2020 | 5/26/2020 | 6/1/2020 | 6/8/2020 | 6/15/2020 | 7/13/2020 | 8/10/2020 | 8/18/2020 |
| | **Day:** | 0 | 8 | 14 | 21 | 28 | 56 | 84 | 92 |
| *Physical* | **Appearance** | Clear | Clear | Clear | Hazy | Clear | Clear | Clear | Clear |
| | **Specific Gravity** | 1.025 | 1.005 | 1.021 | 1.025 | 1.026 | 1.02 | 1.022 | 1.014 |
| | **Color** | Yellow | Light Yellow | Yellow | Light Yellow | Yellow | Yellow | Yellow | Yellow |
| *Dipstick Evaluation* | **Leukocytes** | Negative | Negative | Negative | Negative | Negative | Negative | Negative | Negative |
| | **Nitrite** | Negative | Negative | Negative | Negative | Negative | Negative | Negative | Negative |
| | **pH** | 7 | 7 | 6 | 6 | 5 | 8 | 8 | 7 |
| | **Protein** | Trace | Negative | Trace | Trace | Trace | Trace | Trace | + (30) |
| | **Glucose** | Normal | Normal | Normal | Normal | Normal | Normal | Normal | Normal |
| | **Ketones** | Negative | Negative | Negative | Negative | Negative | Negative | Negative | Negative |
| | **Urobilinogen** | Normal | Normal | Normal | Normal | Normal | Normal | Normal | Normal |
| | **Bilirubin** | Negative | Negative | Negative | Negative | Negative | Negative | Negative | Negative |
| | **Blood** | Negative | Trace | Negative | Trace | Trace | Negative | Negative | Trace |
| *Sediment Evaluation* | **Cast Type** | - | - | - | - | None seen | None seen | None seen | - |
| | **Cast Average/10x Field** | 0 | 0 | 0 | 0 | - | - | - | 0 |
| | **Cast Type** | - | - | - | - | - | - | - | - |
| | **Cast Average/10x Field** | - | - | - | 0 | - | - | - | 0 |
| | **RBC's Average/45x Field** | 0 | 0-2 | 0 | 3-6 | 0-1 | 0-1 | 0 | 1-6 |
| | **WBC's Average/45x Field** | 0-1 | 0 | 0 | Rare | 0 | 0 | 0 | 0 |
| | **Epi cells Type** | Squamous | Squamous | Squamous | Squamous | None seen | Squamous | Transitional | Squamous |
| | **Epi cells Average/45x Field** | 0-1 | 0-1 | 0-2 | 3-large clumps | - | 0-2 | 0-1 | 0-3 |
| | **Epi cells Type** | - | Transitional | - | Transitional | - | - | - | - |
| | **Epi cells Average/10x Field** | - | 2.0-3.0 | - | 0-large clumps | - | - | - | 0 |
| | **Crysal Type** | - | - | - | - | - | - | None seen | Amorphous debri |
| | **Crystal Severity/45x Field** | 0 | 0 | 0 | 0 | 0 | 0 | - | 2+ |
| | **Crystal Type** | - | - | - | - | - | - | - | - |
| | **Crystal Severity/45x Field** | - | - | 0 | 0 | - | - | - | 0 |
| | **Bacteria Type** | - | - | - | Cocci-motile | - | - | - | Cocci |
| | **Bacteria Severity 45x Field** | 0 | 0 | 1+ | 1+ | 0 | 0 | 2+ | 0 - Rare |
| | **Sperm Severity 45x Field** | - | 0 | 0 | 0 | 0 | 0 | 0 | 0 |
| | **Mucus Severity 45x Field** | - | 0 | 0 | 0 | 0 | 0 | 0 | 0 |
| | **Yeast Severity 45x Field** | - | 0 | 0 | 0 | 0 | 0 | 0 | 0 |

**Table S3 (Continued). Urinalysis for NHPs over the course of the PPCN biocompatibility study.**
(n = 4; 2 female (F), 2 male (M)); RBC = red blood cells; WBC = white blood cells.

| | Monkey: | RH9142 (F) | | | | | | | |
|---|---|---|---|---|---|---|---|---|---|
| | Condition: | Day of Tx | | | | | | | Day of Retrieval |
| | Date: | 5/18/2020 | 5/26/2020 | 6/1/2020 | 6/8/2020 | 6/15/2020 | 7/13/2020 | 8/10/2020 | 8/18/2020 |
| | Day: | 0 | 8 | 14 | 21 | 28 | 56 | 84 | 92 |
| *Physical* | Appearance | Clear | Clear | Clear | Clear | Hazy | Clear | Clear | Clear |
| | Specific Gravity | 1.016 | 1.01 | 1.005 | 1.004 | 1.008 | 1.011 | 1.005 | 1.007 |
| | Color | Light Yellow | Yellow | Light Yellow | Pale Yellow | Light Yellow | Light Yellow | Light Yellow | Light Yellow |
| *Dipstick Evaluation* | Leukocytes | Negative | Negative | Negative | Negative | Negative | Negative | Negative | Negative |
| | Nitrite | Negative | Negative | Negative | Positive | Negative | Negative | Negative | Negative |
| | pH | 7 | 8 | 8 | 8 | 5 | 7 | 7 | 5 |
| | Protein | Trace | Negative | Negative | Trace | Negative | Negative | Trace | Trace |
| | Glucose | Normal | Normal | Normal | Normal | Normal | Normal | Normal | Normal |
| | Ketones | Negative | Negative | Negative | Negative | Negative | Negative | Negative | Negative |
| | Urobilinogen | Normal | Normal | Normal | Normal | Normal | Normal | Normal | Normal |
| | Bilirubin | Negative | Negative | Negative | Negative | Negative | Negative | Negative | Negative |
| | Blood | Trace | Trace | 250 Ery/µL | Trace | Trace | Trace | Negative | Trace |
| *Sediment Evaluation* | Cast Type | - | - | - | - | None seen | None seen | None seen | - |
| | Cast Average/10x Field | 0 | 0 | 0 | 0 | - | - | - | 0 |
| | Cast Type | - | - | - | - | - | - | - | - |
| | Cast Average/10x Field | - | - | - | 0 | - | - | - | 0 |
| | RBC’s Average/45x Field | 0-1 | 0 | 0 | 0-1 | 0 | 1-2 | 0 | 0-2 |
| | WBC’s Average/45x Field | 0 | 0 | 0 | 0 | 0 | 0 | 0 | 0 |
| | Epi cells Type | Squamous | Squamous | Squamous | Squamous | None seen | Squamous | - | Squamous |
| | Epi cells Average/45x Field | 0-1 | 0-1 | 0-1 | 1-3 | - | 0-1 | 0 | 0-2 |
| | Epi cells Type | - | - | Transitional | - | - | - | - | - |
| | Epi cells Average/10x Field | - | - | 0-1 | 0 | - | - | - | 0 |
| | Crysal Type | - | - | - | Amorphous | None seen | - | None seen | - |
| | Crystal Severity/45x Field | 0 | 0 | 0 | 1+ | - | 0 | - | 0 |
| | Crystal Type | - | - | - | - | - | - | - | - |
| | Crystal Severity/45x Field | - | - | - | 0 | - | - | - | 0 |
| | Bacteria Type | - | - | - | cocci-motile (clostridium) | - | - | - | Cocci-motile |
| | Bacteria Severity 45x Field | 1+ | 0 | 1+ | 1+ | 0 | 1+ | 1+ | 1+ |
| | Sperm Severity 45x Field | 0 | 0 | - | 0 | 0 | 0 | 0 | 0 |
| | Mucus Severity 45x Field | 0 | 0 | - | 0 | 0 | 0 | 0 | 0 |
| | Yeast Severity 45x Field | 0 | 0 | - | 0 | 0 | 0 | 0 | 0 |

**Table S3 (Continued). Urinalysis for NHPs over the course of the PPCN biocompatibility study.**
(n = 4; 2 female (F), 2 male (M)); RBC = red blood cells; WBC = white blood cells.

| | Monkey: | RH9145 (M) | | | | | | | |
|---|---|---|---|---|---|---|---|---|---|
| | Condition: | Day of Tx | | | | | | | Day of Retrieval |
| | Date: | 5/18/2020 | 5/26/2020 | 6/1/2020 | 6/8/2020 | 6/15/2020 | 7/13/2020 | 8/10/2020 | 8/18/2020 |
| | Day: | 0 | 8 | 14 | 21 | 28 | 56 | 84 | 92 |
| *Physical* | Appearance | Clear | Clear | Clear | Cloudy | Clear | Clear | - | Clear |
| | Specific Gravity | 1.026 | 1.006 | 1.02 | 1.026 | 1.002 | 1.007 | 1.013 | 1.032 |
| | Color | Yellow | Colorless | Light Yellow | Yellow | Colorless | Colorless | Yellow | Yellow |
| *Dipstick Evaluation* | Leukocytes | Negative | Negative | Negative | Negative | Negative | Negative | Negative | Negative |
| | Nitrite | Negative | Negative | Negative | Negative | Negative | Negative | Negative | Negative |
| | pH | 7 | 8 | 8 | 9 | 7 | 8 | 8 | 6 |
| | Protein | Trace | Negative | Trace | + (30) | Trace | Trace | Trace | + (30) |
| | Glucose | Normal | Normal | Normal | Normal | Normal | Normal | Normal | Normal |
| | Ketones | Negative | Negative | Negative | Negative | Negative | Negative | Negative | + (Small) |
| | Urobilinogen | Normal | Normal | Normal | Normal | Normal | Normal | Normal | Normal |
| | Bilirubin | Negative | Negative | Negative | Negative | Negative | Negative | Negative | Negative |
| | Blood | Negative | Negative | 50 | 250 Ery/µL | 250 Ery/µL | Trace | Trace | Negative |
| *Sediment Evaluation* | Cast Type | - | - | - | - | None seen | None seen | None seen | - |
| | Cast Average/10x Field | 0 | 0 | 0 | 0 | - | - | - | 0 |
| | Cast Type | - | - | - | - | - | - | - | - |
| | Cast Average/10x Field | - | - | - | 0 | - | - | - | 0 |
| | RBC's Average/45x Field | 0 | 0 | 0 | >50 | 0-1 | 0 | 0-1 | 2-7 |
| | WBC's Average/45x Field | 0 | 0 | 0 | 0 | 0 | 0 | 0 | 0 |
| | Epi cells Type | - | - | - | Transitional | None seen | Squamous | None seen | - |
| | Epi cells Average/45x Field | 0 | - | 0 | 0 -small clumps | - | 1-2 | - | 0 |
| | Epi cells Type | - | - | - | - | - | Transitional | - | - |
| | Epi cells Average/10x Field | - | - | - | 0 | - | 0-1 | - | 0 |
| | Crysal Type | - | - | - | - | None seen | - | None seen | - |
| | Crystal Severity/45x Field | 0 | 0 | 0 | 0 | - | 0 | 0 | 0 |
| | Crystal Type | - | - | - | - | - | - | - | - |
| | Crystal Severity/45x Field | - | - | - | 0 | - | - | - | 0 |
| | Bacteria Type | - | - | - | Cocci-motile | - | - | - | Cocci-motile |
| | Bacteria Severity 45x Field | 1+ | 0 | 1+ | Rare | 0 | 1+ | 2+ | 2+ |
| | Sperm Severity 45x Field | 0 | 0 | 0 | 0 | 0 | 0 | 0 | 0 |
| | Mucus Severity 45x Field | 0 | 0 | 0 | 0 | 0 | 0 | 0 | 0 |
| | Yeast Severity 45x Field | 0 | 0 | 0 | 0 | 0 | 0 | 0 | 0 |

**Table S3 (Continued). Urinalysis for NHPs over the course of the PPCN biocompatibility study.** (n = 4; 2 female (F), 2 male (M)); RBC = red blood cells; WBC = white blood cells.

| | **Monkey:** | **RH9148 (M)** | | | | | | | |
|---|---|---|---|---|---|---|---|---|---|
| | **Condition:** | Day of Tx | | | | | | | Day of Retrieval |
| | **Date:** | 5/18/2020 | 5/26/2020 | 6/1/2020 | 6/8/2020 | 6/15/2020 | 7/13/2020 | 8/10/2020 | 8/18/2020 |
| | **Day:** | 0 | 8 | 14 | 21 | 28 | 56 | 84 | 92 |
| *Physical* | **Appearance** | Clear | Clear | Clear | Hazy | Clear | Clear | Hazy | Clear |
| | **Specific Gravity** | 1.033 | 1.026 | 1.03 | 1.034 | 1.017 | 1.025 | 1.027 | 1.026 |
| | **Color** | Yellow | Yellow | Yellow | Yellow | Light Yellow | Yellow | Yellow | Yellow |
| *Dipstick Evaluation* | **Leukocytes** | Negative | Negative | Negative | Negative | Negative | Negative | Negative | Negative |
| | **Nitrite** | Negative | Negative | Negative | Negative | Negative | Negative | Negative | Negative |
| | **pH** | 6 | 7 | 8 | 6 | 7 | 7 | 6 | 5 |
| | **Protein** | Trace | Trace | Trace | Trace | Negative | Trace | Trace | Trace |
| | **Glucose** | Normal | Normal | Normal | Normal | Normal | Normal | Normal | Normal |
| | **Ketones** | Negative | Negative | Negative | Negative | Negative | Negative | Negative | + (Small) |
| | **Urobilinogen** | Normal | Normal | Normal | Normal | Normal | Normal | Normal | Normal |
| | **Bilirubin** | Negative | Negative | Negative | Negative | Negative | Negative | Negative | Negative |
| | **Blood** | Negative | Negative | Trace | Negative | Trace | Trace | Negative | Negative |
| *Sediment Evaluation* | **Cast Type** | - | - | - | - | None seen | None seen | None seen | - |
| | **Cast Average/10x Field** | 0 | 0 | 0 | 0 | - | - | - | 0 |
| | **Cast Type** | - | - | - | - | - | - | - | - |
| | **Cast Average/10x Field** | - | - | - | 0 | - | - | - | 0 |
| | **RBC's Average/45x Field** | 0 | 0 | 0 | 0 | 0-1 | 0-1 | 0 | 0-3 |
| | **WBC's Average/45x Field** | 0 | 0 | 0 | 0 | 0 | 0 | 0 | 0-1 |
| | **Epi cells Type** | Squamous | Squamous | - | Transitional | None seen | None seen | None seen | Transitional |
| | **Epi cells Average/45x Field** | 0-2 | 0-1 | 0 | 0- large clumps | - | - | - | 0-3 |
| | **Epi cells Type** | - | - | - | Squamous | - | - | - | Squamous |
| | **Epi cells Average/10x Field** | - | - | - | Rare | - | - | - | 0-4 |
| | **Crysal Type** | - | - | - | - | None seen | None seen | None seen | - |
| | **Crystal Severity/45x Field** | 0 | 0 | 0 | 0 | - | - | - | 0 |
| | **Crystal Type** | - | - | - | - | - | - | - | - |
| | **Crystal Severity/45x Field** | - | - | 0 | 0 | - | - | - | 0 |
| | **Bacteria Type** | - | - | - | Cocci-motile | - | - | - | Cocci-motile |
| | **Bacteria Severity 45x Field** | 0 | 0 | 1+ | 3+ | 0 | 1+ | 2+ | 1+ |
| | **Sperm Severity 45x Field** | 0 | 0 | 0 | 0 | 0 | 0 | 0 | - |
| | **Mucus Severity 45x Field** | 0 | 0 | 0 | 0 | 0 | 0 | 0 | - |
| | **Yeast Severity 45x Field** | 0 | 0 | 0 | 0 | 0 | 0 | 0 | - |

**Table S4. Body temperature of NHPs over the course of the PPCN biocompatibility study.**
(n = 4; 2 female (F), 2 male (M)).

| **Body Temperature (°F)** | | | | | | | |
|---|---|---|---|---|---|---|---|
| **Condition:** | Day of Tx | | | | | | Day of Retrieval |
| **Date:** | 5/18/2020 | 5/26/2020 | 6/1/2020 | 6/8/2020 | 6/15/2020 | 7/13/2020 | 8/18/2020 |
| **Day:** | 0 | 8 | 14 | 21 | 28 | 56 | 92 |
| **RH9138** | 100.7 | 99.9 | 101.1 | 101.4 | 101.4 | 102.2 | 100.4 |
| **RH9142** | 100.2 | 101.4 | 101.4 | 100.6 | 101.6 | 101.3 | 99.6 |
| **RH9145** | 98.8 | 102.1 | 101.1 | 101.5 | 101.7 | 102.3 | 99.4 |
| **RH9148** | 98.3 | 102.4 | 101.3 | 101.6 | 101.3 | 101.3 | 100.1 |

**Table S5. Complete blood counts for NHPs over the course of the TP-IAT to the omentum with PPCN study.** (n = 2; 1 female (F), 1 male (M)); WBC = white blood cells; RBC = red blood cells; HGB = hemoglobin; HCT = hematocrit; MCV = mean corpuscular volume; MCH = mean corpuscular hemoglobin; MCHC = mean corpuscular hemoglobin concentration; PLT = platelets; MPV = mean platelet volume; RDW = red cell distribution width; NEUT = neutrophils; LYMPH = lymphocytes; MONO = monocytes; EOS = eosinophils; BASO = basophils; LUC = large unstained cells; RETIC =reticulocytes; MACRO = macrocytosis; HC-VAR = hemoglobin concentration variance; PLATCLMP = platelet clumps; HYPER = hyperchromia.

| Monkey: | RH9139 (F) | | | | | | | | | |
|---|---|---|---|---|---|---|---|---|---|---|
| Condition: | Baseline | IVDTT Baseline | Pre-Transplant | Day of Transplant | Post-Transplant | | | | | |
| Date: | 8/4/2022 | 8/9/2022 | 8/15/2022 | 8/16/2022 | 8/19/2022 | 8/23/2022 | 8/30/2022 | 9/6/2022 | 9/13/2022 | |
| Day: | -12 | -7 | -1 | 0 | 3 | 7 | 14 | 21 | 28 | Units |
| WBC | 6.03 | 7.23 | 6.08 | 11.35 | 9.83 | 11.04 | 8.78 | 7.95 | 7.69 | x10^3/µL |
| RBC | 4.83 | 4.50 | 4.24 | 4.35 | 4.24 | 4.18 | 4.27 | 4.55 | 4.43 | x10^6/µL |
| HGB | 11.7 | 11.2 | 10.4 | 10.6 | 10.1 | 10.1 | 10.5 | 11.0 | 10.8 | g/dL |
| HCT | 38.3 | 35.5 | 33.0 | 34.4 | 33.4 | 33.8 | 33.4 | 36.0 | 35.4 | % |
| MCV | 79.4 | 78.9 | 77.8 | 79.0 | 78.9 | 80.8 | 78.2 | 79.1 | 79.9 | fL |
| MCH | 24.3 | 24.9 | 24.6 | 24.5 | 23.9 | 24.1 | 24.5 | 24.2 | 24.3 | pg |
| MCHC | 30.6 | 31.6 | 31.6 | 31 | 30.3 | 29.8 | 31.3 | 30.6 | 30.4 | g/dL |
| PLT | 291 | 264 | 334 | 340 | 372 | 576 | 369 | 313 | 335 | x10^3/µL |
| MPV | 7.3 | 8.1 | 7.2 | 7.1 | 8.8 | 7.8 | 8.0 | 8.9 | 8.3 | fL |
| RDW | 12.9 | 12.6 | 12.6 | 12.7 | 12.8 | 13.0 | 12.9 | 13.1 | 14.0 | % |
| %NEUT | 31.6 | 56.2** | 40.6 | 80.5***** | 63.0***** | 62.1***** | 48.2 | 49.6 | 40.4 | % |
| %LYMPH | 61.4 | 39.3 | 53.5 | 12.1*** | 30.1 | 29.1 | 41.4 | 43.9 | 52.9 | % |
| %MONO | 2.9 | 1.2 | 0.9** | 1.6 | 3.8 | 1.2 | 1.8 | 3.7 | 1.9 | % |
| %EOS | 3.0 | 2.4 | 4.0 | 5.3 | 2.1 | 6.6*** | 7.5** | 1.4 | 3.4 | % |
| %BASO | 0.2 | 0.4 | 0.1 | 0.1 | 0.1 | 0.4 | 0.2 | 0.3 | 0.4 | % |
| %LUC | 0.9 | 0.4 | 0.9 | 0.4 | 0.9 | 0.7 | 0.9 | 1.1 | 1 | % |
| #NEUT | 1.91 | 4.07 | 2.47 | 9.13***** | 6.19 | 6.85***** | 4.23 | 3.94 | 3.11 | x10^3/µL |
| #LYMPH | 3.70 | 2.84 | 3.25 | 1.37*** | 2.96 | 3.21 | 3.63 | 3.49 | 4.07 | x10^3/µL |
| #MONO | 0.17 | 0.09 | 0.05 | 0.18 | 0.38 | 0.13 | 0.16 | 0.29 | 0.15 | x10^3/µL |
| #EOS | 0.18 | 0.17 | 0.24 | 0.60*** | 0.21 | 0.72*** | 0.66** | 0.12 | 0.26 | x10^3/µL |
| #BASO | 0.01 | 0.03 | 0.01 | 0.01 | 0.01 | 0.04 | 0.02 | 0.02 | 0.03 | x10^3/µL |
| #LUC | 0.06 | 0.03 | 0.06 | 0.05 | 0.08 | 0.08 | 0.08 | 0.08 | 0.08 | x10^3/µL |
| %RETIC | 1.21 | 1.67 | 2.03 | 2.02 | 2.68 | 2.61 | 3.1 | 3.3 | 2.72 | % |
| #RETIC | 58.4 | 75.0**** | 86.2 | 87.8 | 113.5 | 109.1 | 132.1 | 150.2 | 120.8 | x10^9/µL |

* Elevated/Low Levels, Comparable to baseline
** Elevated/Low Levels, Not of clinical concern
***Elevated/Low Levels likely due to stress response associated with surgery
****Reticulocytes not considered significant unless animal anemic; RBCs normal
*****Neutrophils: elevated due to surgical inflammation
*.* Elevated/ Low Levels possibly due to Immunosuppression/implanted biomaterial/encapsulated cells/cell products
*_* Elevated levels possibly due to neutrophils being counted as eosinophils

**Table S5 (Continued). Complete blood counts for NHPs over the course of the TP-IAT to the omentum with PPCN study.**

(n = 2; 1 female (F), 1 male (M)); WBC = white blood cells; RBC = red blood cells; HGB = hemoglobin; HCT = hematocrit; MCV = mean corpuscular volume; MCH = mean corpuscular hemoglobin; MCHC = mean corpuscular hemoglobin concentration; PLT = platelets; MPV = mean platelet volume; RDW = red cell distribution width; NEUT = neutrophils; LYMPH = lymphocytes; MONO = monocytes; EOS = eosinophils; BASO = basophils; LUC = large unstained cells; RETIC =reticulocytes; MACRO = macrocytosis; HC-VAR = hemoglobin concentration variance; PLATCLMP = platelet clumps; HYPER = hyperchromia.

| Monkey: | RH9139 (F) | | | | | | | | |
|---|---|---|---|---|---|---|---|---|---|
| Condition: | IVDTT Post-Transplant | Glucagon Post-Transplant | Post-Transplant | | IVDTT Post-Transplant | Post-Transplant | | | |
| Date: | 9/15/2022 | 9/20/2022 | 9/27/2022 | 10/4/2022 | 10/10/2022 | 10/18/2022 | 10/25/2022 | 11/1/2022 | |
| Day: | 30 | 35 | 42 | 49 | 55 | 63 | 70 | 77 | Units |
| WBC | 9.18 | 8.01 | 10.26 | 8.75 | 6.59 | 13.02 | 13.6 | 11.16 | x10^3/µL |
| RBC | 4.39 | 4.57 | 4.57 | 4.59 | 4.47 | 4.27 | 4.56 | 4.46 | x10^6/µL |
| HGB | 10.9 | 11.0 | 11.3 | 11.1 | 11.0 | 10.5 | 11.2 | 11.1 | g/dL |
| HCT | 34.6 | 36.0 | 36.6 | 36.3 | 35.4 | 34.6 | 35.8 | 35.3 | % |
| MCV | 79.0 | 78.8 | 80.0 | 79.1 | 79.3 | 81.0 | 78.5 | 79.0 | fL |
| MCH | 24.8 | 24.1 | 24.7 | 24.2 | 24.6 | 24.7 | 24.4 | 24.9 | pg |
| MCHC | 31.4 | 30.6 | 30.9 | 30.6 | 31 | 30.4 | 31.1 | 31.5 | g/dL |
| PLT | 332 | 377 | 406 | 363 | 347 | 427 | 429 | 362 | x10^3/µL |
| MPV | 8.1 | 8.1 | 8.1 | 8.0 | 7.7 | 8.1 | 7.8 | 7.9 | fL |
| RDW | 14.2 | 14.3 | 13.8 | 14.2 | 14.2 | 14.1 | 14.5 | 14.4 | % |
| %NEUT | 51.7 | 50.3 | 37.9 | 46.7 | 74.2** | 68.7** | 68.1** | 62.4** | % |
| %LYMPH | 43.4 | 42.4 | 52.8 | 47.5 | 23** | 27.5 | 29.0 | 32.1 | % |
| %MONO | 1.4 | 3.3 | 2.3 | 2.5 | 1.8 | 1.9 | 2.1 | 1.9 | % |
| %EOS | 2.3 | 2.7 | 5.7 | 1.9 | 0.2 | 1.1 | 0.3 | 2.6 | % |
| %BASO | 0.2 | 0.3 | 0.3 | 0.4 | 0 | 0.2 | 0.2 | 0.2 | % |
| %LUC | 0.9 | 1 | 0.9 | 1 | 0.7 | 0.5 | 0.3 | 0.8 | % |
| #NEUT | 4.75 | 4.03 | 3.89 | 4.09 | 4.89 | 8.95** | 9.27** | 6.97** | x10^3/µL |
| #LYMPH | 3.98 | 3.4 | 5.42 | 4.16 | 1.52 | 3.59 | 3.95 | 3.59 | x10^3/µL |
| #MONO | 0.13 | 0.26 | 0.23 | 0.22 | 0.12 | 0.25 | 0.29 | 0.21 | x10^3/µL |
| #EOS | 0.21 | 0.22 | 0.58** | 0.17 | 0.02 | 0.14 | 0.04 | 0.29 | x10^3/µL |
| #BASO | 0.02 | 0.02 | 0.03 | 0.03 | 0 | 0.02 | 0.02 | 0.02 | x10^3/µL |
| #LUC | 0.08 | 0.08 | 0.09 | 0.09 | 0.04 | 0.07 | 0.04 | 0.08 | x10^3/µL |
| %RETIC | 1.98 | 2.17 | 2.37 | 2.7 | 2.13 | 2.69 | 2.13 | 2.35 | % |
| #RETIC | 86.7 | 98.9 | 108.4 | 123.8 | 94.9 | 114.6 | 97.2 | 104.7 | x10^9/µL |

* Elevated/Low Levels, Comparable to baseline
** Elevated/Low Levels, Not of clinical concern
***Elevated/Low Levels likely due to stress response associated with surgery
****Reticulocytes not considered significant unless animal anemic; RBCs normal
*****Neutrophils: elevated due to surgical inflammation
*.* Elevated/ Low Levels possibly due to Immunosuppression/implanted biomaterial/encapsulated cells/cell products
*_* Elevated levels possibly due to neutrophils being counted as eosinophils

**Table S5 (Continued). Complete blood counts for NHPs over the course of the TP-IAT to the omentum with PPCN study.**

(n = 2; 1 female (F), 1 male (M)); WBC = white blood cells; RBC = red blood cells; HGB = hemoglobin; HCT = hematocrit; MCV = mean corpuscular volume; MCH = mean corpuscular hemoglobin; MCHC = mean corpuscular hemoglobin concentration; PLT = platelets; MPV = mean platelet volume; RDW = red cell distribution width; NEUT = neutrophils; LYMPH = lymphocytes; MONO = monocytes; EOS = eosinophils; BASO = basophils; LUC = large unstained cells; RETIC =reticulocytes; MACRO = macrocytosis; HC-VAR = hemoglobin concentration variance; PLATCLMP = platelet clumps; HYPER = hyperchromia.

| Monkey: | RH9139 (F) | | | | | | | |
|---|---|---|---|---|---|---|---|---|
| Condition: | Post-Transplant | | | IVDTT Post-Transplant | Survival Omentectomy Post-Transplant | IVDTT Post - Omentectomy | Termination & Necropsy | |
| Date: | 11/7/2022 | 11/15/2022 | 11/22/2022 | 11/29/2022 | 12/2/2022 | 12/5/2022 | 12/8/2022 | |
| Day: | 83 | 91 | 98 | 105 | 108 | 111 | 114 | Units |
| WBC | 6.69 | 7.98 | 9.24 | 8.97 | 5.68 | 6.74 | 6.92 | x10^3/µL |
| RBC | 4.44 | 4.47 | 4.66 | 4.69 | 4.34 | 4.23 | 4.09 | x10^6/µL |
| HGB | 11.0 | 10.8 | 11.4 | 11.3 | 10.7 | 10.4 | 10.0 | g/dL |
| HCT | 34.3 | 35.3 | 37.0 | 37.4 | 34.2 | 34.1 | 31.6 | % |
| MCV | 77.2 | 79.0 | 79.4 | 79.6 | 78.7 | 80.8 | 77.2 | fL |
| MCH | 24.8 | 24.2 | 24.5 | 24.1 | 24.6 | 24.6 | 24.5 | pg |
| MCHC | 32.2 | 30.6 | 30.9 | 30.2 | 31.3 | 30.4 | 31.7 | g/dL |
| PLT | 388 | 393 | 376 | 386 | 356 | 359 | 397 | x10^3/µL |
| MPV | 7.9 | 8.4 | 7.9 | 8.1 | 8.0 | 8.2 | 8.5 | fL |
| RDW | 14.4 | 14.5 | 14.5 | 14.5 | 14.8 | 14.6 | 14.6 | % |
| %NEUT | 48.9 | 69.3** | 55.5** | 54.4** | 43.5 | 69.1***** | 39.8 | % |
| %LYMPH | 44.6 | 26.5 | 40.3 | 40.7 | 50.7 | 24.8*** | 50.1 | % |
| %MONO | 1.5 | 2.1 | 2.8 | 2.6 | 2.9 | 1.5 | 2.6 | % |
| %EOS | 3.9 | 1.3 | 0.4 | 1.2 | 1.2 | 3.5 | 6.4 | % |
| %BASO | 0.2 | 0 | 0.1 | 0.1 | 0.1 | 0.1 | 0.1 | % |
| %LUC | 0.9 | 0.8 | 0.9 | 1 | 1.6 | 1 | 1 | % |
| #NEUT | 3.27 | 5.53 | 5.13 | 4.88 | 2.47 | 4.66 | 2.75 | x10^3/µL |
| #LYMPH | 2.98 | 2.11 | 3.72 | 3.65 | 2.88 | 1.67 | 3.47 | x10^3/µL |
| #MONO | 0.1 | 0.17 | 0.26 | 0.23 | 0.16 | 0.1 | 0.18 | x10^3/µL |
| #EOS | 0.26 | 0.10 | 0.03 | 0.11 | 0.07 | 0.23 | 0.45 | x10^3/µL |
| #BASO | 0.02 | 0 | 0.01 | 0.01 | 0.01 | 0.01 | 0.01 | x10^3/µL |
| #LUC | 0.06 | 0.06 | 0.09 | 0.09 | 0.09 | 0.07 | 0.07 | x10^3/µL |
| %RETIC | 1.69 | 2.95 | 1.92 | 2.13 | 2.27 | 2.87 | 1.89 | % |
| #RETIC | 75.2**** | 131.9 | 89.4 | 99.8 | 98.5 | 121.4 | 77.1**** | x10^9/µL |

* Elevated/Low Levels, Comparable to baseline
** Elevated/Low Levels, Not of clinical concern
***Elevated/Low Levels likely due to stress response associated with surgery
****Reticulocytes not considered significant unless animal anemic; RBCs normal
*****Neutrophils: elevated due to surgical inflammation
*.* Elevated/ Low Levels possibly due to Immunosuppression/implanted biomaterial/encapsulated cells/cell products
*_* Elevated levels possibly due to neutrophils being counted as eosinophils

**Table S5 (Continued). Complete blood counts for NHPs over the course of the TP-IAT to the omentum with PPCN study.**

(n = 2; 1 female (F), 1 male (M)); WBC = white blood cells; RBC = red blood cells; HGB = hemoglobin; HCT = hematocrit; MCV = mean corpuscular volume; MCH = mean corpuscular hemoglobin; MCHC = mean corpuscular hemoglobin concentration; PLT = platelets; MPV = mean platelet volume; RDW = red cell distribution width; NEUT = neutrophils; LYMPH = lymphocytes; MONO = monocytes; EOS = eosinophils; BASO = basophils; LUC = large unstained cells; RETIC =reticulocytes; MACRO = macrocytosis; HC-VAR = hemoglobin concentration variance; PLATCLMP = platelet clumps; HYPER = hyperchromia.

| Monkey: | RH9144 (M) | | | | | | | | | |
|---|---|---|---|---|---|---|---|---|---|---|
| Condition: | Baseline | Pre-Transplant | Post-Transplant | | | | | | | |
| Date: | -6 | 0 | 7 | 14 | 21 | 28 | 30 | 42 | 49 | |
| Day: | 6/15/2022 | 6/20/2022 | 6/28/2022 | 7/5/2022 | 7/12/2022 | 7/19/2022 | 7/21/2022 | 8/2/2022 | 8/9/2022 | Units |
| WBC | 7.1 | 10.19 | 27.72*** | 15.31 | 10.56 | 11.39 | 8.95 | 9.72 | 12.25 | x10^3/µL |
| RBC | 5.31 | 4.80 | 4.38 | 4.49 | 4.94 | 5.22 | 5.05 | 5.11 | 5.36 | x10^6/µL |
| HGB | 12.3 | 11.1 | 10.1 | 10.4 | 11.3 | 12.1 | 11.8 | 12.1 | 12.7 | g/dL |
| HCT | 38.8 | 34.7 | 33.0 | 32.9 | 36.7 | 38.9 | 38.0 | 38.5 | 40.8 | % |
| MCV | 73.0 | 72.4 | 75.3 | 73.3 | 74.2 | 74.5 | 75.1 | 75.4 | 76.1 | fL |
| MCH | 23.2 | 23.1 | 23.1 | 23.1 | 23.0 | 23.1 | 23.3 | 23.7 | 23.6 | pg |
| MCHC | 31.8 | 32 | 30.7 | 31.6 | 30.9 | 31 | 31 | 31.4 | 31 | g/dL |
| PLT | 393 | 398 | 882*** | 813*** | 639 | 511 | 500 | 549 | 492 | x10^3/µL |
| MPV | 7.5 | 7.0 | 6.8 | 7.1 | 7.3 | 7.5 | 8.1 | 7.1 | 9.0 | fL |
| RDW | 12.2 | 12.0 | 12.9 | 12.8 | 12.8 | 12.6 | 12.7 | 12.7 | 12.5 | % |
| %NEUT | 55.5 | 63.5* | 89.2***** | 80.5***** | 60.9* | 71.9*.* | 42.1 | 49.2 | 51.6 | % |
| %LYMPH | 39.4 | 31.2 | 6.1*** | 13.3*** | 28.1 | 22.4*.* | 45.3 | 41.5 | 41.3 | % |
| %MONO | 3.7 | 2.8 | 2.4 | 2.9 | 3.1 | 2.3 | 3.9 | 3.5 | 2.3 | % |
| %EOS | 0.2 | 1.2 | 1.6 | 2.1 | 5.9 | 2.0 | 6.2 | 3.9 | 3.0 | % |
| %BASO | 0.2 | 0.2 | 0.1 | 0.3 | 0.5 | 0.1 | 0.2 | 0.3 | 0.5 | % |
| %LUC | 1 | 1.1 | 0.6 | 1 | 1.5 | 1.4 | 2.3 | 1.6 | 1.3 | % |
| #NEUT | 3.94 | 6.47** | 24.72***** | 12.32***** | 6.44*.* | 8.19*.* | 3.77 | 4.79 | 6.32*.* | x10^3/µL |
| #LYMPH | 2.80 | 3.18 | 1.69 | 2.04 | 2.97 | 2.55 | 4.05 | 4.04 | 5.06 | x10^3/µL |
| #MONO | 0.27 | 0.28 | 0.68 | 0.44 | 0.33 | 0.26 | 0.35 | 0.34 | 0.28 | x10^3/µL |
| #EOS | 0.01 | 0.13 | 0.44 | 0.32 | 0.63*.* | 0.23 | 0.56*.* | 0.37 | 0.37 | x10^3/µL |
| #BASO | 0.02 | 0.02 | 0.03 | 0.04 | 0.05 | 0.01 | 0.02 | 0.03 | 0.06 | x10^3/µL |
| #LUC | 0.07 | 0.11 | 0.17 | 0.15 | 0.16 | 0.16 | 0.2 | 0.15 | 0.16 | x10^3/µL |
| %RETIC | 1.03 | 1.25 | 3.66**** | 3.60**** | 2.77 | 2.39 | 2.53 | 2.25 | 2.16 | % |
| #RETIC | 54.4 | 59.7** | 160.6 | 161.6 | 137.1 | 124.9 | 127.9 | 115 | 115.5 | x10^9/µL |

* Elevated/Low Levels, Comparable to baseline
**Elevated/ Low Levels, Not of clinical concern
***Elevated or Low Levels likely due to stress response associated with surgery
****Reticulocytes not considered significant unless animal anemic; RBCs normal
*****Neutrophils: elevated due to surgical inflammation
*.* Elevated/ Low Levels possibly due to Immunosuppression/implanted biomaterial/encapsulated cells/cell products
*_* Elevated levels possibly due to neutrophils being counted as eosinophils

**Table S5 (Continued). Complete blood counts for NHPs over the course of the TP-IAT to the omentum with PPCN study.**

(n = 2; 1 female (F), 1 male (M)); WBC = white blood cells; RBC = red blood cells; HGB = hemoglobin; HCT = hematocrit; MCV = mean corpuscular volume; MCH = mean corpuscular hemoglobin; MCHC = mean corpuscular hemoglobin concentration; PLT = platelets; MPV = mean platelet volume; RDW = red cell distribution width; NEUT = neutrophils; LYMPH = lymphocytes; MONO = monocytes; EOS = eosinophils; BASO = basophils; LUC = large unstained cells; RETIC =reticulocytes; MACRO = macrocytosis; HC-VAR = hemoglobin concentration variance; PLATCLMP = platelet clumps; HYPER = hyperchromia.

| **Monkey:** | **RH9144** (M) | | | | | | | |
|---|---|---|---|---|---|---|---|---|
| **Condition:** | Post-Transplant | | | | | | | |
| **Date:** | 63 | 70 | 77 | 84 | 91 | 98 | 105 | |
| **Day:** | 8/23/2022 | 8/30/2022 | 9/6/2022 | 9/13/2022 | 9/20/2022 | 9/27/2022 | 10/4/2022 | **Units** |
| **WBC** | 11.64 | 11.8 | 10.91 | 12.37 | 12.69 | 11.16 | 11.65 | x10^3/µL |
| **RBC** | 5.13 | 5.39 | 5.35 | 5.40 | 5.25 | 5.39 | 5.71 | x10^6/µL |
| **HGB** | 12.1 | 12.7 | 12.4 | 12.6 | 12.2 | 12.8 | 13.6 | g/dL |
| **HCT** | 39.1 | 39.8 | 39.6 | 40.9 | 38.9 | 40.5 | 42.7 | % |
| **MCV** | 76.1 | 73.8 | 74.1 | 75.7 | 74.0 | 75.2 | 74.8 | fL |
| **MCH** | 23.5 | 23.6 | 23.3 | 23.3 | 23.3 | 23.8 | 23.9 | pg |
| **MCHC** | 30.8 | 32 | 31.4 | 30.8 | 31.4 | 31.7 | 31.9 | g/dL |
| **PLT** | 492 | 503 | 493 | 460 | 462 | 473 | 535 | x10^3/µL |
| **MPV** | 8.0 | 8.3 | 9.0 | 8.8 | 7.9 | 9.4 | 9.1 | fL |
| **RDW** | 12.7 | 12.6 | 12.7 | 14.3 | 14.5 | 14.1 | 14.6 | % |
| **%NEUT** | 48.1 | 44.4 | 44.8 | 58.6* | 55.9* | 44.4 | 44 | % |
| **%LYMPH** | 45.5 | 48.5 | 48.0 | 34.8 | 36.1 | 45.1 | 47.3 | % |
| **%MONO** | 2.0 | 3.0 | 2.5 | 2.0 | 2.7 | 3.7 | 3.4 | % |
| **%EOS** | 2.8 | 2.8 | 2.7 | 2.9 | 4.2 | 5.2 | 3.0 | % |
| **%BASO** | 0.4 | 0.2 | 0.4 | 0.4 | 0.2 | 0.2 | 0.5 | % |
| **%LUC** | 1.2 | 1.1 | 1.5 | 1.3 | 1 | 1.4 | 1.8 | % |
| **#NEUT** | 5.59 | 5.24 | 4.89 | 7.25*.* | 7.09*.* | 4.96 | 5.13 | x10^3/µL |
| **#LYMPH** | 5.3 | 5.73 | 5.24 | 4.3 | 4.57 | 5.03 | 5.51 | x10^3/µL |
| **#MONO** | 0.23 | 0.35 | 0.28 | 0.25 | 0.34 | 0.41 | 0.4 | x10^3/µL |
| **#EOS** | 0.32 | 0.33 | 0.29 | 0.36 | 0.54** | 0.59** | 0.35 | x10^3/µL |
| **#BASO** | 0.04 | 0.03 | 0.05 | 0.05 | 0.02 | 0.02 | 0.05 | x10^3/µL |
| **#LUC** | 0.14 | 0.13 | 0.16 | 0.16 | 0.13 | 0.16 | 0.21 | x10^3/µL |
| **%RETIC** | 2.1 | 1.74 | 1.86 | 1.43 | 1.22 | 1.6 | 1.76 | % |
| **#RETIC** | 107.9 | 93.7 | 99.8 | 77.5 | 64**** | 86.4 | 100.3 | x10^9/µL |

* Elevated/Low Levels, Comparable to baseline
**Elevated/ Low Levels, Not of clinical concern
***Elevated or Low Levels likely due to stress response associated with surgery
****Reticulocytes not considered significant unless animal anemic; RBCs normal
*****Neutrophils: elevated due to surgical inflammation
*.* Elevated/ Low Levels possibly due to Immunosuppression/implanted biomaterial/encapsulated cells/cell products
*_* Elevated levels possibly due to neutrophils being counted as eosinophils

**Table S5 (Continued). Complete blood counts for NHPs over the course of the TP-IAT to the omentum with PPCN study.**

(n = 2; 1 female (F), 1 male (M)); WBC = white blood cells; RBC = red blood cells; HGB = hemoglobin; HCT = hematocrit; MCV = mean corpuscular volume; MCH = mean corpuscular hemoglobin; MCHC = mean corpuscular hemoglobin concentration; PLT = platelets; MPV = mean platelet volume; RDW = red cell distribution width; NEUT = neutrophils; LYMPH = lymphocytes; MONO = monocytes; EOS = eosinophils; BASO = basophils; LUC = large unstained cells; RETIC =reticulocytes; MACRO = macrocytosis; HC-VAR = hemoglobin concentration variance; PLATCLMP = platelet clumps; HYPER = hyperchromia.

| Monkey: | **RH9144** (M) | | |
|---|---|---|---|
| Condition: | Post-Transplant | Post-Omentectomy | |
| Date: | 112 | 118 | |
| Day: | 10/11/2022 | 10/17/2022 | Units |
| WBC | 10.61 | 11.11 | x10^3/µL |
| RBC | 5.66 | 5.55 | x10^6/µL |
| HGB | 13.3 | 13.0 | g/dL |
| HCT | 41.9 | 42.6 | % |
| MCV | 74.0 | 76.9 | fL |
| MCH | 23.5 | 23.4 | pg |
| MCHC | 31.8 | 30.5 | g/dL |
| PLT | 476 | 501 | x10^3/µL |
| MPV | 7.5 | 8.4 | fL |
| RDW | 14.2 | 13.7 | % |
| %NEUT | 55.8* | 83***** | % |
| %LYMPH | 38.8 | 10.7*** | % |
| %MONO | 3.0 | 3.7 | % |
| %EOS | 0.6 | 1.0 | % |
| %BASO | 0.2 | 0.1 | % |
| %LUC | 1.6 | 1.5 | % |
| #NEUT | 5.92 | 9.22***** | x10^3/µL |
| #LYMPH | 4.12 | 1.19*** | x10^3/µL |
| #MONO | 0.32 | 0.41 | x10^3/µL |
| #EOS | 0.06 | 0.11 | x10^3/µL |
| #BASO | 0.02 | 0.01 | x10^3/µL |
| #LUC | 0.17 | 0.16 | x10^3/µL |
| %RETIC | 1.35 | 1.35 | % |
| #RETIC | 76.7**** | 75.1**** | x10^9/µL |

* Elevated/Low Levels, Comparable to baseline
**Elevated/ Low Levels, Not of clinical concern
***Elevated or Low Levels likely due to stress response associated with surgery
****Reticulocytes not considered significant unless animal anemic; RBCs normal
*****Neutrophils: elevated due to surgical inflammation
*.* Elevated/ Low Levels possibly due to Immunosuppression/implanted biomaterial/encapsulated cells/cell products
*_* Elevated levels possibly due to neutrophils being counted as eosinophils

**Table S6. Blood chemistry panel for NHPs over the course of the TP-IAT to the omentum with PPCN study.** (n = 2; 1 female (F), 1 male (M)); ALT = alanine transaminase; AST = aspartate aminotransferase; BUN = blood urea nitrogen; GGT = gamma-glutamyl transferase; I = inorganic.

| **Monkey:** | **RH9139 (F)** | | | | | | | | | |
|---|---|---|---|---|---|---|---|---|---|---|
| **Condition:** | Baseline | IVDTT Baseline | Pre-Transplant | Day of Transplant | Post-Transplant | | | | | |
| **Date:** | 8/4/2022 | 8/9/2022 | 8/15/2022 | 8/16/2022 | 8/19/2022 | 8/23/2022 | 8/30/2022 | 9/6/2022 | 9/13/2022 | |
| **Day:** | -12 | -7 | -1 | 0 | 3 | 7 | 14 | 21 | 28 | **Units** |
| Albumin | 4.198 | 4.374 | 4.047 | 3.516 | 3.361 | 3.788 | 3.806 | 3.856 | 3.724 | g/dL |
| Alkaline Phosphatase | 126 | 135 | 123 | 115 | 126 | 151 | 140 | 157 | 141 | U/L |
| ALT | 20 | 37***** | 32***** | 51***** | 44***** | 93***** | 72***** | 75***** | 57***** | U/L |
| AST | 19 | 25 | 22 | 88*** | 24 | 36 | 35 | 37 | 32 | U/L |
| Urea Nitrogen (BUN) | 22 | 11 | 17 | 19 | 13 | 15 | 15 | 15 | 13 | mg/dL |
| Calcium | 9.88 | 9.17 | 9.14 | 8.39 | 9.06 | 9.14 | 9.73 | 9.23 | 9.21 | mg/dL |
| Creatinine | 0.7784 | 0.7131 | 0.663 | 0.7321 | 0.7134 | 0.6466 | 0.6045 | 0.636 | 0.5912 | mg/dL |
| GGT | 38 | 39 | 34 | 33 | 31 | 37 | 40 | 46 | 41 | U/L |
| Glucose | 102 | 78 | 89 | 55 | 136 | 96 | 98 | 175 | 123 | mg/dL |
| I. Phosphorous | 3.6 | 5.3 | 5.3 | 3.5 | 4.4 | 4.7 | 3.9 | 4.3 | 4.7 | mg/dL |
| Total Bilirubin | 0.13 | 0.15 | 0.14 | 0.16 | 0.12 | 0.08 | 0.12 | 0.16 | 0.16 | mg/dL |
| Total Protein | 6.087 | 6.268 | 5.987 | 5.047*** | 5.493*** | 6.149 | 5.625 | 5.911 | 5.621 | g/dL |
| Sodium | 144 | 142 | 144 | 145 | 143 | 153 | 144 | 141 | 144 | mmol/L |
| Potassium | 3.8 | 3.1* | 3.4* | 3.0** | 4.1 | 3.9 | 3.5 | 3.5 | 3.4 | mmol/L |
| Chloride | 105 | 103* | 104* | 108 | 103* | 108 | 105* | 103* | 105* | mmol/L |
| Globulin | 1.889 | 1.894* | 1.94* | 1.531** | 2.132** | 2.361** | 1.819* | 2.055* | 1.897* | |
| Alb/Glob Ratio | 2 | 2 | 2 | 2 | 2 | 2 | 2 | 2 | 2 | |

* Elevated/Low Levels, Comparable to baseline
** Elevated/Low Levels, Not of clinical concern
***Elevated/Low Levels likely due to stress response associated with surgery
****Muscle component, indicative of possible damage to muscle - may be related to anesthesia injections or damage to muscle during surgery
*****Elevated ALT levels are considered of clinical concern only if the levels are more than 100 IU/L.
******Elevated Globulin levels than Albumin levels are indicative of inflammation, possibly due to implanted microcapsules, encapsulated cells or cell products
*******Elevated levels, Not of clinical concern; Possibly mild inflammation of liver but transient
*.* Elevated/ Low Levels likely due to Immunosuppression/implanted biomaterial/encapsulated cells/cell products
*_* Elevated due to diabetic condition of the animal

**Table S6 (Continued). Blood chemistry panel for NHPs over the course of the TP-IAT to the omentum with PPCN study.**

(n = 2; 1 female (F), 1 male (M)); ALT = alanine transaminase; AST = aspartate aminotransferase; BUN = blood urea nitrogen; GGT = gamma-glutamyl transferase; I = inorganic.

| **Monkey:** | **RH9139 (F)** | | | | | | | | |
|---|---|---|---|---|---|---|---|---|---|
| **Condition:** | IVDTT Post-Transplant | Glucagon Post-Transplant | Post-Transplant | | IVDTT Post-Transplant | Post-Transplant | | | |
| **Date:** | 9/15/2022 | 9/20/2022 | 9/27/2022 | 10/4/2022 | 10/10/2022 | 10/18/2022 | 10/25/2022 | 11/1/2022 | |
| **Day:** | 30 | 35 | 42 | 49 | 55 | 63 | 70 | 77 | **Units** |
| Albumin | 3.764 | 3.342 | 3.351 | 3.621 | 3.895 | 3.559 | 3.434 | 3.395 | g/dL |
| Alkaline Phosphatase | 138 | 122 | 130 | 113 | 117 | 126 | 104 | 103 | U/L |
| ALT | 81***** | 36***** | 25***** | 36***** | 39***** | 45***** | 30***** | 27***** | U/L |
| AST | 43 | 25 | 31 | 27 | 37 | 29 | 28 | 23 | U/L |
| Urea Nitrogen (BUN) | 13 | 20 | 23 | 20 | 18 | 21 | 25 | 25 | mg/dL |
| Calcium | 9.38 | 8.52 | 9.15 | 8.7 | 8.62 | 9.03 | 9.16 | 8.36 | mg/dL |
| Creatinine | 0.5242 | 0.6451 | 0.6927 | 0.5705 | 0.5513 | 0.526 | 0.5724 | 0.5328 | mg/dL |
| GGT | 42 | 33 | 32 | 33 | 34 | 37 | 32 | 32 | U/L |
| Glucose | 142 | 80 | 118 | 113 | 153 | 206 | 121 | 110 | mg/dL |
| I. Phosphorous | 5.8 | 4.9 | 3.9 | 5.2 | 5.3 | 3.8 | 4 | 3.7 | mg/dL |
| Total Bilirubin | 0.14 | 0.15 | 0.11 | 0.11 | 0.16 | 0.1 | 0.14 | 0.12 | mg/dL |
| Total Protein | 5.652 | 5.157* | 5.217* | 5.529* | 5.728 | 6.739 | 5.16* | 5.201* | g/dL |
| Sodium | 143 | 142 | 157 | 144 | 131* | 140 | 142 | 142 | mmol/L |
| Potassium | 3.7 | 3.1* | 4 | 3.2* | 3.4 | 4 | 3.6 | 3.3 | mmol/L |
| Chloride | 106* | 103* | 112 | 103* | 97* | 105* | 105* | 106* | mmol/L |
| Globulin | 1.888* | 1.815* | 1.866* | 1.908* | 1.833* | - | 1.726* | 1.806* | |
| Alb/Glob Ratio | 2 | 2 | 2 | 2 | 2 | - | 2 | 2 | |

* Elevated/Low Levels, Comparable to baseline
** Elevated/Low Levels, Not of clinical concern
***Elevated/Low Levels likely due to stress response associated with surgery
****Muscle component, indicative of possible damage to muscle - may be related to anesthesia injections or damage to muscle during surgery
*****Elevated ALT levels are considered of clinical concern only if the levels are more than 100 IU/L.
******Elevated Globulin levels than Albumin levels are indicative of inflammation, possibly due to implanted microcapsules, encapsulated cells or cell products
*******Elevated levels, Not of clinical concern; Possibly mild inflammation of liver but transient
*.* Elevated/ Low Levels likely due to Immunosuppression/implanted biomaterial/encapsulated cells/cell products
*_* Elevated due to diabetic condition of the animal

**Table S6 (Continued). Blood chemistry panel for NHPs over the course of the TP-IAT to the omentum with PPCN study.**

(n = 2; 1 female (F), 1 male (M)); ALT = alanine transaminase; AST = aspartate aminotransferase; BUN = blood urea nitrogen; GGT = gamma-glutamyl transferase; I = inorganic.

| **Monkey:** | **RH9139 (F)** | | | | | | | |
|---|---|---|---|---|---|---|---|---|
| **Condition:** | Post-Transplant | | | IVDTT Post-Transplant | Survival Omentectomy Post-Transplant | IVDTT Post - Omentectomy | Termination & Necropsy | |
| **Date:** | 11/7/2022 | 11/15/2022 | 11/22/2022 | 11/29/2022 | 12/2/2022 | 12/5/2022 | 12/8/2022 | |
| **Day:** | 83 | 91 | 98 | 105 | 108 | 111 | 114 | **Units** |
| Albumin | 3.486 | 3.403 | 3.289 | 3.306 | 3.359 | 3.201 | 3.233 | g/dL |
| Alkaline Phosphatase | 111 | 101 | 96 | 116 | 107 | 166 | 117 | U/L |
| ALT | 32***** | 36***** | 29***** | 31***** | 39***** | 43***** | 43***** | U/L |
| AST | 24 | 28 | 23 | 28 | 29 | 30 | 40 | U/L |
| Urea Nitrogen (BUN) | 12 | 17 | 20 | 19 | 14 | 26 | 14 | mg/dL |
| Calcium | 8.38 | 8.28 | 8.26 | 8.22 | 8.16 | 8.23 | 8.37 | mg/dL |
| Creatinine | 0.5241 | 0.4902 | 0.4688 | 0.5528 | 0.4947 | 0.6162 | 0.4789 | mg/dL |
| GGT | 33 | 32 | 30 | 34 | 34 | 34 | 35 | U/L |
| Glucose | 73 | 68 | 142 | 149 | 56 | 199 | 89 | mg/dL |
| I. Phosphorous | 5.8 | 5.4 | 4.3 | 4.5 | 5.6 | 6 | 5.1 | mg/dL |
| Total Bilirubin | 0.16 | 0.14 | 0.13 | 0.16 | 0.14 | 0.24 | 0.14 | mg/dL |
| Total Protein | 5.559* | 5.2* | 5.008* | 4.94* | 5.399* | 5.574* | 5.426* | g/dL |
| Sodium | 143 | 143 | 139* | 140 | 141 | 137* | 145 | mmol/L |
| Potassium | 3.3 | 3.1* | 3.3 | 3.2* | 3.1* | 3.6 | 3.0** | mmol/L |
| Chloride | 104* | 104* | 103* | 104* | 102* | 94* | 102* | mmol/L |
| Globulin | 2.073* | 1.797* | 1.719* | 1.634** | 2.04* | 2.373** | 2.193** | |
| Alb/Glob Ratio | 2 | 2 | 2 | 2 | 2 | 1 | 1 | |

* Elevated/Low Levels, Comparable to baseline
** Elevated/Low Levels, Not of clinical concern
***Elevated/Low Levels likely due to stress response associated with surgery
****Muscle component, indicative of possible damage to muscle - may be related to anesthesia injections or damage to muscle during surgery
*****Elevated ALT levels are considered of clinical concern only if the levels are more than 100 IU/L.
******Elevated Globulin levels than Albumin levels are indicative of inflammation, possibly due to implanted microcapsules, encapsulated cells or cell products
*******Elevated levels, Not of clinical concern; Possibly mild inflammation of liver but transient
*.* Elevated/ Low Levels likely due to Immunosuppression/implanted biomaterial/encapsulated cells/cell products
*_* Elevated due to diabetic condition of the animal

**Table S6 (Continued). Blood chemistry panel for NHPs over the course of the TP-IAT to the omentum with PPCN study.**

(n = 2; 1 female (F), 1 male (M)); ALT = alanine transaminase; AST = aspartate aminotransferase; BUN = blood urea nitrogen; GGT = gamma-glutamyl transferase; I = inorganic.

| **Monkey:** | **RH9144 (M)** | | | | | | | | | |
|---|---|---|---|---|---|---|---|---|---|---|
| **Condition:** | Pre-Transplant | | Post-Transplant | | | | | | | |
| **Date:** | -6 | 0 | 7 | 14 | 21 | 28 | 30 | 42 | 63 | |
| **Day:** | 6/15/2022 | 6/20/2022 | 6/28/2022 | 7/5/2022 | 7/12/2022 | 7/19/2022 | 7/21/2022 | 8/2/2022 | 8/23/2022 | **Units** |
| Albumin | 4.771 | 4.426 | 3.693 | 3.912 | 4.096 | 4.367 | 4.283 | 4.433 | 4.616 | g/dL |
| Alkaline Phosphatase | 391 | 338 | 336 | 327 | 365 | 439 | 426 | 419 | 506 | U/L |
| ALT | 32***** | 31***** | 27***** | 29***** | 42***** | 50***** | 59***** | 36***** | 39***** | U/L |
| AST | 47 | 24 | 23 | 28 | 23 | 31 | 39 | 26 | 25 | U/L |
| Urea Nitrogen (BUN) | 23 | 16 | 16 | 15 | 16 | 14 | 13 | 18 | 21 | mg/dL |
| Calcium | 10.1 | 9.9 | 10.02 | 10.48 | 10.37 | 10.02 | 9.78 | 10.03 | 10.65 | mg/dL |
| Creatinine | 0.7818 | 0.764 | 0.7105 | 0.6511 | 0.6658 | 0.7103 | 0.6814 | 0.7567 | 0.8618 | mg/dL |
| GGT | 75 | 59 | 45 | 53 | 66 | 73 | 68 | 77 | 87 | U/L |
| Glucose | 116 | 114 | 302*_* | 62 | 147 | 56 | 103 | 106 | 150 | mg/dL |
| I. Phosphorous | 3.4 | 6.4 | 5 | 5.5 | 6.5 | 5.3 | 6.6 | 6 | 6.1 | mg/dL |
| Total Bilirubin | 0.24 | 0.22 | 0.14 | 0.11 | 0.14 | 0.13 | 0.18 | 0.14 | 0.12 | mg/dL |
| Total Protein | 6.814 | 6.234 | 6.015 | 6.64 | 6.607 | 6.529 | 6.44 | 6.656 | 6.717 | g/dL |
| Sodium | 144 | 144 | 143 | 143 | 144 | 143 | 145 | 143 | 151 | mmol/L |
| Potassium | 3.9 | 4.5 | 4.4 | 4.3 | 4 | 3.3 | 3.8 | 3.7 | 4 | mmol/L |
| Chloride | 103 | 103* | 100* | 102* | 102* | 101* | 102* | 101* | 108 | mmol/L |
| Globulin | 2.043 | 1.808* | 2.322* | 2.728 | 2.511* | 2.162* | 2.157* | 2.223* | 2.101* | |
| Alb/Glob Ratio | 2 | 2 | 2 | 1 | 1 | 2 | 2 | 2 | 2 | |

* Elevated/Low Levels, Comparable to baseline

** Elevated/Low Levels, Not of clinical concern

***Elevated/Low Levels likely due to stress response associated with surgery

****Muscle component, indicative of possible damage to muscle - may be related to anesthesia injections or damage to muscle during surgery

*****Elevated ALT levels are considered of clinical concern only if the levels are more than 100 IU/L.

******Elevated Globulin levels than Albumin levels are indicative of inflammation, possibly due to implanted microcapsules, encapsulated cells or cell products

*******Elevated levels, Not of clinical concern; Possibly mild inflammation of liver but transient

*.* Elevated/ Low Levels likely due to Immunosuppression/implanted biomaterial/encapsulated cells/cell products

*_* Elevated due to diabetic condition of the animal

**Table S6 (Continued). Blood chemistry panel for NHPs over the course of the TP-IAT to the omentum with PPCN study.**
(n = 2; 1 female (F), 1 male (M)); ALT = alanine transaminase; AST = aspartate aminotransferase; BUN = blood urea nitrogen; GGT = gamma-glutamyl transferase; I = inorganic.

| **Monkey:** | **RH9144 (M)** | | | | | | | | |
|---|---|---|---|---|---|---|---|---|---|
| **Condition:** | Post-Transplant | | | | | | | Post-Omentectomy | |
| **Date:** | 70 | 77 | 84 | 91 | 98 | 105 | 112 | 118 | |
| **Day:** | 8/30/2022 | 9/6/2022 | 9/13/2022 | 9/20/2022 | 9/27/2022 | 10/4/2022 | 10/11/2022 | 10/17/2022 | **Units** |
| Albumin | 4.403 | 4.289 | 4.302 | 4.242 | 4.57 | 4.536 | 4.527 | 4.686 | g/dL |
| Alkaline Phosphatase | 471 | 501 | 488 | 484 | 477 | 487 | 479 | 455 | U/L |
| ALT | 40***** | 38***** | 31***** | 30***** | 28***** | 29***** | 31***** | 59***** | U/L |
| AST | 31 | 23 | 21 | 25 | 31 | 23 | 29 | 42 | U/L |
| Urea Nitrogen (BUN) | 20 | 19 | 19 | 21 | 21 | 20 | 12 | 28** | mg/dL |
| Calcium | 10.36 | 10.25 | 10.21 | 10.01 | 10.3 | 10.32 | 10.16 | 10.11 | mg/dL |
| Creatinine | 0.7737 | 0.7748 | 0.8474 | 0.8133 | 0.8157 | 0.8883 | 0.8628 | 1.0647 | mg/dL |
| GGT | 88 | 91 | 86 | 82 | 90 | 88 | 84 | 79 | U/L |
| Glucose | 110 | 109 | 229 | 70 | 126 | 132 | 112 | 237*_* | mg/dL |
| I. Phosphorous | 5.8 | 5.6 | 6.6 | 5.9 | 6.7 | 6.3 | 6.5 | 8 | mg/dL |
| Total Bilirubin | 0.17 | 0.18 | 0.2 | 0.19 | 0.17 | 0.18 | 0.2 | 0.27 | mg/dL |
| Total Protein | 6.305 | 6.45 | 6.546 | 6.307 | 6.685 | 6.698 | 6.5 | 7.228 | g/dL |
| Sodium | 144 | 141 | 147 | 143 | 155 | 144 | 143 | 144 | mmol/L |
| Potassium | 3.8 | 4.1 | 4.4 | 3.5 | 3.8 | 3.8 | 3.7 | 4.7 | mmol/L |
| Chloride | 103* | 102* | 106* | 102* | 109 | 101* | 102* | 96* | mmol/L |
| Globulin | 1.902* | 2.161* | 2.244* | 2.065* | 2.115* | 2.162* | 2.062* | 2.542 | |
| Alb/Glob Ratio | 2 | 2 | 2 | 2 | 2 | 2 | 2 | 2 | |

* Elevated/Low Levels, Comparable to baseline
** Elevated/Low Levels, Not of clinical concern
***Elevated/Low Levels likely due to stress response associated with surgery
****Muscle component, indicative of possible damage to muscle - may be related to anesthesia injections or damage to muscle during surgery
*****Elevated ALT levels are considered of clinical concern only if the levels are more than 100 IU/L.
******Elevated Globulin levels than Albumin levels are indicative of inflammation, possibly due to implanted microcapsules, encapsulated cells or cell products
*******Elevated levels, Not of clinical concern; Possibly mild inflammation of liver but transient
*.* Elevated/ Low Levels likely due to Immunosuppression/implanted biomaterial/encapsulated cells/cell products
*_* Elevated due to diabetic condition of the animal

**Table S7. Urinalysis for NHPs over the course of the TP-IAT to the omentum with PPCN study.** (n = 2; 1 female (F), 1 male (M)); RBC = red blood cells; WBC = white blood cells.

| | **Monkey:** | **RH9139 (F)** | | | | | | | | |
|---|---|---|---|---|---|---|---|---|---|---|
| | **Condition:** | IVDTT Baseline | Pre-Transplant | Day of Transplant | Post-Transplant | | | | | Glucagon Post-Transplant |
| | **Date:** | 8/9/2022 | 8/15/2022 | 8/16/2022 | 8/19/2022 | 8/23/2022 | 8/30/2022 | 9/6/2022 | 9/13/2022 | 9/20/2022 |
| | **Day:** | -7 | -1 | 0 | 3 | 7 | 14 | 21 | 28 | 35 |
| ***Physical*** | **Appearance** | Clear | Clear | Clear | Clear | Clear | Clear | Clear | Hazy | Hazy |
| | **Specific Gravity** | 1.002 | 1.009 | 1.012 | 1.008 | 1.028 | 1.005 | 1.003 | 1.003 | 1.01 |
| | **Color** | Colorless | Light Yellow | Light Yellow | Colorless | Yellow | Colorless | Colorless | Colorless | Light Yellow |
| ***Dipstick Evaluation*** | **Leukocytes** | Negative | Negative | Negative | Negative | Negative | Negative | Negative | + | Negative |
| | **Nitrite** | Negative | Negative | Negative | Negative | Negative | Negative | Negative | Negative | Negative |
| | **pH** | 7 | 8 | 6 | 7 | 8 | 8 | 7 | 8 | 8 |
| | **Protein** | Trace | Negative | Trace | Trace | Trace | Negative | Negative | Trace | Trace |
| | **Glucose** | Normal | Normal | Normal | 100mg/dl | Normal | Normal | 250 mg/dl | Normal | Normal |
| | **Ketones** | Negative | Negative | Negative | Negative | Negative | Negative | Negative | Negative | Negative |
| | **Urobilinogen** | Normal | Normal | Normal | Normal | Normal | Normal | Normal | Normal | Normal |
| | **Bilirubin** | Negative | Negative | Negative | Negative | Negative | Negative | Negative | Negative | Negative |
| | **Blood** | Negative | Trace | 50 | Negative | Negative | Negative | Negative | Negative | 250 |

| | **Monkey:** | **RH9139 (F)** | | | | | | | | |
|---|---|---|---|---|---|---|---|---|---|---|
| | **Condition:** | Post-Transplant | | IVDTT Post-Transplant | Post-Transplant | | | | | |
| | **Date:** | 9/27/2022 | 10/4/2022 | 10/10/2022 | 10/18/2022 | 10/25/2022 | 11/1/2022 | 11/7/2022 | 11/15/2022 | 11/22/2022 |
| | **Day:** | 42 | 49 | 55 | 63 | 70 | 77 | 83 | 91 | 98 |
| ***Physical*** | **Appearance** | Clear | Clear | Clear | Clear | Clear | Clear | Clear | Clear | Clear |
| | **Specific Gravity** | 1.01 | 1.003 | 1.003 | 1.007 | 1.004 | 1.005 | 1.004 | 1.012 | 1.003 |
| | **Color** | Light Yellow | Colorless | Colorless | Colorless | Colorless | Colorless | Light Yellow | Light Yellow | Colorless |
| ***Dipstick Evaluation*** | **Leukocytes** | Negative | Negative | Negative | Negative | Negative | Negative | Negative | + | Negative |
| | **Nitrite** | Negative | Negative | Negative | Negative | Negative | Negative | Negative | Negative | Negative |
| | **pH** | 8 | 8 | 5 | 7 | 7 | 5 | 7 | 8 | 7 |
| | **Protein** | Trace | Trace | Negative | Negative | Trace | Trace | Trace | Trace | Trace |
| | **Glucose** | Normal | Normal | 500 mg/dl | 500 mg/dl | 100mg/dl | Normal | Normal | Normal | 50 mg/dl |
| | **Ketones** | Negative | Negative | Negative | Negative | Negative | Negative | Negative | Negative | Negative |
| | **Urobilinogen** | Normal | Normal | Normal | Normal | Normal | Normal | Normal | Normal | Normal |
| | **Bilirubin** | Negative | Negative | Negative | Negative | Negative | Negative | Negative | Negative | Negative |
| | **Blood** | Negative | Negative | Negative | Negative | Negative | Negative | Negative | Trace | Negative |

**Table S7 (Continued). Urinalysis for NHPs over the course of the TP-IAT to the omentum with PPCN study.** (n = 2; 1 female (F), 1 male (M)); RBC = red blood cells; WBC = white blood cells.

| | Monkey: | RH9139 (F) | | | |
|---|---|---|---|---|---|
| | **Condition:** | IVDTT Post-Transplant | Survival Omentectomy Post-Transplant | IVDTT Post - Omentectomy | Termination & Necropsy |
| | **Date:** | 11/29/2022 | 12/2/2022 | 12/5/2022 | 12/8/2022 |
| | **Day:** | 105 | 108 | 111 | 114 |
| *Physical* | **Appearance** | Clear | Clear | Clear | Hazy |
| | **Specific Gravity** | 1.012 | 1.023 | 1.025 | 1.014 |
| | **Color** | Light Yellow | Yellow | Light Yellow | Light Yellow |
| *Dipstick Evaluation* | **Leukocytes** | Negative | Negative | Negative | Negative |
| | **Nitrite** | Negative | Negative | Negative | Negative |
| | **pH** | 8 | 9 | 5 | 8 |
| | **Protein** | Trace | 100 | Negative | Trace |
| | **Glucose** | 250mg/dl | Normal | 1000mg/dl | Normal |
| | **Ketones** | Negative | Negative | ++ (Moderate) | Negative |
| | **Urobilinogen** | Normal | Normal | Normal | Normal |
| | **Bilirubin** | Negative | Negative | Negative | Negative |
| | **Blood** | Trace | Negative | Negative | Negative |

**Table S7 (Continued). Urinalysis for NHPs over the course of the TP-IAT to the omentum with PPCN study.** (n = 2; 1 female (F), 1 male (M)); RBC = red blood cells; WBC = white blood cells.

| | Monkey: | RH9144 (M) | | | | | | | | |
|---|---|---|---|---|---|---|---|---|---|---|
| | Condition: | Pre-Transplant | | Post-Transplant | | | | | | |
| | Date: | 0 | 7 | 14 | 21 | 28 | 42 | 49 | 63 | 70 |
| | Day: | 6/20/2022 | 6/28/2022 | 7/5/2022 | 7/12/2022 | 7/19/2022 | 8/2/2022 | 8/9/2022 | 8/23/2022 | 8/30/2022 |
| *Physical* | Appearance | Clear | Clear | Clear | Clear | Clear | Clear | Clear | Clear | Clear |
| | Specific Gravity | 1.004 | 1.016 | 1.02 | 1.003 | 1.007 | 1.008 | 1.004 | 1.023 | 1.011 |
| | Color | Light Yellow | Colorless | Yellow | Light Yellow | Light Yellow | Yellow | Colorless | Light Yellow | Light Yellow |
| *Dipstick Evaluation* | Leukocytes | Negative | Negative | Trace | Negative | Negative | Negative | Negative | Negative | Negative |
| | Nitrite | Negative | Negative | Negative | Negative | Negative | Negative | Negative | Negative | Negative |
| | pH | 7 | 7 | 8 | 7 | 8 | 8 | 7 | 8 | 8 |
| | Protein | Trace | Trace | Trace | Trace | Trace | Trace | Trace | Trace | Trace |
| | Glucose | Normal | 500mg/dl | Normal | 50mg/dl | Normal | Normal | Normal | Normal | Normal |
| | Ketones | Negative | Negative | Negative | Negative | Negative | Negative | Negative | Negative | Negative |
| | Urobilinogen | Normal | Normal | Normal | Normal | Normal | Normal | Normal | Normal | Normal |
| | Bilirubin | Negative | Negative | Negative | Negative | Negative | Negative | Negative | Negative | Negative |
| | Blood | 250Ery/µl | Trace | 50Ery/µl | Negative | Negative | Negative | Negative | Negative | 50 |

| | Monkey: | RH9144 (M) | | | | | | |
|---|---|---|---|---|---|---|---|---|
| | Condition: | Post-Transplant | | | | | | Post-Omentectomy |
| | Date: | 77 | 84 | 91 | 98 | 105 | 112 | 118 |
| | Day: | 9/6/2022 | 9/13/2022 | 9/20/2022 | 9/27/2022 | 10/4/2022 | 10/11/2022 | 10/17/2022 |
| *Physical* | Appearance | Clear | Clear | Hazy | Clear | Clear | Clear | Clear |
| | Specific Gravity | 1.003 | 1.006 | 1.008 | 1.027 | 1.004 | 1.02 | 1.025 |
| | Color | Colorless | Light Yellow | Light Yellow | Yellow | Light Yellow | Yellow | Light Yellow |
| *Dipstick Evaluation* | Leukocytes | Negative | Negative | Negative | Negative | Negative | Negative | Negative |
| | Nitrite | Negative | Negative | Negative | Negative | Negative | Negative | Negative |
| | pH | 7 | 7 | 7 | 9 | 8 | 9 | 5 |
| | Protein | Trace | Trace | Trace | 30 | Negative | 30 | Trace |
| | Glucose | Normal | 500mg/dl | Normal | Normal | Normal | Normal | 1000mg/dl |
| | Ketones | Negative | Negative | Negative | Negative | Negative | Negative | ++ |
| | Urobilinogen | Normal | Normal | Normal | Normal | Normal | Normal | Normal |
| | Bilirubin | Negative | Negative | Negative | Negative | Negative | Negative | Negative |
| | Blood | Negative | Negative | Negative | Negative | Negative | Negative | Negative |
